# Altering translation allows *E. coli* to overcome chemically stabilized G-quadruplexes

**DOI:** 10.1101/2024.08.12.607615

**Authors:** Rachel R Cueny, Andrew F Voter, Aidan M McKenzie, Marcel Morgenstern, Kevin S Myers, Michael M Place, Jason M. Peters, Joshua J Coon, James L Keck

## Abstract

G-quadruplex (G4) structures can form in guanine-rich DNA or RNA and have been found to modulate cellular processes including replication, transcription, and translation. Many studies on the cellular roles of G4s have focused on eukaryotic systems, with far fewer probing bacterial G4s. Using a chemical-genetic approach, we identified genes in *Escherichia coli* that are important for growth in G4-stabilizing conditions. Reducing levels of elongation factor Tu or slowing translation elongation with chloramphenicol suppress the effects of G4 stabilization. In contrast, reducing expression of certain translation termination or ribosome recycling proteins is detrimental to growth in G4-stabilizing conditions. Proteomic and transcriptomic analyses demonstrate that ribosome assembly factors and other proteins involved in translation are less abundant in G4-stabilizing conditions. Our integrated systems approach allowed us to propose a model for how RNA G4s can present barriers to *E. coli* growth and that reducing the rate of translation can compensate for G4-related stress.

## Introduction

G-quadruplexes (G4s) are nucleic acid structures that can fold in guanine-rich stretches of DNA or RNA (1). The structures are comprised of stacked G-tetrads (four guanines hydrogen bonded to one another) surrounding a core of monovalent cations. Interest in possible biological roles for G4s emerged upon discovering that guanine-rich telomeres in humans and other eukaryotes could fold into G4 structures (2–6).

G4s have been found to modulate replication, transcription, and translation processes. For example, G4s can act as barriers to DNA replication in eukaryotes, an effect that is exacerbated upon depletion of accessory DNA helicases (7–10). DNA G4s can also impact transcription in a strand- and location-dependent manner (11–14). Finally, G4s found in mRNA transcripts can regulate translation efficiency, with G4s in the open reading frame decreasing translation and G4s in the untranslated regions having various effects on translation in eukaryotes and prokaryotes (1,11,15–18). Several proteins that can bind and unwind G4s have been shown to govern G4 homeostasis in cells (1,19–26).

G-rich repeats in neurodegenerative disorders such as amyotrophic lateral sclerosis, G4 sequences in oncogenes, and G4s at telomeres have motivated studies to understand G4s in eukaryotes (2,12,13,17,27–29). Despite advances in understanding the roles of G4s in eukaryotic systems, parallel studies in bacteria have been far more limited. One notable exception is from the pathogenic bacteria *Neisseria gonorrhoeae*, in which a G4 has been found to be essential for antigenic variation (30,31). Beyond this example, most studies investigating G4s in *Escherichia coli* and other bacteria have focused on identifying potential quadruplex forming sequences or have investigated the effects of non-native G4 sequences in plasmids on growth or gene expression (1,11,32,33). Other approaches have taken a candidate-based approach to examine roles of selected DNA helicases or DNA repair proteins in G4 processing *in vitro* or *in vivo* (1,23,26).

To better understand the challenges presented by G4s that exist in bacteria, we carried out tandem chemical-genetic screens to identify genes that are important for *E. coli* growth in the presence of the G4-stabilizing compound N-methyl-mesoporphyrin IX (NMM). A transposon-sequencing (Tn-seq) screen showed that two genes, *tufA* and *tufB*, contained a disproportionately high amount of transposon insertions in G4-stabilizing conditions, indicating that disruption of either gene was strongly selected. *tufA* and *tufB* encode for the same protein, elongation factor (EF) Tu, which escorts charged tRNAs to the ribosome during translation and impacts the rate of translation and cell growth (34–36). Disruption of either *tufA* or *tufB* reduced cellular EF-Tu levels, suggesting that slowing translation could counter the negative effects of G4 stabilization. In concurrence with this idea, low doses of the bacterial translation inhibitor chloramphenicol improved the growth of *E. coli* in the presence of NMM. Next, a CRISPR interference (CRISPRi) screen identified the importance of a ribosome release factor (RF1) and an elongation factor involved in ribosome recycling (EF-G) for growth in NMM, further supporting a model in which translation factors are linked to overcoming stabilized G4s *E. coli*. Proteomic and transcriptomic analysis of cells grown ± NMM identified several proteins/transcripts that were differentially expressed in the presence of NMM, with an overrepresentation of downregulated ribosome assembly/biogenesis terms. Our observations collectively point to a model in which RNA G4s influence translation and alterations to translation processes can allow *E. coli* to overcome the detrimental impacts of stabilized G4s. These results provide new insight into RNA G4 biology both in bacteria and in RNA G4 homeostasis, and they suggest that transiently folding RNA G4s could be an unexpected target for modulating bacterial cell viability.

## Results

### Disruption of TolC efflux is necessary for efficient NMM retention

To better understand the effects of stabilized G4s in *E. coli*, Tn-seq was used to identify genes that alter cell growth in the presence of the G4 stabilizer, NMM. NMM stacks atop G4 structures, leading to stabilized G4s that can act as barriers to cellular processes such as DNA replication, transcription, or translation (**Figure 1A**) (1,26). We predicted that strains with transposon insertions in genes that help withstand G4-stabilized conditions would be selected against when grown in the presence of NMM while strains with insertions in genes that impair growth in G4-stabilized conditions would be positively selected (**Figure 1B**). An initial screen using a transposon insertion library made in *E. coli* MG1655 (∼200,000 individual strains) revealed that insertions in *tolC*, which encodes a component of two major efflux systems (37), sensitized cells to NMM (Figure S1). These data suggested that TolC efflux can export NMM from the *E. coli* cytoplasm, which agreed with a previous finding that loss of *tolC* in *E. coli* causes cytoplasmic accumulation of porphyrins (38,39).

**Figure 1.**
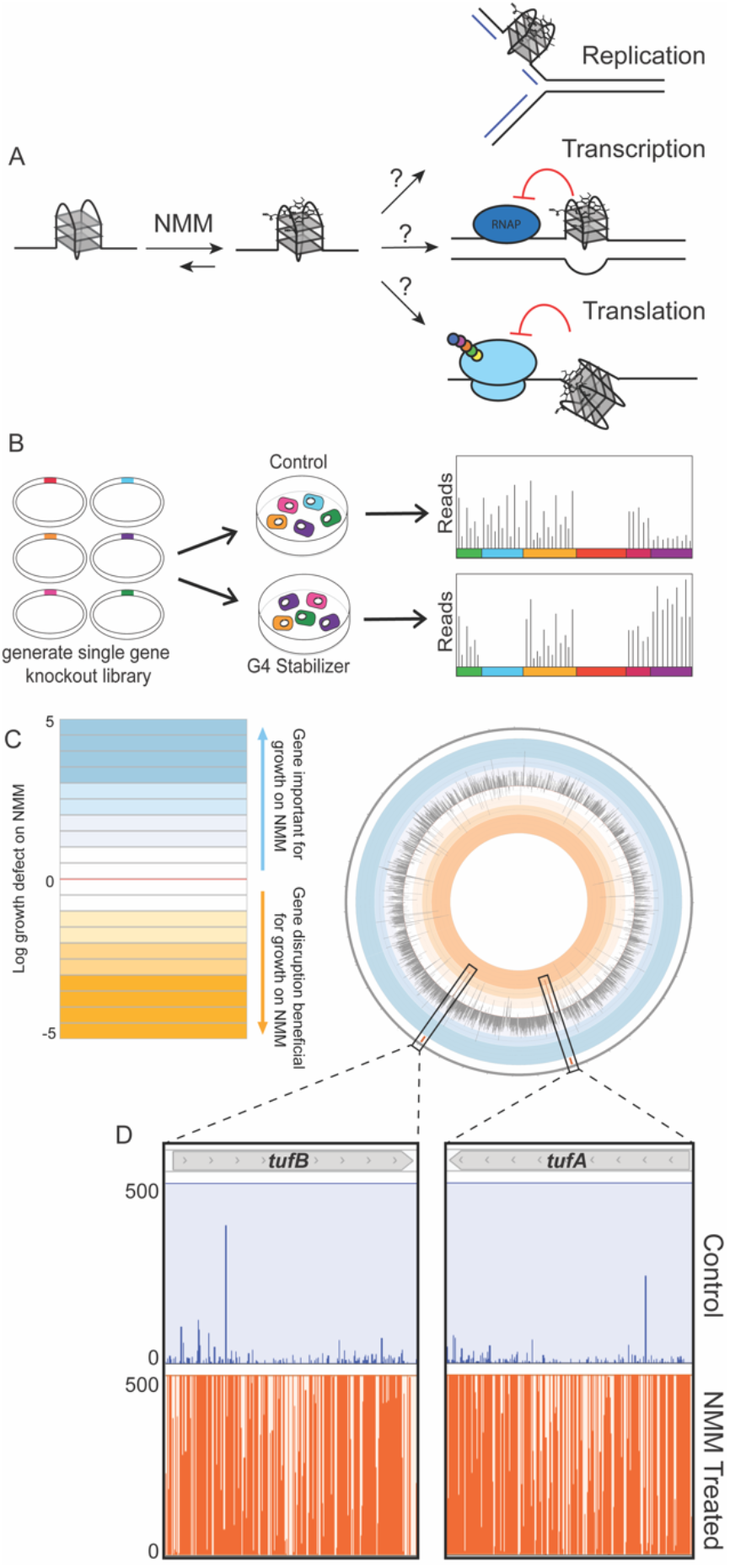
Transposon sequencing reveals genes important for overcoming stabilized G-quadruplexes. **(A)**. Cartoon depicting the potential consequences of chemically stabilized G-quadruplexes using NMM. **(B)**. Depiction of Tn-seq experiment. A single gene knockout library is generated and grown in control or G4-stabilizing conditions, and then sequenced to determine where transposon insertions are tolerated in each condition. **(C)**. Circos plot of the log_10_(NMM weighted reads/control weighted reads). **(D)**. Zoom-in on insertions across two genes of interest, *tufA* and *tufB*, in control and NMM-treated conditions. Each line is an insertion with the height reflecting the number of reads within a given insertion.

### Disruption of translation genes improves relative fitness in G4-stabilizing conditions

To facilitate cytoplasmic NMM retention in cells, a Δ*tolC E. coli* strain was used in a second set of Tn-seq experiments. Three biological replicate libraries of Δ*tolC E. coli* transposon insertion mutants (∼500,000 total) were generated and grown with or without NMM to obtain ∼1.5 million colonies selected from each growth condition. Transposon position identification through sequencing mapped the position and abundance of insertions within the populations. Normalized weighted reads ratios were determined for each gene based on transposon insertion tolerance in control and NMM-treated growth conditions and gene length (40). Positive or negative log_10_(normalized weighted reads ratio) (log_10_(n.w.r.r.)) values corresponded to genes in which transposon insertion is selected against or for, respectively, in NMM-treated conditions compared to control growth conditions. Of the 4,312 genes assessed, 3,900 allowed transposon insertions and 2,210 genes contained an average of five unique hits in the control library, showing strong coverage of transposon insertions across the libraries.

Comparison of transposon insertion tolerance in control and NMM-treated conditions revealed the impact of gene disruption in each growth condition (**Figure 1C**). Measured log_10_(n.w.r.r.) values ranged from 3.70 to -4.78 with 577 genes having values ≥ 1.5 or ≤ -1.5, reflecting a wide-ranging impact of NMM on transposon selection. To systematically identify pathways that are most impacted by transposon insertions by stabilized G4s, gene ontology (GO) term analyses were carried out using genes with log_10_(n.w.r.r.) ≥ 1.5 or ≤ -1.5 (Tables S1 & S2). The analyses indicated that disruptions in a variety of pathways impact growth, positively or negatively, in the presence of NMM. Consistent with previous work, *recA* and *rep*, which encode proteins involved in homologous recombination and accessory helicase activity, respectively, were conditionally important genes with log_10_(n.w.r.r.) values of 1.98 and 1.19 (26). In contrast, insertions in genes with GO terms related to ribosome assembly or regulation of translation were strongly enriched in cells grown in G4-stabilizing conditions, consistent with perturbation of translation aiding growth when G4s are stabilized. For example, GO term analysis revealed that of the 86 genes found to have ≤ -1.5 log_10_(n.w.r.r.) values, terms related to translation such as ribosome small subunit assembly and translation were enriched 15.97-fold and 4.73-fold, respectively, with several other translation-related terms having fold-enrichment values within that range (Table S2). Additionally, in the GO term analysis assessing genes with log_10_(n.w.r.r.) ≥ 1.5, cytoplasmic translation was found to be under-represented, with < 0.01-fold enrichment for genes annotated with this term.

Among the genes with increased insertions in G4-stabilizing conditions, *tufA* and *tufB* stood out. Collectively, *tufA* and *tufB* contained ∼10% of the weighted reads in NMM growth conditions and had log_10_(n.w.r.r.) of -3.52 and -3.19, respectively (**Figure 1C & D**), indicating a strong connection between their disruption and improved growth in the presence of the NMM. *tufA* and *tufB* insertion also aided growth in the pilot Tn-Seq screen with *tolC*^*+*^ cells, with log_10_(n.w.r.r.) of -0.53 and -0.75, respectively (Figure S2). These striking results necessitated a deeper investigation into *tufA* and *tufB*, both of which encode for the same protein (described further below).

To validate the impact of *tufA* or *tufB* deletion in desensitizing *E. coli* to NMM, *tufA* or *tufB* deletion strains were generated in the Δ*tolC* strain. As anticipated from the Tn-seq results, the plating efficiencies of the Δ*tolC tufA::kan* and Δ*tolC tufB::kan* strains were greater on NMM media than the Δ*tolC* parent strain, forming colonies at a 1000-fold more dilute culture than the Δ*tolC* control strain (**Figure 2A and B**). To determine if this effect was due to the G4-stabilization properties of NMM, plating on a second medium containing a structurally distinct G4 stabilizer, Braco-19, was measured. The Δ*tolC tufA::kan* and Δ*tolC tufB::kan* strains once again plated with higher efficiencies in the presence of Braco-19 than the Δ*tolC* control, albeit the effect was less stark than what was observed in NMM conditions (**Figure 2A and C**). Recent works suggests that NMM could stabilize RNA G4s to a greater extent than DNA G4s, which could cause the disparate sensitivity to NMM and Braco-19 (28,41).

**Figure 2.**
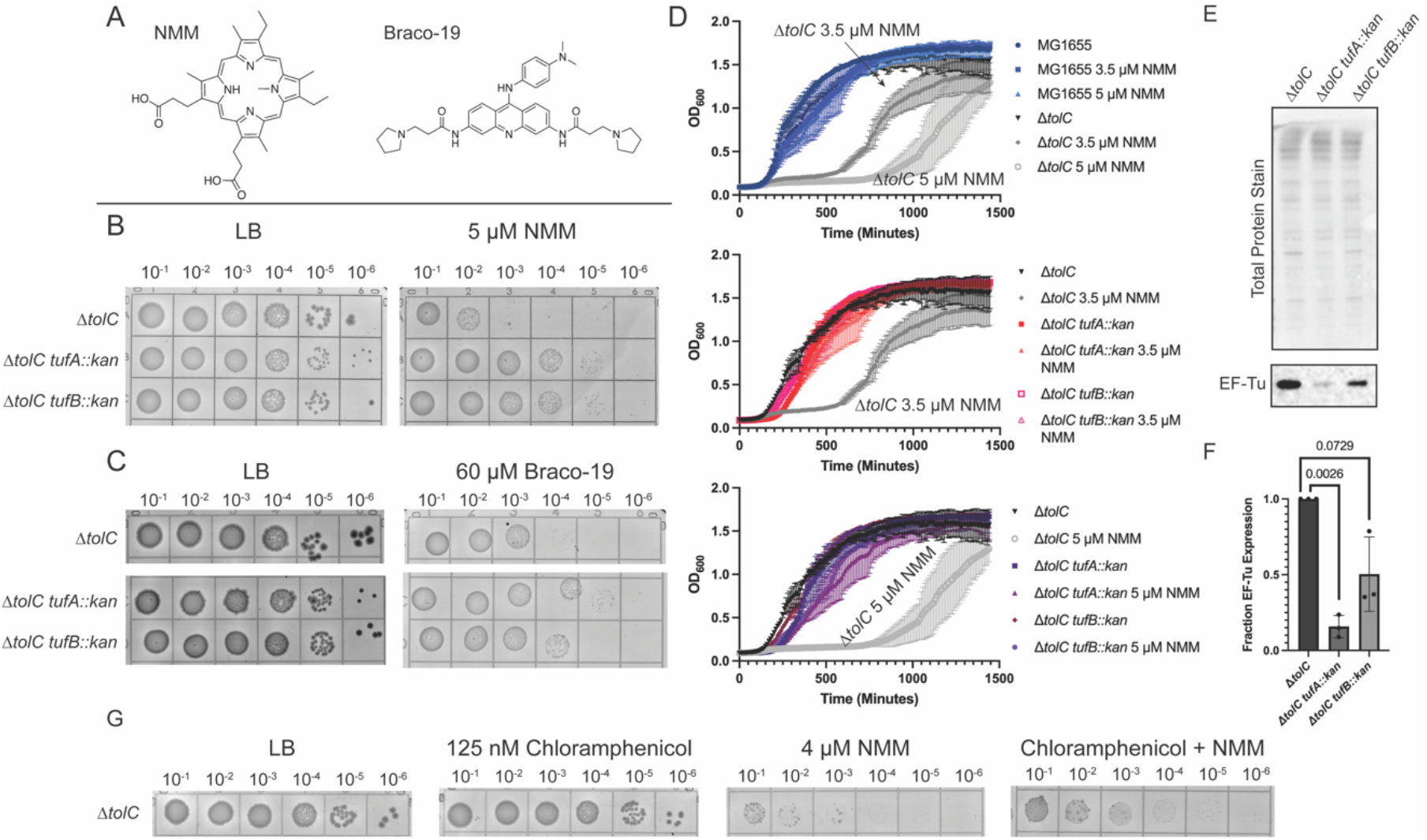
*tufA* and *tufB* deletions suppress growth defects in G4-stabilizing conditions. **(A)**. Structures of NMM and Braco-19, two structurally distinct G4 stabilizers used in this study. **(B)**. Spot plate experiments for Δ*tolC*, Δ*tolC tufA::kan*, and Δ*tolC tufB::kan* strains grown on Luria Broth (LB) or LB supplemented with 5 µM NMM. Each grid shows a 10-fold culture dilution, starting with cells plated at an OD_600 nm_ of 0.1. **(C)**. Spot plate experiments as shown in (B) but grown in the presence and absence of 60 µM Braco-19. **(D)**. Growth curves for MG1655, Δ*tolC*, Δ*tolC tufA::kan*, and Δ*tolC tufB::kan* grown in the presence and absence of 3.5 µM and 5 µM NMM. **(E)**. Western blot and total protein stain of Δ*tolC*, Δ*tolC tufA::kan*, and Δ*tolC tufB::kan* blotting for EF-Tu. **(F)**. Quantification of EF-Tu levels for *tufA* and *tufB* deletion compared to Δ*tolC* cells. **(G)**. Spot dilution plates plating MG1655 Δ*tolC* cells on LB-agar containing NMM, chloramphenicol, or both NMM and chloramphenicol.

Growth in liquid media was also examined ± NMM for MG1655 (*tolC*^*+*^), Δ*tolC*, Δ*tolC tufA::kan*, and Δ*tolC tufB::kan* strains (**Figure 2D**). Growth of the Δ*tolC* control strain was significantly delayed in the presence of NMM, and this delay increased with higher concentrations of NMM. Consistent with the plating results, the NMM-dependent lag phase was effectively eliminated by deletion of either *tufA* or *tufB*, although *tufA* and *tufB* deletion strains grew more slowly than the parent strain in the absence of NMM (**Figure 2D**). Collectively, these data indicate that deletion of either *tufA or tufB* desensitized *E. coli* to the effects of G4 stabilizers.

### Modulating translation elongation attenuates the effect of G4 stabilization

*tufA* and *tufB* both encode for elongation factor Tu (EF-Tu), an essential protein that escorts charged tRNAs to the ribosome during translation elongation (34,42,43). There are an ∼350,000 copies of EF-Tu in *E. coli*, making it the most abundant protein in the cell (34,43). EF-Tu-mediated delivery of aminoacylated tRNAs to the ribosome is thought to help govern the rate of translation elongation in *E. coli* (43). Previous work has demonstrated that deletion of *tufA* leads to a decrease in both cell growth and protein synthesis rates (35,36). We hypothesized that deletion of *tufA* or *tufB* would lead to decreased levels of EF-Tu in cells, which could alter translation elongation and contribute to cellular resistance to G4 stabilization.

Quantitative western blot analysis of Δ*tolC*, Δ*tolC tufA::kan*, and Δ*tolC tufB::kan E. coli* strains demonstrated that deletion of either *tufA* or *tufB* led to decreases in relative EF-Tu levels, with the *tufA* deletion having the greater impact (∼10-fold decrease in EF-Tu levels) (**Figure 2E & F**). Thus, a reduction in EF-Tu levels is correlated with the ability of the cell to overcome G4 stabilization. However, direct evidence that altering translation elongation through *tufA* or *tufB* deletion desensitizes cells to the effects of stabilized G4s was lacking.

To further test the idea that reduced translation rates allow *E. coli* to overcome, we next asked whether interfering with translation though a non-genetic intervention would improve cell growth in the presence G4 stabilizers. This possibility was tested by plating cells on media containing NMM and a sublethal dose of chloramphenicol, which interferes with bacterial translation elongation (44,45). Consistent with the idea that disruption to translation elongation helps to suppress the negative effects of stabilized G4s, the addition of chloramphenicol improved plating efficiency in the presence of NMM (Figure 2G).

### CRISPR interference screen identifies translation termination genes that are important under G4 stabilizing conditions

Because Tn-seq relies on gene disruptions, the approach cannot identify roles for essential genes in G4 stabilizing conditions. We therefore used a previously described CRISPR interference (CRISPRi) method (46) to examine the effects of individually reducing the levels of 536 targeted genes on cell growth on media supplemented with NMM. The screen identified 16 genes that, when targeted by CRISPRi machinery, sensitized cells to NMM, including genes involved in liposaccharide biosynthesis and transport (*lpxD, lpxB, lptE, lptC, lptA*), tRNA ligase activity (*leuS, aspS, valS*), ribosome large subunit assembly (*rplX*), protoporphyrinogen IX biosynthetic processes (*hemA*), methionine biosynthetic processes (*metE, metL*), DNA replication (*dnaA*), cell division (*ftsZ*), and, notably, in translation elongation and termination (*fusA* and *prfA*) (Figure S3). We were motivated to investigate *fusA* and *prfA* as our Tn-seq screen had already shown that translation is impacted by G4-stabilizers. Elongation factor G (EF-G, encoded by *fusA*) aids ribosome translocation, ribosome recycling, and is involved in a ribosome rescue pathway (47,48). Release factor 1 (RF1, encoded by *prfA*) is involved in initiating translation termination at UAG and UAA stop codons (47). Given the roles for EF-Tu in NMM resistance, EF-G an RF1 were further investigated.

To characterize the effects of *prfA* and *fusA* knockdowns with G4 stabilizers, *tolC* deletions were generated in strains containing the IPTG-inducible CRISPRi machinery targeted to *prfA* and *fusA*. Control strains that lacked CRISPRi or in which CRISPRi targeted a gene (*aroC*) that was not sensitized to NMM in the first screen were also tested. The strains were plated on media that included either IPTG, NMM, or both IPTG and NMM to assess how knockdowns of RF1 and EF-G impacted *E. coli* growth (**Figure 3A**). Both Δ*tolC* CRISPRi *prfA* and Δ*tolC* CRISPRi *fusA* strains grew poorly in the presence of IPTG and NMM compared to the Δ*tolC* and the Δ*tolC* CRISPRi *aroC* control strains (**Figure 3A**). These findings were mirrored in a background with competent efflux pumps (*tolC*^*+*^) (Figure S4).

**Figure 3.**
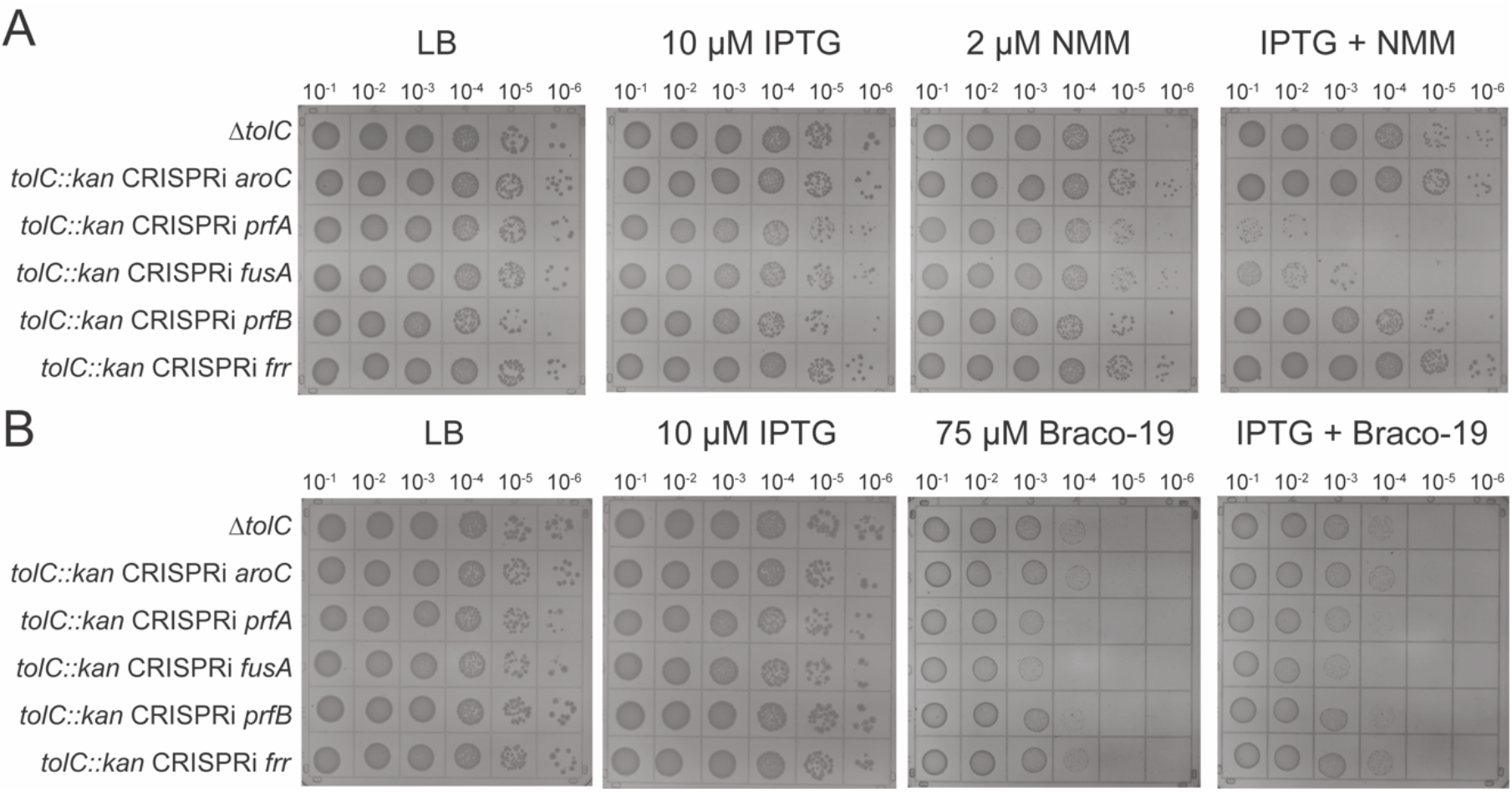
*prfA* and *fusA* knockdowns sensitize cells to G4 stabilizers. **(A)**. LB-agar spot dilution plates in the presence of IPTG (induce CRISPRi machinery), NMM, or both to assess impacts of knockdown strains on growth in G4-stabilizing conditions **(B)**. Same as (A) with Braco-19 as G4 stabilizing compound.

To test whether the growth defects of the Δ*tolC prfA* or Δ*tolC fusA* knockdown strains were specific to NMM or more broadly observed with a structurally distinct G4 stabilizer, the strains were also grown in the presence of Braco-19 (**Figure 3B**). These strains plated less efficiently than the control strains, although the effect was not as dramatic as that observed with NMM and was similar with or without IPTG. Nonetheless, the reduced plating efficiency of Δ*tolC* CRISPRi *prfA* and Δ*tolC* CRISPRi *fusA* strains indicate that the strains have a mild fitness defect in the presence of Braco-19 compared to the control strains. (28,41)

Given the importance of *prfA* and *fusA* in suppressing the effects of G4 stabilizers, we explored the impact of two additional essential translation termination genes, *prfB* and *rrf*. Release factor 2 (RF2, encoded by *prfB*) initiates translation termination at UGA and UAA codons (47) and ribosome recycling factor (RRF, encoded by *rrf*) is involved in ribosome recycling in conjunction with EF-G (47). Unlike *prfA* and *fusA*, CRISPRi-targeted knockdown of neither *prfB* nor *rrf* impacted *E. coli* cell plating efficiency on NMM or Braco-19 (**Figure 3**). Thus, the effects observed with *prfA* and *fusA* were specific. The fact that *prfA* and *fusA* knockdowns were detrimental in the presence of stabilized G4s whereas *prfB* and *rrf* were not could indicate that RF1 and EF-G have roles outside of translation termination and ribosome recycling that are important for overcoming stabilized G4s. Indeed, EF-G ‘s role in ribosome translocation and ribosome rescue could be important in the presence of stabilized G4s.

### Proteomic analysis reveals translation-related proteins are significantly altered in response to stabilized G4s

Identification of the importance of translation factors in overcoming G4s led to the question of how *E. coli* cells generally respond to chemicals that stabilize such structures. As a first step in addressing this question, a proteomic analysis was carried out to measure the quantitative effects of NMM on the levels of individual proteins in *E. coli*. Protein levels from early log-phase cultures of Δ*tolC* and Δ*tolC tufA::kan* strains grown in the presence or absence of NMM were measured to assess how reduced EF-Tu levels and NMM impacted expression. A total of 2509 proteins were detected in the dataset, approaching the limit of total proteins detected in previous *E. coli* proteomic studies (49,50) (Figure S5). Additionally, the replicates correlated well with one another assessed by principal component analysis and Pearson Correlation Coefficients (Figure S5).

GO term analyses were carried out for all proteins that were changed at least 2-fold in abundance when comparing two sample conditions (Tables S3-S9). Comparing each dataset, the largest overall difference was observed between the Δ*tolC* and NMM-treated Δ*tolC* cultures (**Figure 4** and S6). Hundreds of proteins levels were increased or decreased at least 2-fold in the presence of the G4-stabilizing compound. GO analyses carried out for proteins with statistically significant (as determined via q-value) ≥ 2-fold changes in the Δ*tolC* strain ± NMM (Tables S3 & S4) revealed that many of the significantly impacted pathways were related to translation, ribosome assembly, and ribosome biogenesis (**Figure 4A**). For example, ribosomal large subunit assembly and translation GO terms were 6.47 and 4.32-fold enriched, respectively, in proteins ≥ 2-fold less abundant in the Δ*tolC* NMM cultures compared to Δ*tolC* without NMM (Table S4). This striking trend led us to investigate the overlap between GO clusters for genes that had increased levels of insertions under NMM-growth condition in the Tn-seq experiment (GO terms identified in Table S1) and those that are downregulated in the presence of NMM. This analysis revealed many overlapping GO terms, including several terms associated with translation, ribosome assembly, and protein-RNA complex assembly (Table S5).

**Figure 4.**
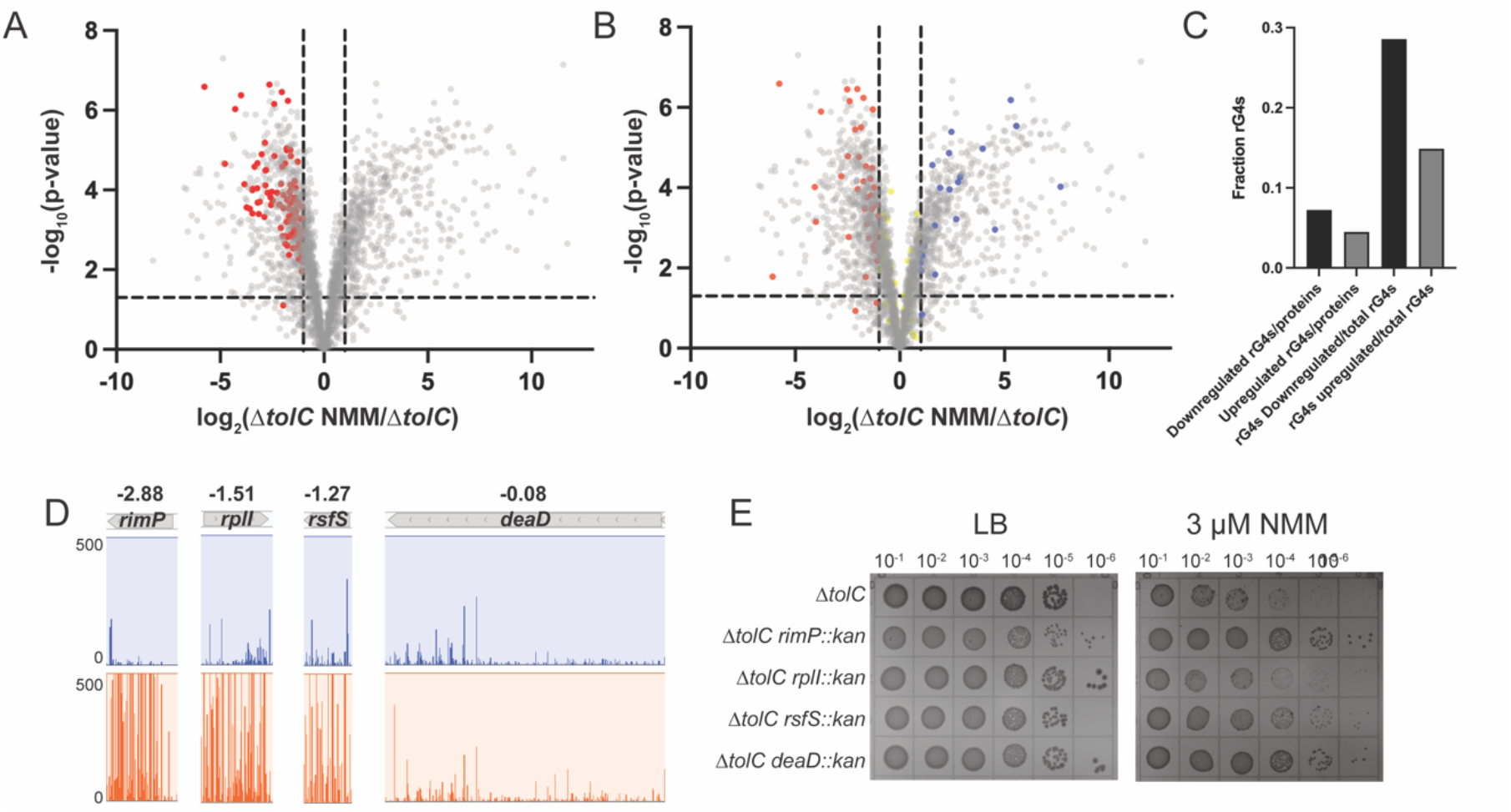
Proteomics of Δ*tolC* in the presence and absence of G4 stabilizer. **(A and B)**. Volcano plot of differences in individual protein levels between Δ*tolC* cells grown in the presence and absence of G4 stabilizer. Horizontal line indicates p-value of 0.05 and vertical lines indicate a 2-fold increase or decrease in protein levels. (A) Translation and ribosome biogenesis/assembly proteins that are downregulated in the presence of G4 stabilizer are depicted in red. (B) RNA G4s mapped onto the dataset, with red showing G4s in downregulated proteins, blue showing G4s in upregulated proteins, and yellow showing proteins that are not significantly changed in the proteomics experiment. **(C)** Quantification of the number of RNA G4s found in transcripts of proteins that are downregulated in the presence of NMM divided by total downregulated proteins and G4s found in proteins upregulated in the presence of NMM divided by total upregulated proteins. Additional quantification of the number of G4s found in the upregulated or downregulated proteomics category divided by the total number of G4s identified in *E. coli* (56). **(D)** Tn-seq results of four genes that were identified from proteomic analysis as proteins downregulated in the presence of NMM. Log_10_(ratio weighted reads) values are included above each transposon insertion profile. **(E)** Spot dilution plates of Δ*tolC* and Δ*tolC* strains harboring the gene deletions shown in (D) in the absence (left) or presence (right) of NMM.

To better understand the impact of reductions in protein levels of translation factors, four non-essential translation-related proteins found to be downregulated in the presence of NMM were selected for further analysis: RimP, RplI, RsfS, and DeaD (51–54). These proteins are associated with ribosome maturation or bind directly to the ribosome. Each is significantly less abundant in Δ*tolC* cells grown in the presence of NMM condition than in control conditions. Genes for three of these proteins, *rimP, rplI*, and *rsfS*, tolerated higher levels of insertions in the NMM-treated conditions than in control growth conditions, whereas the level of transposon insertions in the fourth, *deaD*, was nearly the same in the two conditions (**Figure 4D**). However, DeaD was one of the most downregulated proteins in the proteomic analysis, with a 54-fold decrease in expression in G4 stabilizing conditions.

To test possible effects of deletions of these genes on NMM-dependent growth, Δ*tolC deaD::kan*, Δ*tolC rimP::kan*, Δ*tolC rplI::kan*, and Δ*tolC rsfS::kan* strains were generated and grown ± NMM. Two of the strains (Δ*tolC deaD::kan* and Δ*tolC rimP::kan*) plated significantly more efficiently than the Δ*tolC* control strain in the presence of the G4 stabilizer NMM, whereas Δ*tolC rsfS::kan* plated modestly more efficient than the Δ*tolC* control (**Figure 4E**). These findings further bolster the idea that altering translation processes aids growth in G4-stabilizing conditions.

Since *tufA::kan* suppresses the negative effects of NMM, we predicted that the presence or absence of NMM would have little impact on gene expression within this strain. Indeed, comparing the Δ*tolC tufA::kan* ± NMM revealed only one protein (Spy, a chaperone (55)) that was differentially detected via mass spectrometry in these samples. This aligns well with our initial finding that Δ*tolC tufA::kan* cells are not sensitized to stabilized G4s.

### Mapping RNA G4s to the Δ*tolC* ± NMM proteomic dataset

One model that could explain why certain protein levels were reduced in the presence of NMM is that the compound is stabilizing G4s within transcripts which impedes translation. We investigated whether RNA G4s that are predicted to fold in *E. coli* correlated with less abundant proteins in the proteomic dataset. Previously, 168 RNA sequences within the coding sequences for proteins in *E. coli* transcripts were found to be able to fold into G4s (56). Mapping these putative RNA G4s onto the Δ*tolC* ± NMM proteomics results revealed that 56.5% of RNA G4s are contained in the coding sequences of proteins that are not significantly changed in the presence of NMM or not detected in the proteomics dataset (**Figure 4B-C**). However, for those elements that were found in transcripts of proteins with altered expression levels, 28.6% of G4s were found in transcripts encoding proteins that are downregulated in the presence of NMM compared to 14.9% of G4s found in proteins that are upregulated in G4 stabilizing conditions (**Figure 4C**). There was no connection between the positions of RNA G4s in the open reading frame and the effects on protein abundance (Figure S6F).

### Transcripts of several translation-related genes are less abundant in NMM-treated conditions

After determining proteome changes in each growth condition, we next sought to identify if changes in protein abundance were due to differences in transcript abundance. We utilized RNA-seq to determine how transcript levels varied in Δ*tolC* and Δ*tolC tufA::kan* cells grown ± NMM. RNA from cells grown to mid-log phase was sequenced an Illumina platform targeting 20 million paired end reads. Each sample had similar levels of normalized gene expression via DeSeq2 analysis (Figure S7) (57). Changes in transcripts were considered biologically and statistically significant with a ≥ 2-fold change in transcript detection and a statistically significant adjusted p-value of ≤ 0.05. Hundreds of significant transcript level differences were observed between cultures of: (*1*) Δ*tolC* cells ± NMM, (*2*) Δ*tolC* cells + NMM and Δ*tolC tufA::kan* cells + NMM, and (*3*) Δ*tolC* and Δ*tolC tufA::kan* cells (**Figures 5 &** S7C, and Tables S10-15). As observed in the proteomic analysis, far fewer transcriptome changes were observed between the Δ*tolC tufA::kan* ± NMM samples, with only one transcript (*mqo*) being less abundantly detected in Δ*tolC tufA::kan* + NMM. While several transcripts were more abundant in Δ*tolC tufA::kan* + NMM than cells grown without NMM, many of these transcripts mapped to genes associated with porphyrin biology (Table S16), likely an off-target effect of NMM.

**Figure 5.**
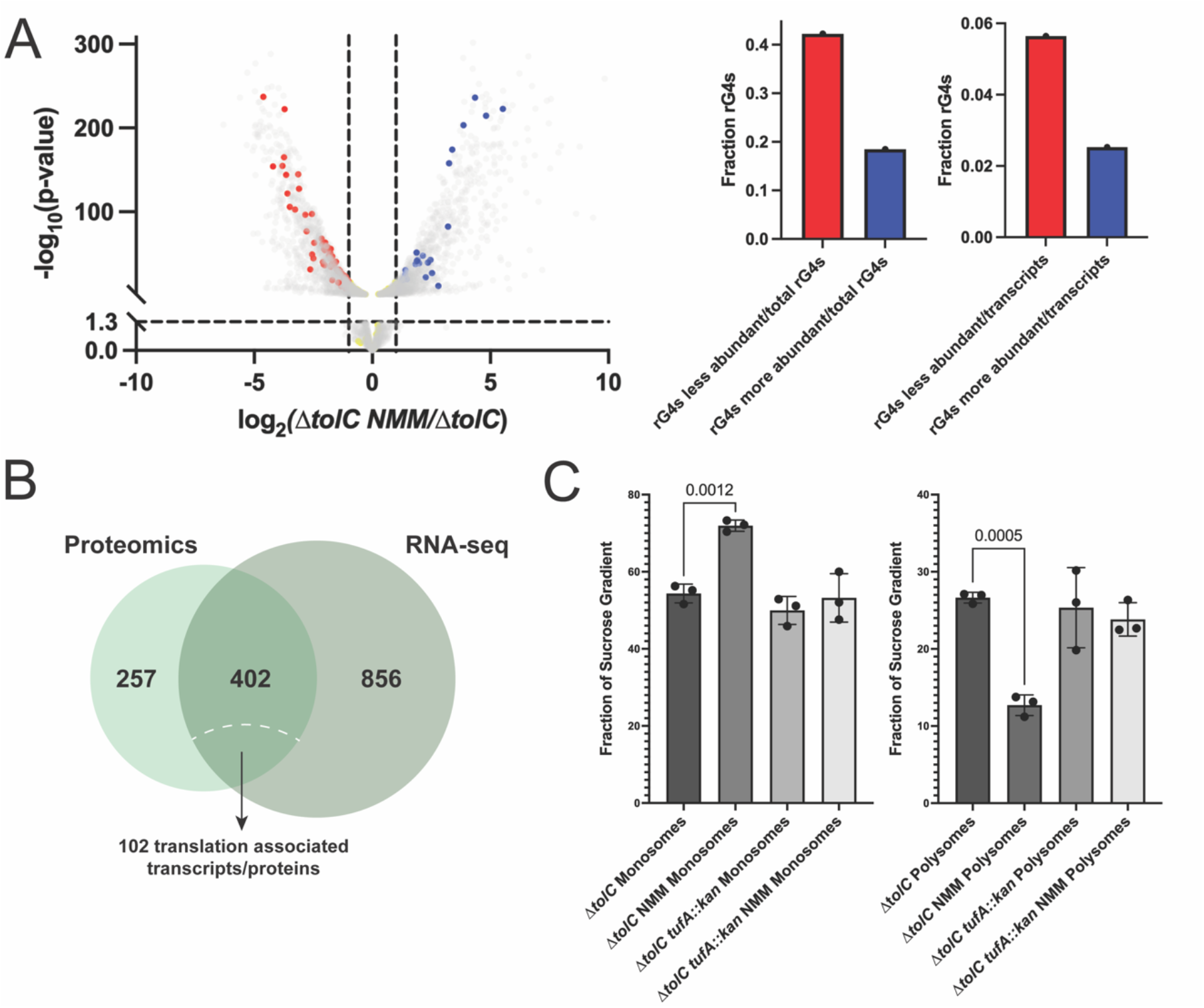
RNA-seq and sucrose gradient analysis of ribosomes reveal changes in translation in G4-stabilizing conditions. **(A)**. RNA G4 forming sequences mapped onto the transcriptomics volcano plot for Δ*tolC* ± NMM, with red points indicating transcripts less abundant in the presence of NMM that contain RNA G4s and blue points indicate transcripts that are more abundant in the presence of NMM and contain RNA G4s. Yellow points indicate transcripts not significantly altered by the presence of NMM and contain RNA G4s. Right: quantification of RNA G4s that were contained in more or less abundant transcripts shown as a fraction of total RNA G4s (left) and as a fraction of total upregulated or downregulated transcripts, respectively (right). **(B)** Overlap of proteins and transcripts identified as less abundant in the presence of NMM. **(C)** Quantification of the fraction of the area under the curve for monosomes and polysomes for Δ*tolC* and Δ*tolC tufA::kan* ± NMM cultures.

For Δ*tolC* cells ± NMM, many of the transcripts that were less abundant in the presence of NMM encode for proteins involved in translation (Table S11). Comparing the Δ*tolC* NMM transcriptomic and proteomic results, 402 genes had decreased transcript and protein levels in Δ*tolC* + NMM compared to Δ*tolC* (**Figure 5B**). Among these, over 100 are broadly involved in translation, indicating that at both at the transcript and protein level, gene expression is altered in response to NMM.

We next assessed whether RNA G-quadruplex forming sequences correlated to Δ*tolC* ± NMM transcripts level differences. Seventy-one RNA G4s were present in transcripts with reduced levels in the + NMM condition, 31 RNA G4s were found transcripts with increased levels in the + NMM condition, and 66 were found in transcripts that were not significantly changed (**Figure 5A**). Interestingly, this is different than what was observed for the proteomic dataset, where the majority of the RNA G4s mapped to proteins that were not significantly changed in the Δ*tolC* ± NMM dataset comparison (**Figure 4**). The presence of NMM may lead to stabilization of RNA G4s in these transcripts, leading to either degradation of the transcript or decreased translation of the implicated gene. However, decreases in transcript level do not necessarily lead to a rapid decrease in the associated protein levels, denoting a more nuanced relationship between transcript and protein levels that could stem from cellular protein stability and bacterial mRNA instability.

### Global ribosome occupancy is altered in the presence of NMM

To assess whether ribosome occupancy on transcripts was altered in the presence and absence of NMM, we used sucrose gradients to analyze ribosome occupancy from Δ*tolC* cells ± NMM and Δ*tolC tufA::kan* ± NMM. The fraction of monosomes in Δ*tolC* NMM cells was significantly increased compared to all other samples (**Figure 5C** and S8). This was matched with a subsequent decrease in the polysome fraction (**Figure 5C**). In contrast, the fractions of small and large subunit fractions were within experimental error for all conditions tested (Figure S8). NMM-treated samples from *tufA::kan* cultures matched those from untreated controls, indicating that a reduction of EF-Tu levels attenuates the difference in monosome/polysome ratio. The change in ratios indicates that translation is significantly altered by stabilized G4s, raising the possibility that RNA G4s are inducing ribosome collisions. While ribosome collisions are typically observed with an increase in polysome abundance, these collisions are also thought to signal for ribosome rescue, leading to clearance of the stalled polysomes (58). If G4 stabilization induces ribosome collisions, our data are consistent with slowing translation through either alteration of translation factor levels or chemical intervention (chloramphenicol) helping alleviate G4 toxicity. Additionally, this would indicate that factors involved in preventing ribosome collisions or rescuing ribosomes from these sites of damage could be important in the presence of G4 stabilizers. Consistent with this notion, genes encoding several ribosome rescue proteins (*arfA, arfB, hflX*, and *smpB*, (47,48)) were identified as modestly important genes in G4-stabilizing conditions in the Tn-seq screen (Figure S9). However, deletions of these genes or of *smrB*, a recently identified gene important in ribosome collision prevention (58) did not produce cells with altered NMM sensitivity (Figure S9), indicating that their impact on G4 resistance is less pronounced than other translation factors.

## Discussion

G4s are intriguing cellular regulatory elements with roles in all domains of life. As G4 stabilization and *in vivo* detection tools have advanced, many studies have focused on the effects of G4s in eukaryotic systems. However, the roles of G4s in bacteria have received much less attention (1). To better understand the impact of G4 structures on bacteria, we have used chemical-genetic approaches to probe how disruption of genes in *E. coli* affects growth under conditions where transient G4 structures are chemically stabilized. Gene disruptions altering translation factors had a profound impact on G4 tolerance. Transposon insertions in *tufA* of *tufB* genes, both encoding EF-Tu, were strongly selected for in G4-stabilizing conditions whereas CRISPRi-mediated suppression of RF1 or EF-G sensitized cells to stabilized G4s. Disruption of genes encoding translation/ribosome assembly factors DeaD, RimP, and RsfS also enhanced G4 tolerance.

Analysis of protein and transcript levels revealed significant changes in response to chemical stabilization of G4s, with downregulation of translation factors frequently occurring. Our results point to G4 structures in RNA and their interplay with translation processes as a potential regulatory feature in bacteria.

Discovery that alteration of the levels of multiple translation proteins in *E. coli* affect sensitivity to G4-stabilizing compounds strongly suggests that RNA G4 inhibition of protein synthesis underlies impaired growth in the presence of the G4 stabilizers. *tufA* or *tufB* deletion reduce EF-Tu levels in *E. coli*, which is correlated with enhanced cell growth in the presence of either of two structurally distinct G4 stabilizers. EF-Tu brings aminoacylated tRNAs to the ribosome during translation elongation, a function that has been proposed to be the rate-limiting step of translation and to modulate cell growth rates (34–36,43). In contrast, CRISPRi suppression of RF1 or EF-G levels led to decreased viability in the presence of NMM and, to a lesser extent, Braco-19. RF1 initiates translation termination at stop codons UAG and UAA and is also implicated in ribosome rescue (47). Interestingly, RF2, which recognizes stop codons UGA and UAA and is implicated in a ribosome rescue pathway, was not conditionally important in G4 stabilizing conditions (47,48). This indicates that other roles of RF1 that are distinct from RF2, such as roles in ribosome rescue or translation more generally, are likely important in G4 stabilizing conditions. The knockdown sensitivity of RF1 and not RF2 could also be due to residue substitution in RF2 (A246T) present in *E. coli* K-12 strains (59,60). This substitution causes reduced recognition of the UAA stop codon by RF2, making RF1 the major release factor to recognize UAA. This decrease in translation termination efficiency by RF2 could be linked to the conditional importance of RF1 in G4 stabilizing conditions.

EF-G is involved in a variety of functions, including ribosome recycling following translation termination, translocation of the ribosome during translation elongation, and in ribosome recycling in ribosome rescue (47). Reduction of expression of RRF, another protein involved in ribosome recycling after translation termination, did not alter NMM-sensitivity, indicating that the ribosome recycling role of EF-G in translation termination may not be involved in overcoming stabilized G4s. Instead, the role of EF-G in translocation or ribosome rescue may be important for preventing ribosomes stalling at G4s or ribosome collisions due to G4s.

Analysis of NMM-induced changes at the protein and transcript level revealed large effects in which several ribosome assembly, biogenesis, and translation are downregulated (**Figure 4 and 5**). Many of these pathways included genes selected for transposon insertion under + NMM growth conditions. Moreover, treatment with NMM perturbed the polysome/monosome ratio in Δ*tolC* cells. These results reinforce the notion that an overall reduction in translation improves growth in G4-stabilizing conditions.

Although the results presented here connect G4 stabilization and translation, perturbations to several ribosome rescue factors did not have an obvious impact on *E. coli* viability in G4-stabilizing conditions (Figure S9), indicating that only a subset of factors play major roles in dealing with G4 perturbations. An enhanced need for rescue factors could arise if stabilized RNA G4s induce ribosome collisions that are processed by specific ribosome rescue pathways.

Alternatively, if ribosome collisions are not induced by RNA G4s, it may be that pathways that process monosomes stalled at RNA G4s are move critical. Similar effects have been seen for ribosomes stalling at strong hairpin structures that may make ribosome A-sites less accessible for regulatory or rescue proteins (61). It is possible that stabilized RNA G4s could similarly impede ribosome rescue factor access to ribosome A sites, preventing rescue of ribosomes from sites of RNA G4s and inducing the increase in the fraction of monosomes observed with NMM (**Figure 5**). Such a model could help to explain how decreasing EF-Tu levels is beneficial for the cell, as lowering the pool of available EF-Tu would enhance ribosome A site accessibility for release/rescue.

RNA G4s in *E. coli* are all two-tetrad structures, which are less stable than the three-tetrad eukaryotic RNA G4s (1,56). *E. coli* may be sensitized to RNA G4 stabilization because they lack proteins that can efficiently unwind RNA G4s. Previous work found that inserting three-tetrad RNA G4s into *E. coli* was detrimental for growth (62). *E. coli* may have selected against highly stable RNA G4s because of an inability to unwind RNA G4s generally, or perhaps the RNA helicases in *E. coli* are unable to resolve three-tetrad RNA G4s. Nonetheless, it is possible that two-tetrad G4s play regulatory roles in *E. coli* in the absence of chemical stabilizers either when they transiently fold or when stabilized by G4-binding proteins.

Models explaining how *E. coli* can tolerate G4-stabilizing conditions through altered translation emerge from the present study (**Figure 6**). Chemically stabilizing RNA G4s can lead to either stalling of individual ribosomes at RNA G4 sites or could lead to ribosome collisions upstream of RNA G4s. Slowing translation elongation could allow ribosomes to densely coat mRNA, preventing RNA G4s formation/stabilization. Additionally, with reduced EF-Tu levels, ribosome A site accessibility could be increased, enhancing ribosome interaction with regulatory factors. If ribosome interaction with RNA G4s interferes with accessibility of A-sites for ribosome removal, lower EF-Tu could provide greater capacity for ribosome release or rescue factors to remove ribosomes from G4 stalling sites. Several copies of EF-Tu can bind to the ribosome at once, and lowering this pool of EF-Tu could provide access to factors like RF1 and EF-G to evacuate stalled ribosomes (34).

**Figure 6.**
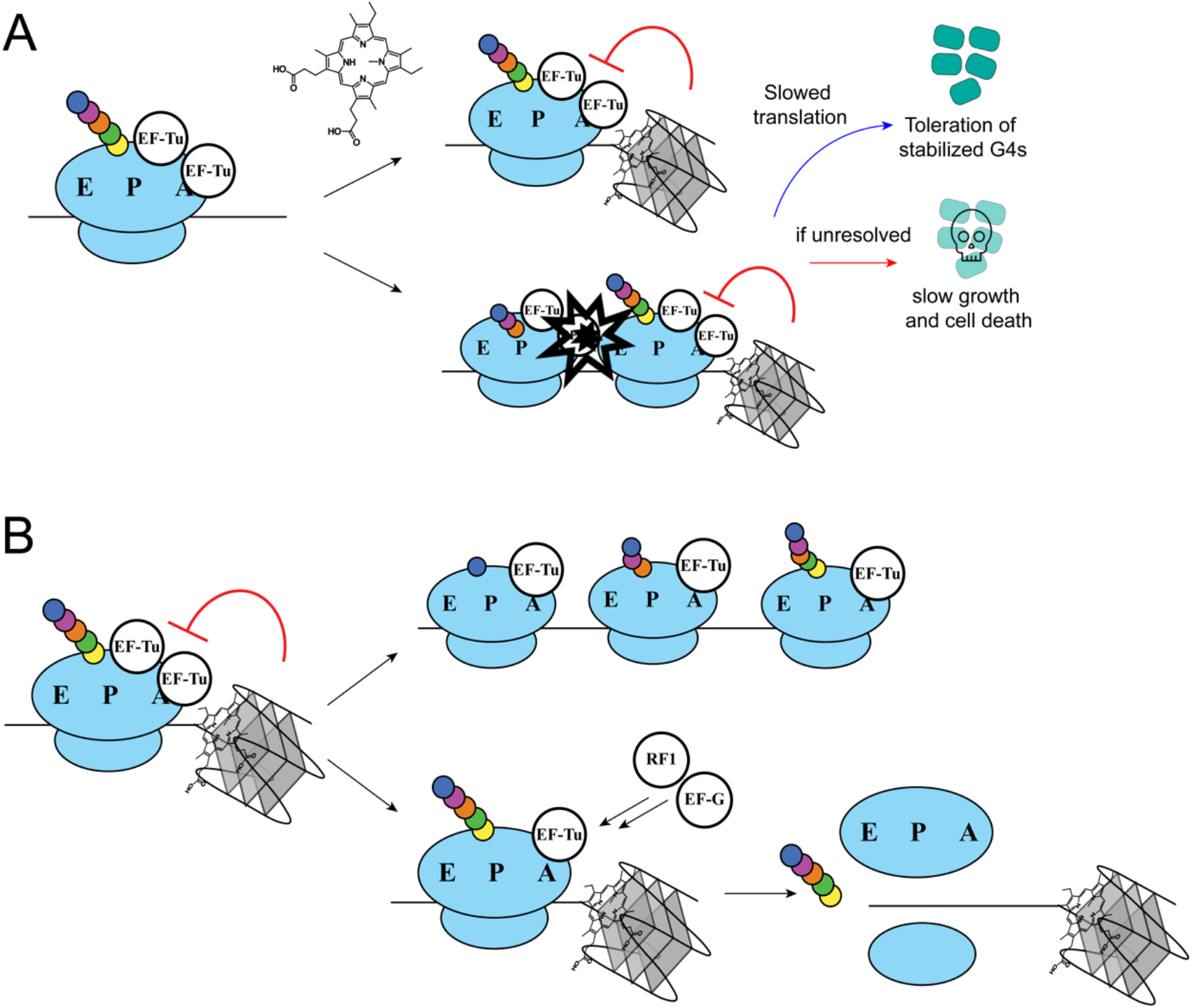
Model of how RNA G4s disrupt growth in *E. coli*. **(A)**. NMM stabilization of RNA G4s could lead to either stalling upstream of the RNA G4 or lead to potential ribosome collisions upstream of the RNA G4. This is lethal or leads to impaired cell growth if unresolved, but slowed translation can increase tolerance for stabilized RNA G4s. **(B)**. Altering translation by decreasing EF-Tu levels or the addition of chloramphenicol could allow for better toleration of G4 stabilization in multiple ways. One possibility is that slowing down translation would allow for ribosomes to densely coat RNA and prevent G4 folding or might slow down translation enough to prevent ribosome collisions. Another possibility is that lower EF-Tu levels would facilitate removal of stalled ribosomes from sites of stabilized G4s.

The results of this study provide new insights into the diverse roles of G4s in bacteria. Related studies in other bacterial species and in eukaryotes will be important for determining how wide-spread translation suppression can be used to overcome RNA G4 structures. If such mechanisms are unique to bacteria, chemically induced G4-linked growth defects could form a novel basis for development of antibacterial agents.

## Materials and Methods

### Strain construction

All cells used in this study are derived from an *Escherichia coli* MG1655 parent strain unless otherwise specified. For CRISPR interference strains, strains were a gift from Jason Peters (46). To generate *E. coli* knockout strains, P1 transductions were carried out using Keio collection strains as the donor strain (63,64). P1 phage lysate was grown on Keio collection donor strains, which was used to transduce the MG1655 strains or CRISPR interference strains (to make the *tolC* knockout of selected CRISPRi strains) which were sensitive to kanamycin. To validate strains, transductions were grown on LB plates supplemented with 50 µg/mL kanamycin and screened using colony PCR to validate proper insertion of the kanamycin resistance cassette. To remove the kanamycin resistant cassette from MG1655 *tolC::kan* to enable additional P1 transductions in this strain, MG1655 *tolC::kan* electrocompetent cells were generated and transformed with a plasmid encoding the FLP recombinase (pCP20) (65). Cells were recovered at 30 ºC and grown overnight on Super Optimal Broth (SOB) plates supplemented with 100 µg/mL ampicillin at 30 ºC. Single colonies from the plate were then grown overnight in LB at 43 ºC to promote loss of the temperature sensitive plasmid. A 10^−6^ dilution of cells was grown on LB plates at 30 ºC overnight to obtain individual colonies, which were then streaked onto LB only, LB supplemented with 100 µg/mL ampicillin, and LB supplemented with 50 µg/mL kanamycin.

Colonies that only grew on the LB without antibiotic plates were selected as MG1655 Δ*tolC* cells that were utilized for downstream applications.

### Transposome preparation and transposition

Transposome preparation was carried out as previously described (66,67). Briefly, the EZ-Tn5 <DHFR-1> transposon kit (Epicentre) and the E54K/M56A/L372P Tn5 hyperactive variant transposase were used for transposon mutagenesis. The Tn5 transposon was amplified using Phusion polymerase (NEB) and oligonucleotide oAM054. Transposase purification was carried out as previously described (66,68). Transposomes were prepared by incubating 2.5 pmol Tn5 DNA with 0.5 nmol Tn5 transposase for 3 hours at ambient temperature and then dialyzed into 1x TE buffer before electroporation.

Electrocompetent *E. coli* cells were generated as previously described (66). Briefly, *E. coli* were grown at 37 ºC to an OD_600_ ∼0.4 and cooled at 4 ºC for an hour. Cells were centrifuged at 10,750 rcf and pellets were washed in 10% glycerol three times. Cells were then resuspended in 2 mL GYT (10% (v/v) glycerol, 0.125% (w/v) yeast extract, and 0.25% (w/v) tryptone) before flash freezing and storing electrocompetent cells at -80 ºC. Five µL of transposome was combined with 100 µL of electrocompetent cells, electroporated, and recovered in 1 mL of SOC media at 37 ºC for 1 hour. Cells were plated on SOB-agar supplemented with 10 µg/mL trimethoprim to select for cells containing transposon insertions. Transposon mutants were pooled (∼200,000 colonies) from plates using 2 mL of LB to scrape colonies off plates and then stored in 50% glycerol at -80 ºC.

### Selection of tolerated transposon mutations in G4 stabilizing conditions

For the pilot Tn-seq experiment (Figure S1), a Tn5 transposase generated library in MG1655 *sulB103* was utilized (67). This library was diluted from a glycerol stock 1:10,000 in fresh LB and 250 µL of dilution was plated on SOC plates and SOC plates supplemented with 10 µM NMM. Plates were grown overnight at 37 ºC and there were estimated ∼100,000 colonies grown on SOC alone and ∼150,000 colonies grown on NMM supplemented plates. Colonies were pooled with LB and samples were diluted to an OD_600_ of ∼4.0 and 1 mL of concentrated cells underwent genomic DNA preparation using the Wizard Genomic DNA Purification Kit (Promega). DNA was quantified using the QuantiFluor ONE dsDNA System (Promega). Genomic DNA underwent shearing to ∼200 bp fragments via sonication and the gDNA fragments were prepared for sequencing using the NEBNext Ultra II DNA Library Prep Kit for Illumina (NEB). Bead-based size selection was employed to enrich for 200 bp fragments and the fragments then underwent a 17-cycle splinkerette PCR using oAM055 as the forward primer and either oAM068 (control) or oAM069 (NMM selected) as the reverse primer for barcoding and multiplexing (67). An additional bead-based size selection was used to clean up the sample before sequencing at the University of Michigan with a MiSeq platform. Primers oAM058 and oAM059 were used as unique sequencing primers for the control and NMM treated condition, respectively.

### Transposon sequencing with Δ*tolC* cells

For the subsequent Tn-seq experiment utilizing an MG1655 Δ*tolC* strain, transposomes were prepared as described above and the library was prepared as before (with the exception of using 1 µg/mL trimethoprim for selection) generating ∼500,000 transposon insertion mutants. MG1655 Δ*tolC* electrocompetent cells were generated as described above.

To select for transposon insertion mutations in control and NMM-treated conditions, libraries were grown on either SOB-agar plates or SOB-agar plates supplemented with 5 µM NMM. To ensure proper coverage after re-selection on plates, ∼1.5 million colonies were collected either from the SOB-agar plates and the SOB-agar plates supplemented with NMM and split into three libraries each. Libraries were passaged a second time in the presence and absence of NMM to generate a second passage library of ∼1.5 million colonies and split into three libraries each.

To prepare DNA for sequencing, libraries were prepared as described in the previous section. Each library was diluted to an OD_600_ of ∼4.0 and 1 mL of concentrated cells underwent genomic DNA preparation using the Wizard Genomic DNA Purification Kit (Promega). DNA was quantified using the QuantiFluor ONE dsDNA System (Promega). Genomic DNA underwent shearing to ∼200 bp fragments via sonication and the gDNA fragments were prepared for sequencing using the NEBNext Ultra II DNA Library Prep Kit for Illumina (NEB). Bead-based size selection was employed to enrich for 200 bp fragments and the fragments then underwent a 20-cycle splinkerette PCR using a Tn5-enrihcing forward primer (oAM55) and custom reverse primers for multiplexing (67). A final bead-based size selection was used to select for the correct length DNA. DNA was sequenced at the University of Michigan Advanced Genomics Core using a NextSeq platform (Illumina) with a custom read primer (oAM58) reading the last 10 nt of the transposon.

PhiX174 DNA spike was added to the run to ensure sufficient sequence diversity on the flow cell. Then, a custom index read primer (oAM59) and standard Illumina primer were used to sequence the index reads and PhiX174, respectively.

### Data analysis for Tn-seq

Tn-seq analysis was done as described previously (67). Tn-seq sequencing was trimmed with fastx_trimmer.pl version 0.0.13.2 (http://hannonlab.cshl.edu/fastx_toolkit). The default parameters were used except the first base to keep (-f flag) was edited to 10 to take out the transposon sequence. Samples were then split with fastx_barcode_splitter.pl, version 0.013.2 (http://hannonlab.cshl.edu/fastx_toolkit) using a file that contained the individual barcode sequence and the sample ID, then the barcode was removed from each read in the FASTQ file using Cutadapt version 1.13 (69). FASTQ files that were trimmed were then aligned to the *E. coli* K-12 MG1655 genome (NC_000913.3) using Bowtie2, version 1.2 on default parameters (70).

Conditional importance or essentiality of genes was determined using TSAS, version 0.3.0 using Analysis_type2 for 2 sample analysis to compare transposon insertion profiles of NMM treated cells to cells grown without G4 stabilizer (40). The weighted reads were determined as previously described (40). The other parameters were kept at default settings.

### Assessing sensitivity to G4 stabilizers using spot plates

Spot plating experiments to assess sensitivity to G4 stabilizing compounds was carried out as previously described (26). Briefly, NMM and Braco-19 were prepared by resuspending the compounds in 18 MΩ ultra-pure water and NMM concentration was assessed using the molar extinction coefficient 145000 M^-1^ cm^-1^ at 379 nm (71).

NMM and Braco-19 solutions were stored at 4 ºC. IPTG solutions were made by resuspension in 18 MΩ ultra-pure water and stored at -20 ºC and chloramphenicol was resuspended in ethanol and stored at -20 ºC. G4 stabilizers, IPTG, or chloramphenicol were added to LB-agar at the indicated concentrations and stored in the dark. Five mL of each *E. coli* strain were grown overnight and diluted in fresh LB to an OD_600_ ∼1. For spot plating, 10^−1^ to 10^−6^ dilutions of strains were made in LB and 10 µL of each dilution was plated onto the LB spot plates. Spot plates were grown overnight and imaged on the Azure c600. Spot plates were done in triplicate.

### CRISPR interference screen

For CRISPR interference screen, strains from the CRISPRi library were grown in plates with 200 µL of LB supplemented with 10 µg/mL chloramphenicol and 4 µL from each glycerol stock of the library. Cells were grown overnight at 37 ºC and then stored at 4 ºC overnight. The following day, plates were shaken at 37 ºC for 5 minutes and then diluted 200-fold into fresh LB and shaken for 5 minutes to mix cells. Two µL were plated onto plates with LB-agar alone or LB-agar supplemented with 15 µM NMM, 10 µM IPTG, or 10 µM IPTG and 15 µM NMM together. Plates were grown overnight at 37 ºC and imaged the following day using the Azure c600.

### Assessing sensitivity to G4 stabilizers using growth curves

MG1655, Δ*tolC*, Δ*tolC tufA::kan*, and Δ*tolC tufB::kan* cells were grown overnight at 37 ºC in Luria Broth (LB). The next day, cells were diluted 100-fold into fresh LB and grown to an OD_600_ ∼0.2. Cells were then diluted 100-fold in a 96-well plate either in the presence or absence of NMM and OD_600_ of the cells was measured every ten minutes over 24 hours with continuous shaking at 37 ºC in a plate reader (BioTek Synergy H1). Growth curves were done in triplicate. The average of each growth condition was then plotted in Prism (10.2.0) with error bars representing the standard error of the mean.

### Western blots

Δ*tolC*, Δ*tolC tufA::kan*, and Δ*tolC tufB::kan* cells overnight cultures were diluted 100-fold in fresh Luria Broth (LB) and grown to an OD_600_ ∼0.3. One mL of cells were pelleted and resuspended in 50 µL of 1x sample buffer (0.8% SDS, 11.5% glycerol, 0.1 M Tris pH 6.8, 0.286 M βME, 0.01% bromophenol blue). Five µL of undiluted sample and 5 µL of sample at various dilutions (1:2, 1:10, or 1:15) were loaded onto a 5-15% PAGE gel (brand) and run in 1x SDS running buffer.

Proteins were then transferred to a nitrocellulose membrane at 4 ºC for 1.5 hours in transfer buffer (25 mM Tris pH 8, 192 mM glycine, 0.03% SDS, 20% methanol). For total protein staining, membranes were incubated with 5 mL total protein stain (LI-COR) for 5 minutes at ambient temperature on a rocking platform before getting rinsed with wash solution (30% methanol, 6.7% acetic acid) and then imaged on the IR700 channel using the Azure c600. Following total protein stain, the membrane was blocked for 1 hour at room temperature in 5% dry milk in 1x PBS (137 mM NaCl, 2.7 mM KCl, 4.3 mM Na_2_HPO_4_, 1.47 mM KH_2_PO_4,_ pH 7.4) and then rinsed with 1x PBS. Membrane was then incubated for 1 hour at ambient temperature with anti-EF-Tu antibody (Hycult Biotech) at 2 µg/mL and then rinsed with 1x PBS. Membrane was then incubated with peroxidase conjugated goat anti-mouse antibody (Invitrogen) at 0.1 µg/mL and then rinsed with 1x PBS. The Amersham ECL Prime Western Blotting detection kit was then used to visualize the blot on the Azure c600 using the chemiluminescence setting. Intensity of bands for total protein normalization and for EF-Tu blots was carried out using ImageJ and plotted using Prism (10.2.0). Significance was determined in Prism using the Welch ‘s two-tailed t-test. Western blots and total protein staining were done in triplicate.

### Proteomics cell growth and cell lysis

Δ*tolC* and Δ*tolC tufA::kan* cells were grown overnight and then diluted 100-fold in fresh LB and grown to an OD_600_ of ∼0.2. Cells were then diluted 100-fold and grown in LB ± 3.5 µM NMM. Cells were grown to an OD_600_ ∼0.2-0.4 and then 45 mL of cells were pelleted via centrifugation. The pellets were washed with 5 mL of 1x PBS to remove any residual media and then pelleted again. Cells pellets were stored at -80 ºC.

Cell lysis was initiated by resuspension in 250 µl of lysis buffer (8 M urea and 100 mM Tris pH 8, supplemented with cOmplete™ protease inhibitor cocktail (Roche) according to the manufacturer ‘s specifications). The cell suspension was then subjected to a two-step process to complete lysis: i) sonication via a probe sonicator for 1 min at medium intensity, followed by a 1 min incubation step on ice. ii) 250 µl of glass beads (1 mm diameter) were added to each sample, and samples were subjected to 4 repetitions of the following bead-beating protocol using a Retsch MM400 oscillation mill: 4 min of milling at 30 Hz, followed by a 1 min incubation step on ice. After lysis, samples were subjected to a clarifying spin and protein concentration was determined via BCA assay (Thermo Pierce). Next, 50 µg of each sample were transferred to a new tube and diluted in lysis buffer to a concentration of 1 mg/mL. To reduce and alkylate cysteine residues, samples were adjusted to 10 mM TCEP and 40 mM chloroacetamide and incubated for 30 min at ambient temperature. Subsequently, sample were diluted in 100 mM Tris pH 8 to a urea concentration of 4 M, followed by the addition of 1 µg LysC (Wako Chemicals) and a four hour incubation at ambient temperature. For o/n tryptic digestion at ambient temperature, 50 µg of trypsin (Promega) were added after diluting samples further down to a urea concentration of 1 M. Next morning, digest was stopped by adjusting samples to 1% TFA and peptides were purified through Strata-X solid phase extraction cartridges (Phenomenex). Peptide eluates were then dried in a vacuum concentrator and afterwards resuspended in 0.2% FA to a concentration of 1 mg/ml, ready for MS analysis.

### LC-MS analysis

For LC-MS analysis, the following setup was employed: a Vanquish Neo UHPLC System was coupled to an Orbitrap Astral mass spectrometer via a Nanospray Flex ionization source (all Thermo Scientific), operated at a source voltage of 2 kV. The Vanquish Neo was equipped with a 40 cm fused silica capillary column (75 μm i.d. and 360 μm o.d., Polymicro Technologies) and pulled, etched and packed in-house using 1.7 µm C18 particles (Waters) as described previously (72). Individual MS experiments were conducted by separating 1 µg of peptides at a flow rate of 300 nL/min at 55°C via a 2 h gradient (Mobile phase A: 0.2% FA, mobile phase B: 0.2% FA, 80% acetonitrile). MS experiments were conducted using a data-dependent acquisition (DDA) regime combining Orbitrap (MS1) and ion trap (MS2) scans under the following parameters: MS1 scans were recorded at a resolution of 240k, a scan range of 300-1350 m/z and a normalized AGC target of 250% with a maximum injection time of 50 ms. MS2 scan were recorded with an isolation window of 0.5 m/z, an HCD collision energy of 23%, at a “Turbo” scan rate speed, a scan range of 150-1350 m/z and a normalized AGC target of 250% with a maximum injection time of 14 ms. MIPS, charge state and dynamic exclusion filters were employed.

### Proteomics data processing and analysis

LC-MS analysis resulted in 12 Thermo RAW files (2 strains x 2 growth conditions x 3 biological replicates), which were processed with MaxQuant, version 2.4.2.0 (73). MaxQuant was run using the default settings, with the following specifications and changes: i) RAW files were searched against the *E*.*coli* Uniprot reference proteome (Organism ID: 83333, downloaded in Sep 2023). All 12 files were searched together but separated into four experiments with three biological replicates per experiment. ii) Under *Group-specific parameters*, LFQ was enabled. Under *Global parameters, Min. unique peptides* was set to 1 and *Match between runs* as well as *iBAQ* were enabled. MaxQuant output files were subsequently analyzed via Perseus, version 2.0.11.0 (74). In Perseus, LFQ intensities were log-transformed, followed by a data filtering step, requiring three out of three valid LFQ intensity values for at least one of the four experiments. Next, missing values were imputed from a normal distribution using Perseus ‘ default settings. Differences across experiments were then assessed via a two-sided two-sample t-test. To address the multiple testing problem, a permutation-based false discovery rate calculation based on 250 randomizations was included. Our analysis yielded mean log2 LFQ intensity ratios, p-values and q-values for 2,468 protein groups.

### RNA-seq sample growth and sequencing

Δ*tolC* and Δ*tolC tufA::kan* cells were grown overnight, back diluted 100-fold into fresh LB, and grown to an OD_600_ of ∼0.2. Cells were then diluted 100-fold into 100 mL fresh LB ± 3.5 µM NMM. Cells were then grown to an OD_600_ of ∼0.2-0.4. Cells were harvested via centrifugation at 3214 x g at 4 ºC for 10 minutes. Pellets were transferred to 1.7 mL tubes and flash frozen in LN_2_ and stored at -80 ºC.

Pellets were submitted to Genewiz for RNA extraction and library preparation using their RNA-seq with rRNA depletion package. Sequencing was done using the Illumina 2×150 bp platform targeting 20 million paired-end reads per sample. Data analysis was done through Genewiz using DeSeq2 to normalize datasets and generate plots shown in Figure S7 (75). P-values were determined via the Wald test p-value and adjusted p-values were determined via the Benjamini-Hochberg adjusted p-value.

### Sucrose gradients for polysome traces

Δ*tolC* and Δ*tolC tufA::kan* cells were grown in 50 mL of LB overnight. Cultures were then back diluted 100-fold into fresh LB, grown to an OD_600_ of ∼0.2, and then diluted 100-fold into 1 L of LB ± 3.5 µM NMM. Cells were then grown to an OD_600_ of ∼0.3-0.5. Cells were harvested via filtration with 0.45 µm filter (Whatman) and stored at -80 ºC.

Cells were then lysed via cryomilling in 1 mL of lysis buffer (20 mM Tris pH 8.0, 10 mM MgCl_2_, 100 mM NH_4_Cl, 5 mM CaCl_2_, 100 U/mL DNase I, 1 mM chloramphenicol) at 10 s^-1^ for 1 minute three times. Cells were clarified via centrifugation at 20,000 x g at 4 ºC.

For sucrose gradients, ∼12.5 AU of RNA were loaded onto a 10-50% sucrose gradient (20 mM Tris pH 8.0, 15 mM MgCl_2_, 100 mM NH_4_Cl, 2 mM DTT) prepared using the Biocomp gradient station. Sucrose gradients were ultracentrifuged at 201,000 x g at 4 ºC for 2.5 hours at maximum acceleration and deceleration in the SW 41 Ti rotor. Sucrose gradients were then fractionated on the Biocomp Gradient Station and A_260_ measurements were monitored. Sucrose gradients were done in triplicate.

To analyze the area under the curve (AUC) corresponding to the small subunit, large subunit, monosomes, and polysomes, the script from (76) was used in RStudio to quantify the fraction of the area under the curve for each component. This was done for each sucrose gradient and the AUC percentages were plotted in Prism (10.2.0). P values were determined using Welch ‘s two-tailed t-test.

## Author Contributions

Rachel R Cueny: Conceptualization, methodology, investigation, analysis, data curation, writing-original draft and review and editing, visualization. Andrew F Voter: Conceptualization, methodology, analysis, investigation, writing-review and editing. Aidan M McKenzie: methodology, investigation, and writing-review and editing. Marcel Morgenstern: methodology, investigation, analysis, and data curation. Michael Place: methodology, analysis, and investigation. Kevin Myers: methodology, investigation, analysis, and data curation. Jason Peters: resources and writing-review and editing. Joshua Coon: methodology, resources, supervision, funding acquisition. James L Keck: Conceptualization, methodology, resources, writing-original and review and editing, supervision, funding acquisition.

## Acknowledgments

The authors thank the Keck lab for critical evaluation of the manuscript. The authors would also like to thank Aude Trinquier and the Wang lab for training on the Biocomp Gradient Station Maker (funded by Supplement 3R35GM127088-04S1). This work was supported in part by the National Institute of General Medical Sciences (grants P41GM108538 and R35GM118110 to J.J.C.).

## References

1. Cueny RR, McMillan SD, Keck JL. G-quadruplexes in bacteria: insights into the regulatory roles and interacting proteins of non-canonical nucleic acid structures. Vol. 57, Critical Reviews in Biochemistry and Molecular Biology. Taylor and Francis Ltd.; 2022. p. 539–61.

2. Moyzis RK, Buckingham JM, Crams LS, Dani M, Deavent LL, Jones MD, et al. A highly conserved repetitive DNA sequence, (TTAGGG)n, present at the telomeres of human chromosomes. Proc Natl Acad Sci USA [Internet]. 1988;85:6622–6. Available from: https://www.pnas.org

3. Henderson E, Hardin CC, Walk SK, Tinoco L, Blackburn EH. Telomeric DNA Oligonucleotides Form Novel Intramolecular Structures Containing Guanine-Guanine Base Pairs. Vol. 51, Cell. 1987.

4. Greider CW, Blackburn EH. The Telomere Terminal Transferase of Tetrahymena Is a Ribonucleoprotein Enzyme with Two Kinds of Primer Specificity. Vol. 51, Cell. 1987.

5. Sen D, Gilbert W. Formation of parallel four-stranded complexes by guanine-rich motifs in DNA and its implications for meiosis. Nature. 1988;334(28):364–6.

6. Cech TR. G-strings at chromosome ends. Nature. 1988;332(28):777–8.

7. Di Antonio M, Ponjavic A, Radzevičius A, Ranasinghe RT, Catalano M, Zhang X, et al. Single-molecule visualization of DNA G-quadruplex formation in live cells. Nat Chem. 2020 Sep 1;12(9):832–7.

8. Paeschke K, Capra JA, Zakian VA. DNA Replication through G-Quadruplex Motifs Is Promoted by the Saccharomyces cerevisiae Pif1 DNA Helicase. Cell. 2011 May 27;145(5):678–91.

9. Lee WTC, Yin Y, Morten MJ, Tonzi P, Gwo PP, Odermatt DC, et al. Single-molecule imaging reveals replication fork coupled formation of G-quadruplex structures hinders local replication stress signaling. Nat Commun. 2021 Dec 1;12(1).

10. Sato K, Martin-Pintado N, Post H, Altelaar M. Multistep mechanism of G-quadruplex resolution during DNA replication. Vol. 7, Sci. Adv. 2021.

11. Holder IT, Hartig JS. A matter of location: Influence of G-quadruplexes on escherichia coli gene expression. Chem Biol. 2014 Nov 20;21(11):1511–21.

12. Cogoi S, Xodo LE. G-quadruplex formation within the promoter of the KRAS proto-oncogene and its effect on transcription. Nucleic Acids Res. 2006;34(9):2536–49.

13. Siddiqui-Jain A, Grand CL, Bearss DJ, Hurley LH. Direct evidence for a G-quadruplex in a promoter region and its targeting with a small molecule to repress c-MYC transcription. Proceedings of the National Academy of Sciences [Internet]. 2002;99(18):11593–8. Available from: https://www.pnas.org

14. Lee CY, McNerney C, Ma K, Zhao W, Wang A, Myong S. R-loop induced G-quadruplex in non-template promotes transcription by successive R-loop formation. Nat Commun. 2020 Dec 1;11(1).

15. Myong S, Lee CY, Joshi M, Wang A. 5’UTR G-quadruplex structure enhances translation in size dependent manner. Res Sq [Internet]. 2023;1–26. Available from: 10.21203/rs.3.rs-3352233/v1

16. Bugaut A, Balasubramanian S. 5′-UTR RNA G-quadruplexes: Translation regulation and targeting. Vol. 40, Nucleic Acids Research. 2012. p. 4727–41.

17. Arora A, Suess B. An RNA G-quadruplex in the 3′ UTR of the proto-oncogene PIM1 represses translation. RNA Biol. 2011;8(5):802–5.

18. Yadav P, Kim N, Kumari M, Verma S, Sharma TK, Yadav V, et al. G-quadruplex structures in bacteria: Biological relevance and potential as an antimicrobial target. Vol. 203, Journal of Bacteriology. American Society for Microbiology; 2021.

19. Baran N, Pucshansky L, Marco Y, Benjamin S, Manor H. The SV40 large T-antigen helicase can unwind four stranded DNA structures linked by G-quartets. Nucleic Acids Res. 1997;25(2).

20. Sun H, Karow JK, Hickson ID, Maizels N. The Bloom’s syndrome helicase unwinds G4 DNA. Journal of Biological Chemistry. 1998 Oct 16;273(42):27587–92.

21. Harrington C, Lan Y, Akman SA. The Identification and Characterization of a G4-DNA Resolvase Activity*. J Biol Chem [Internet]. 1997;272(39):24631–6. Available from: http://www.jbc.org

22. Sun H, Bennett RJ, Maizels N. The Saccharomyces cerevisiae Sgs1 helicase efficiently unwinds G-G paired DNAs. Nucleic Acids Res [Internet]. 1999;27(9). Available from: https://academic.oup.com/nar/article/27/9/1978/2847953

23. Voter AF, Qiu Y, Tippana R, Myong S, Keck JL. A guanine-flipping and sequestration mechanism for G-quadruplex unwinding by RecQ helicases. Nat Commun. 2018 Dec 1;9(1).

24. Chen MC, Tippana R, Demeshkina NA, Murat P, Balasubramanian S, Myong S, et al. Structural basis of G-quadruplex unfolding by the DEAH/RHA helicase DHX36. Nature. 2018 Jun 13;558(7710):465–83.

25. Cueny RR, Varma S, Schmidt KH, Keck JL. Biochemical properties of naturally occurring human bloom helicase variants. PLoS One. 2023 Jun 1;18(6 JUNE).

26. Paul T, Voter AF, Cueny RR, Gavrilov M, Ha T, Keck JL, et al. E. coli Rep helicase and RecA recombinase unwind G4 DNA and are important for resistance to G4-stabilizing ligands. Nucleic Acids Res. 2020 Jul 9;48(12):6640–53.

27. Jain A, Vale RD. RNA phase transitions in repeat expansion disorders. Nature. 2017 Jun 8;546(7657):243–7

28. Mei Y, Deng Z, Vladimirova O, Gulve N, Johnson FB, Drosopoulos WC, et al. TERRA G-quadruplex RNA interaction with TRF2 GAR domain is required for telomere integrity. Sci Rep. 2021 Dec 1;11(1).

29. Rankin S, Reszka AP, Huppert J, Zloh M, Parkinson GN, Todd AK, et al. Putative DNA quadruplex formation within the human c-kit oncogene. J Am Chem Soc. 2005 Aug 3;127(30):10584–9.

30. Cahoon LA, Seifer SH. An Alternative DNA Structure Is Necessary for Pilin Antigenic Variation in Neisseria gonorrhoeae. Science (1979). 2009;325(5941):764–7.

31. Sechman E V., Rohrer MS, Seifert HS. A genetic screen identifies genes and sites involved in pilin antigenic variation in Neisseria gonorrhoeae. Mol Microbiol. 2005 Jul;57(2):468–83.

32. Marsico G, Chambers VS, Sahakyan AB, McCauley P, Boutell JM, Antonio M Di, et al. Whole genome experimental maps of DNA G-quadruplexes in multiple species. Nucleic Acids Res. 2019 May 7;47(8):3862–74.

33. Kaplan OI, Berber B, Hekim N, Doluca O. G-quadruplex prediction in E. coli genome reveals a conserved putative G-quadruplex-Hairpin-Duplex switch. Nucleic Acids Res. 2016 Nov 2;44(19):9083–95.

34. Mustafi M, Weisshaar JC. Simultaneous Binding of Multiple EF-Tu Copies to Translating Ribosomes in Live Escherichia coli. mBio. 2018 Jan 1;9(1):1–16.

35. Tubulekas I, Hughes D. Growth and translation elongation rate are sensitive to the concentration of EF-Tu. Mol Microbiol. 1993;8(4):761–70.

36. Gausing K. Rates of Growth, Ribosome Synthesis and Elongation Factor Synthesis in a tufa Defective Strain of E. coli. Mol Gen Genet. 1981;184:272–7.

37. Du D, Wang-Kan X, Neuberger A, van Veen HW, Pos KM, Piddock LJV, et al. Multidrug efflux pumps: structure, function and regulation. Vol. 16, Nature Reviews Microbiology. Nature Publishing Group; 2018. p. 523–39.

38. Tatsumi R, Wachi M. TolC-dependent exclusion of porphyrins in Escherichia coli. J Bacteriol. 2008 Sep;190(18):6228–33.

39. Turlin E, Heuck G, Simões Brandão MI, Szili N, Mellin JR, Lange N, et al. Protoporphyrin (PPIX) efflux by the MacAB-TolC pump in Escherichia coli. Microbiologyopen. 2014 Dec 1;3(6):849–59.

40. Burger BT, Imam S, Scarborough MJ, Noguera DR, Donohue TJ. Combining Genome-Scale Experimental and Computational Methods To Identify Essential Genes in Rhodobacter sphaeroides. mSystems. 2017 Jun 27;2(3).

41. Roschdi S, Yan J, Nomura Y, Escobar CA, Petersen RJ, Bingman CA, et al. An atypical RNA quadruplex marks RNAs as vectors for gene silencing. Nat Struct Mol Biol. 2022 Nov 1;29(11):1113–21.

42. Piepenburg O, Pape T, Pleiss JA, Wintermeyer W, Uhlenbeck OC, Rodnina M V. Intact aminoacyl-tRNA is required to trigger GTP hydrolysis by elongation factor Tu on the ribosome. Biochemistry. 2000;39(7):1734–8.

43. Klumpp S, Scott M, Pedersen S, Hwa T. Molecular crowding limits translation and cell growth. Proc Natl Acad Sci U S A. 2013 Oct 15;110(42):16754–9.

44. Walsh C. Antibiotics: Actions, Origins, Resistance. ASM Press; 2003. 1–335 p.

45. Wilson DN. Ribosome-targeting antibiotics and mechanisms of bacterial resistance. Nat Rev Microbiol. 2014;12(1):35–48.

46. Silvis MR, Rajendram M, Shi H, Osadnik H, Gray AN, Cesar S, et al. Morphological and transcriptional responses to CRISPRi knockdown of essential genes in Escherichia coli. mBio. 2021 Oct 1;12(5).

47. Buskirk AR, Green R. Ribosome pausing, arrest and rescue in bacteria and eukaryotes. Vol. 372, Philosophical Transactions of the Royal Society B: Biological Sciences. Royal Society; 2017.

48. Keiler KC. Mechanisms of ribosome rescue in bacteria. Vol. 13, Nature Reviews Microbiology. Nature Publishing Group; 2015. p. 285–97.

49. Mori M, Zhang Z, Banaei-Esfahani A, Lalanne J, Okano H, Collins BC, et al. From coarse to fine: the absolute Escherichia coli proteome under diverse growth conditions. Mol Syst Biol. 2021 May;17(5).

50. Mateus A, Hevler J, Bobonis J, Kurzawa N, Shah M, Mitosch K, et al. The functional proteome landscape of Escherichia coli. Nature. 2020 Dec 17;588(7838):473–8.

51. Häuser R, Pech M, Kijek J, Yamamoto H, Titz B, Naeve F, et al. RsFA (YbeB) proteins are conserved ribosomal silencing factors. PLoS Genet. 2012 Jul;8(7).

52. Naganathan A, Wood MP, Moore SD. The large ribosomal subunit protein L9 enables the growth of EF-P deficient cells and enhances small subunit maturation. PLoS One. 2015 Apr 16;10(4).

53. Charollais J, Dreyfus M, Iost I. CsdA, a cold-shock RNA helicase from Escherichia coli, is involved in the biogenesis of 50S ribosomal subunit. Nucleic Acids Res. 2004;32(9):2751–9.

54. Nord S, Bylund GO, Lövgren JM, Wikström PM. The RimP Protein Is Important for Maturation of the 30S Ribosomal Subunit. J Mol Biol. 2009 Feb 27;386(3):742–53.

55. Mitra R, Gadkari V V., Meinen BA, van Mierlo CPM, Ruotolo BT, Bardwell JCA. Mechanism of the small ATP-independent chaperone Spy is substrate specific. Nat Commun. 2021 Dec 1;12(1).

56. Shao X, Zhang W, Umar MI, Wong HY, Seng Z, Xie Y, et al. RNA G-quadruplex structures mediate gene regulation in bacteria. mBio. 2020 Jan 1;11(1).

57. Love MI, Huber W, Anders S. Moderated estimation of fold change and dispersion for RNA-seq data with DESeq2. Genome Biol. 2014 Dec 5;15(12).

58. Saito K, Kratzat H, Campbell A, Buschauer R, Burroughs AM, Berninghausen O, et al. Ribosome collisions induce mRNA cleavage and ribosome rescue in bacteria. Nature. 2022 Mar 17;603(7901):503–8.

59. Dincbas-Renqvist V, Engstrom A, Mora L, Heurgue-Hamard V, Buckingham R, Ehrenberg M. A post-translational modification in the GGQ motif of RF2 from Escherichia coli stimulates termination of translation. EMBO J [Internet]. 2000;19(24):6900–7. Available from: https://www.embopress.org

60. Uno M, Lto K, Nakamura Y. Functional specificity of amino acid at position 246 in the tRNA mimicry domain of bacterial release factor 2. Vol. 78, Biochimie. 1996.

61. Bao C, Loerch S, Ling C, Korostelev AA, Grigorieff N, Ermolenko DN. mRNA stem-loops can pause the ribosome by hindering A-site tRNA binding. Elife. 2020 May 1;9:1–67.

62. Guo JU, Bartel DP. RNA G-quadruplexes are globally unfolded in eukaryotic cells and depleted in bacteria. Science (1979). 2016 Sep 23;353(6306).

63. Thomason LC, Costantino N, Court DL. E. coli Genome Manipulation by P1 Transduction. Curr Protoc Mol Biol. 2007 Jul;79(1).

64. Baba T, Ara T, Hasegawa M, Takai Y, Okumura Y, Baba M, et al. Construction of Escherichia coli K-12 in-frame, single-gene knockout mutants: The Keio collection. Mol Syst Biol. 2006 May 16;2.

65. Cherepanov PP, Wackernagel W. Gene disruption in Escherichia coli: TcR and KmR cassettes with the option of Flp-catalyzed excision of the antibiotic-resistance determinant. Gene. 1995;158:9–14.

66. Byrne RT, Chen SH, Wood EA, Cabot EL, Cox MM. Escherichia coli genes and pathways involved in surviving extreme exposure to ionizing radiation. J Bacteriol. 2014;196(20):3534–45.

67. McKenzie AM, Henry C, Myers KS, Place MM, Keck JL. Identification of genetic interactions with priB links the PriA/PriB DNA replication restart pathway to double-strand DNA break repair in Escherichia coli. G3: Genes, Genomes, Genetics. 2022 Dec 1;12(12).

68. Bhasin A, Goryshin IY, Reznikoff WS. Hairpin Formation in Tn5 Transposition*. Journal of Biological Chemistry [Internet]. 1999;274(52):37021–9. Available from: http://www.jbc.org

69. Martin M. Cutadapt removes adapter sequences from high-throughput sequencing reads. EMBnet [Internet]. 2011;17(1):10–2. Available from: http://www-huber.embl.de/users/an-

70. Langmead B, Salzberg SL. Fast gapped-read alignment with Bowtie 2. Nat Methods. 2012 Apr;9(4):357–9.

71. Ren J, Chaires JB. Sequence and structural selectivity of nucleic acid binding ligands. Biochemistry. 1999 Dec 7;38(49):16067–75.

72. Shishkova E, Hebert AS, Westphall MS, Coon JJ. Ultra-High Pressure (>30,000 psi) Packing of Capillary Columns Enhancing Depth of Shotgun Proteomic Analyses. Anal Chem. 2018 Oct 2;90(19):11503–8.

73. Cox J, Mann M. MaxQuant enables high peptide identification rates, individualized p.p.b.-range mass accuracies and proteome-wide protein quantification. Nat Biotechnol. 2008 Dec;26(12):1367–72.

74. Tyanova S, Temu T, Sinitcyn P, Carlson A, Hein MY, Geiger T, et al. The Perseus computational platform for comprehensive analysis of (prote)omics data. Vol. 13, Nature Methods. Nature Publishing Group; 2016. p. 731–40.

75. Love MI, Huber W, Anders S. Moderated estimation of fold change and dispersion for RNA-seq data with DESeq2. Genome Biol. 2014 Dec 5;15(12).

76. Lecampion C, Floris M, Fantino JR, Robaglia C, Laloi C. An easy method for plant polysome profiling. Journal of Visualized Experiments. 2016 Aug 28;2016(114).

