## Supplemental Information for "Altering translation allows *E. coli* to overcome chemically stabilized G-quadruplexes"

### SI Figures and Tables for Altering translation allows *E. coli* to overcome chemically stabilized G-quadruplexes

#### Figures and Tables with legends

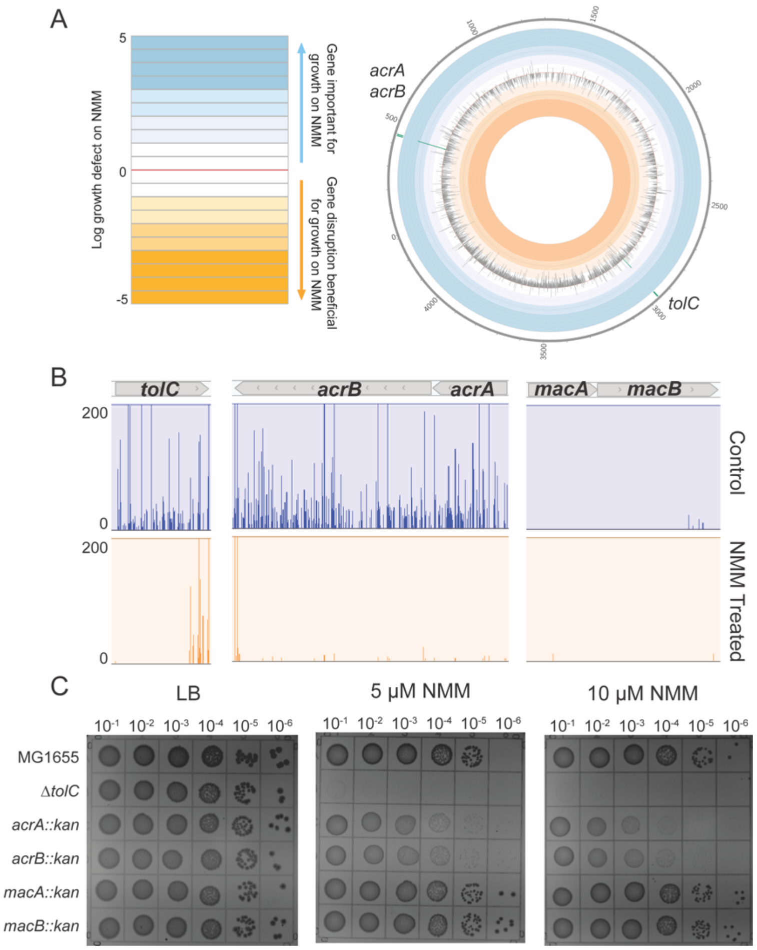


**Figure S1: Tn-Seq reveals that *E. coli* are sensitized to NMM when efflux pumps are compromised. (A)**. Circos plot showing $\log_{10}$(control weighted reads/NMM weighted reads) with genes involved in AcrAB-TolC efflux pump shown in green. **(B).** Mochiview plots of insertions across genes *acrA, acrB, tolC, macA,* and *macB*. **(C)**. Spot dilution plates of LB-agar and increasing concentrations of NMM with strains containing deletions of *tolC*, *acrA, acrB, macA,* and *macB*.

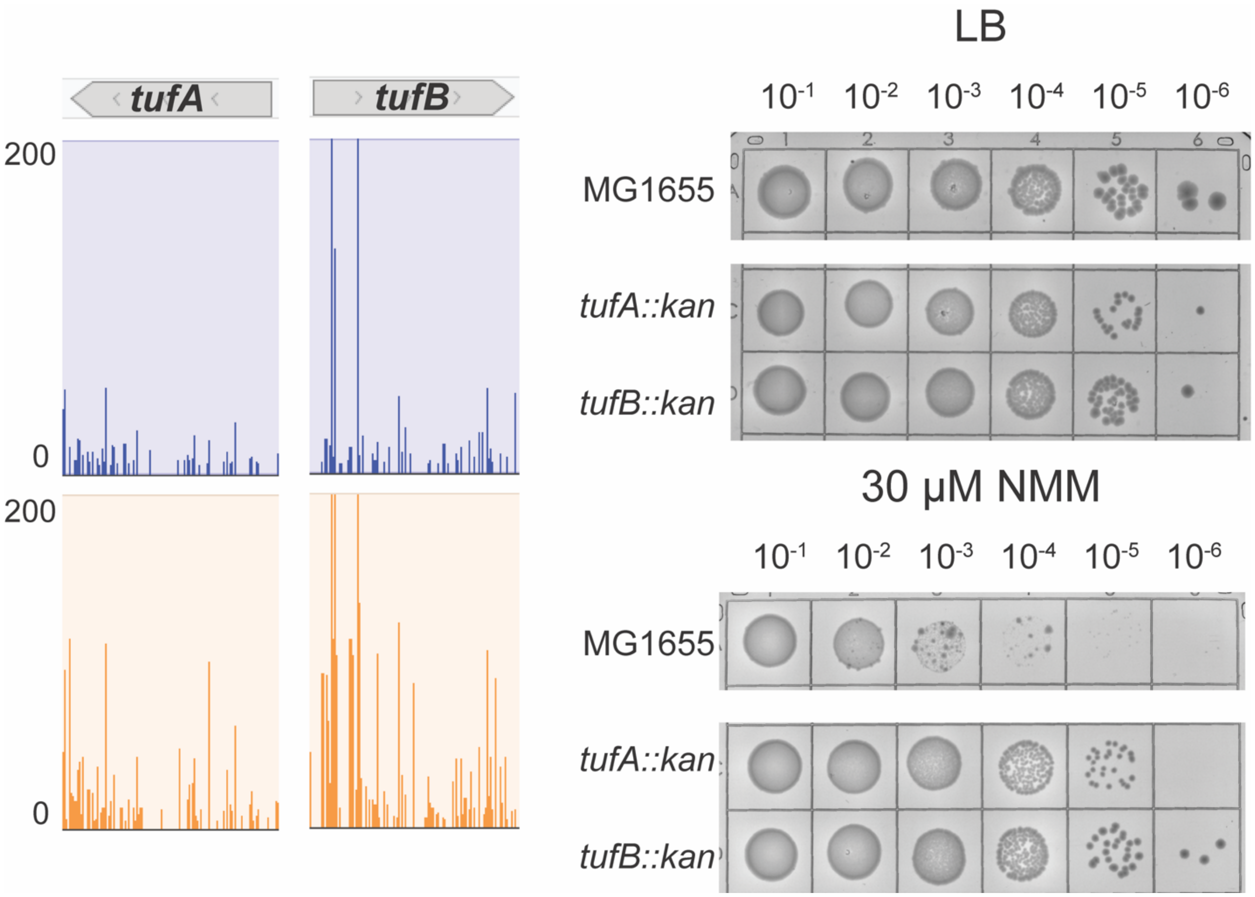


**Figure S2: Slowing down translation elongation allows cells to better in the presence of G4 stabilizer.** Mochiview insertion plots of *tufA* and *tufB* in the original Tn-seq screen and spot dilution plates containing the *tufA* and *tufB* gene deletions in the MG1655 cell background on LB-agar alone or LB-agar supplemented with NMM.

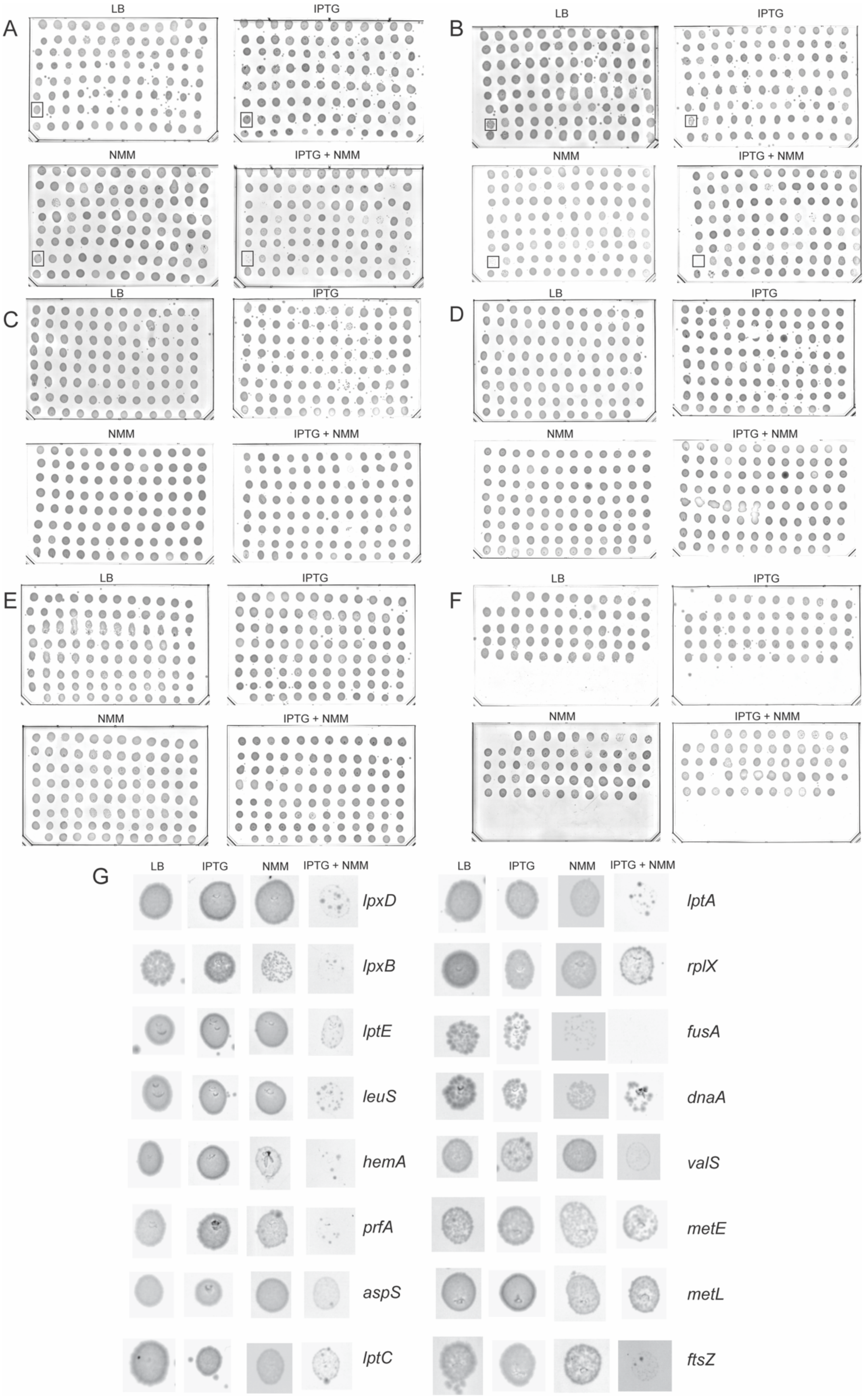
A
LB
IPTG
NMM
IPTG + NMM
B
LB
IPTG
NMM
IPTG + NMM
C
LB
IPTG
NMM
IPTG + NMM
D
LB
IPTG
NMM
IPTG + NMM
E
LB
IPTG
NMM
IPTG + NMM
F
LB
IPTG
NMM
IPTG + NMM
G
LB
IPTG
NMM
IPTG + NMM
lpxD
lpxB
lptE
leuS
hemA
prfA
aspS
lptC
lptA
rplX
fusA
dnaA
valS
metE
metL
ftsZ

**Figure S3: CRISPR interference plates testing sensitivity of knockdown strains to G4 stabilizer. (A-F)**. Test of each CRISPR interference strain for sensitivity to NMM, IPTG, and NMM IPTG and NMM together. The *prfA* knockdown strain is boxed in (A) and the *fusA* knockdown is boxed in (B). **(G).** Enlargement of spots for 16 gene knockdowns that had an impact on growth in NMM-treated conditions.

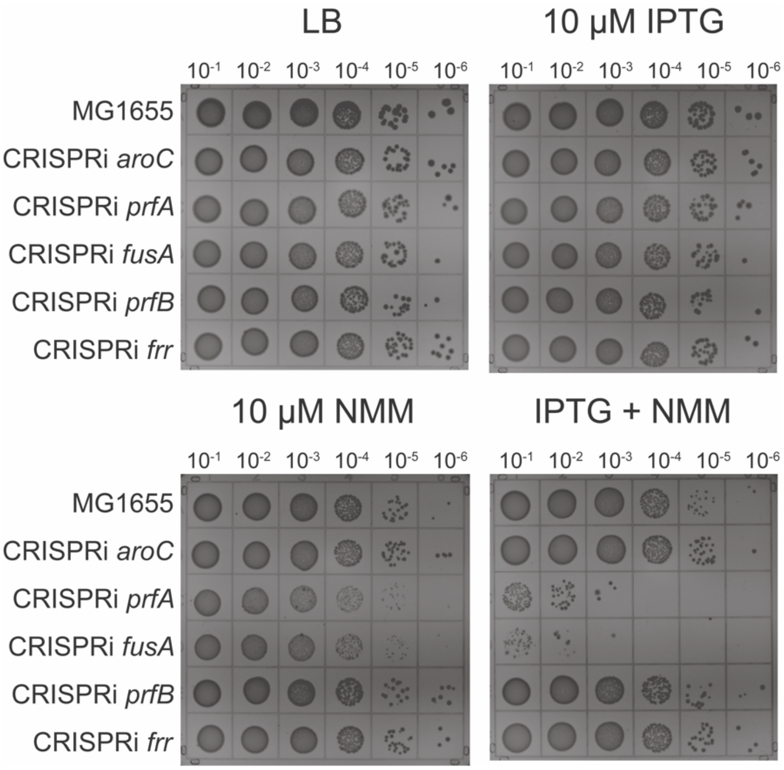


**Figure S4: CRISPR interference machinery targeting *prfA* and *fusA* are sensitive to NMM.** Spot dilution plates showing CRISPR interference machinery targeted to various genes. Spot plates contain LB-agar alone or LB-agar supplemented with IPTG, NMM, or both IPTG and NMM.

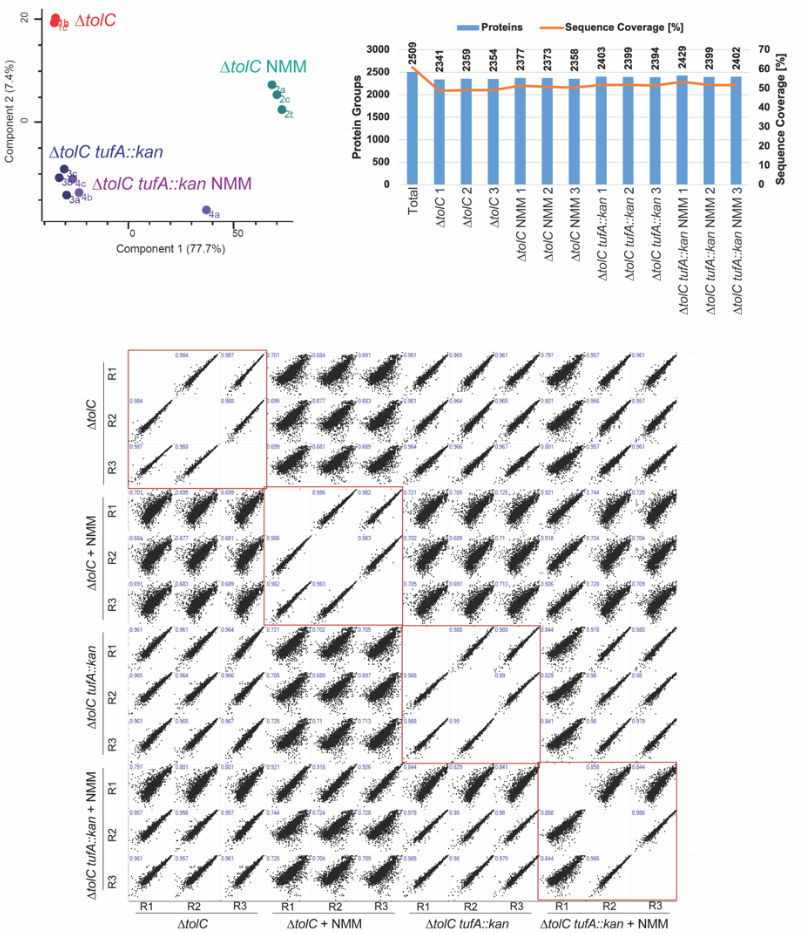


**Figure S5: Proteomics quality control.** Top left: principal component analysis of Δ*tolC,* Δ*tolC* NMM, Δ*tolC tufA::kan,* and Δ*tolC tufA::kan* NMM. Top right: total protein and sequence coverage for each replicate of the proteomics dataset. Bottom: Pearson correlation coefficients of each replicate/growth condition. Boxed in red are plots for correlation between replicates of each growth condition.

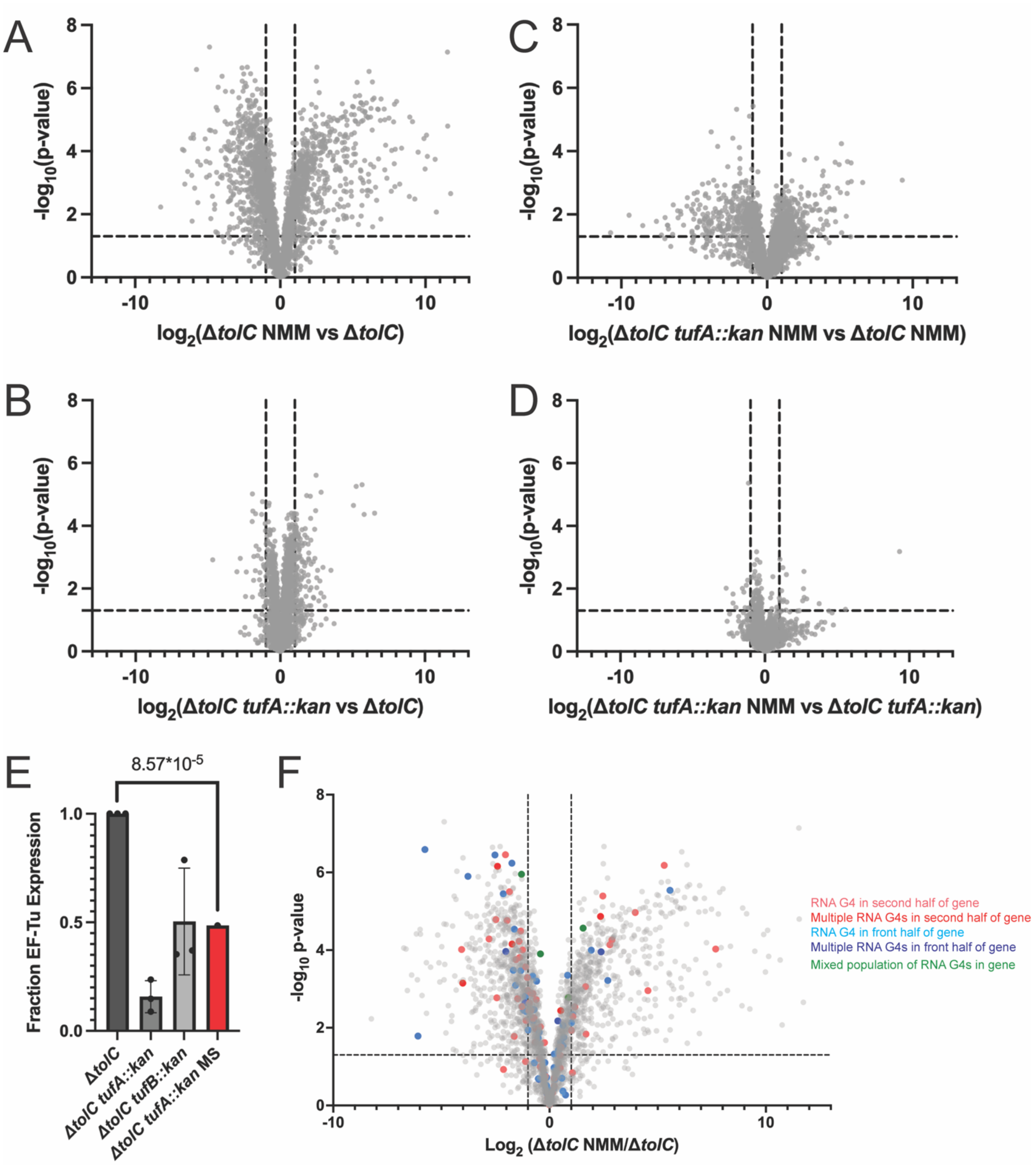


**Figure S6: Proteomics dataset contains largest differences in ∆*tolC* NMM vs ∆*tolC* cells. (A-D)**. Volcano plots comparing proteomic landscape of different growth conditions. Horizontal dashed line indicates p-value of 0.05, and vertical lines indicate 2-

fold change in protein detection. Each dot represents a different protein detected in the dataset. **(E)**. Decrease in EF-Tu levels detected via mass spectrometry shown in red compared to what was detected via western blotting. **(F).** Location of RNA G4s in ORF mapped onto $\Delta$*tolC* ± NMM Volcano plot.

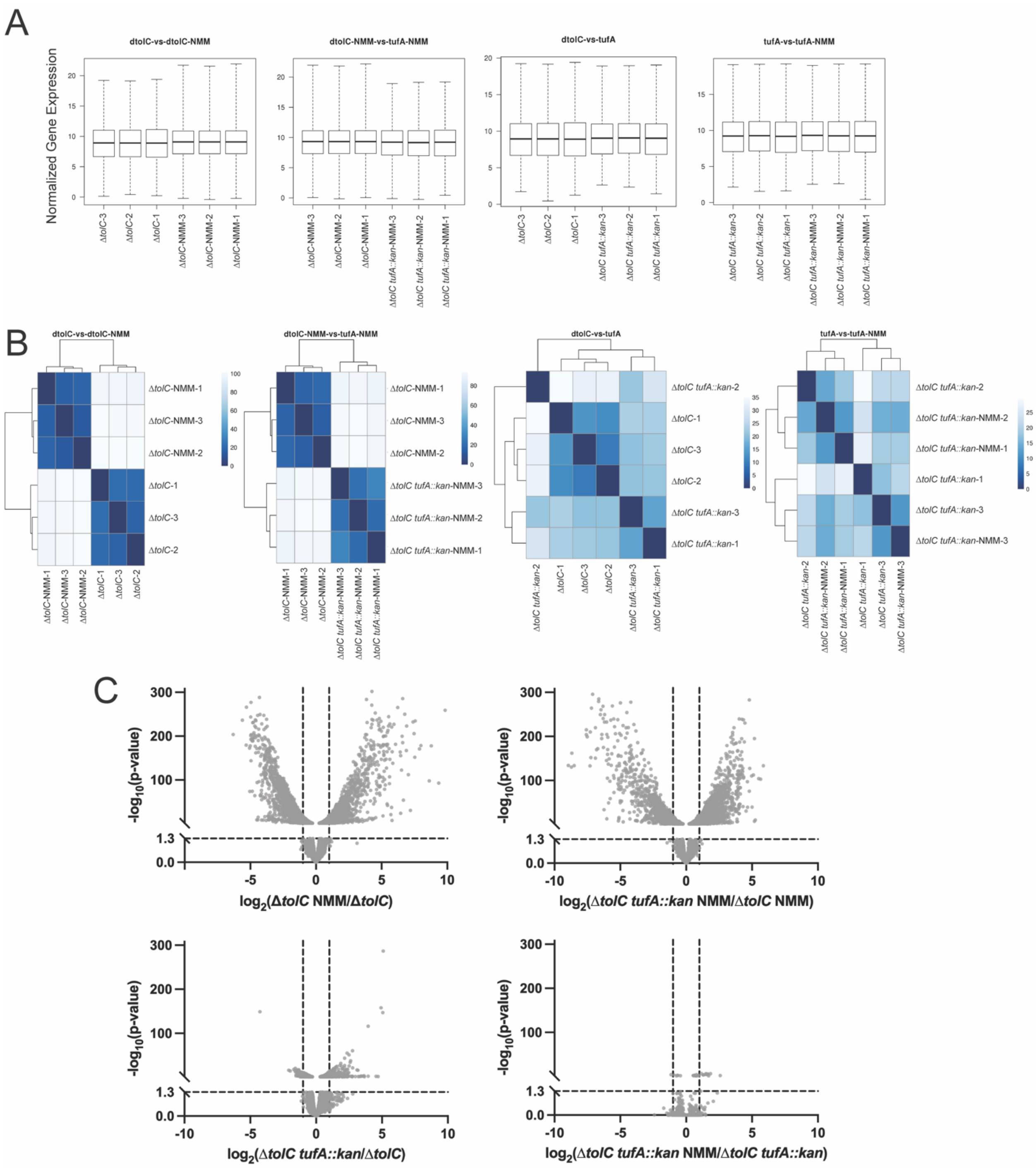


**Figure S7: RNA-seq results. (A).** Box and whisker plots of normalized gene expression for each growth condition. **(B)**. Sample distance plots generated using DeSeq2. Darker blue indicates closer correlation of expression values between samples. **(C).** Volcano plots of $log_2$(Fold Change between normalized reads) for each transcript. Each y-axis had a break at p-value 0.05 to indicate significance and contains vertical lines at a two-fold change in detection of the transcript.

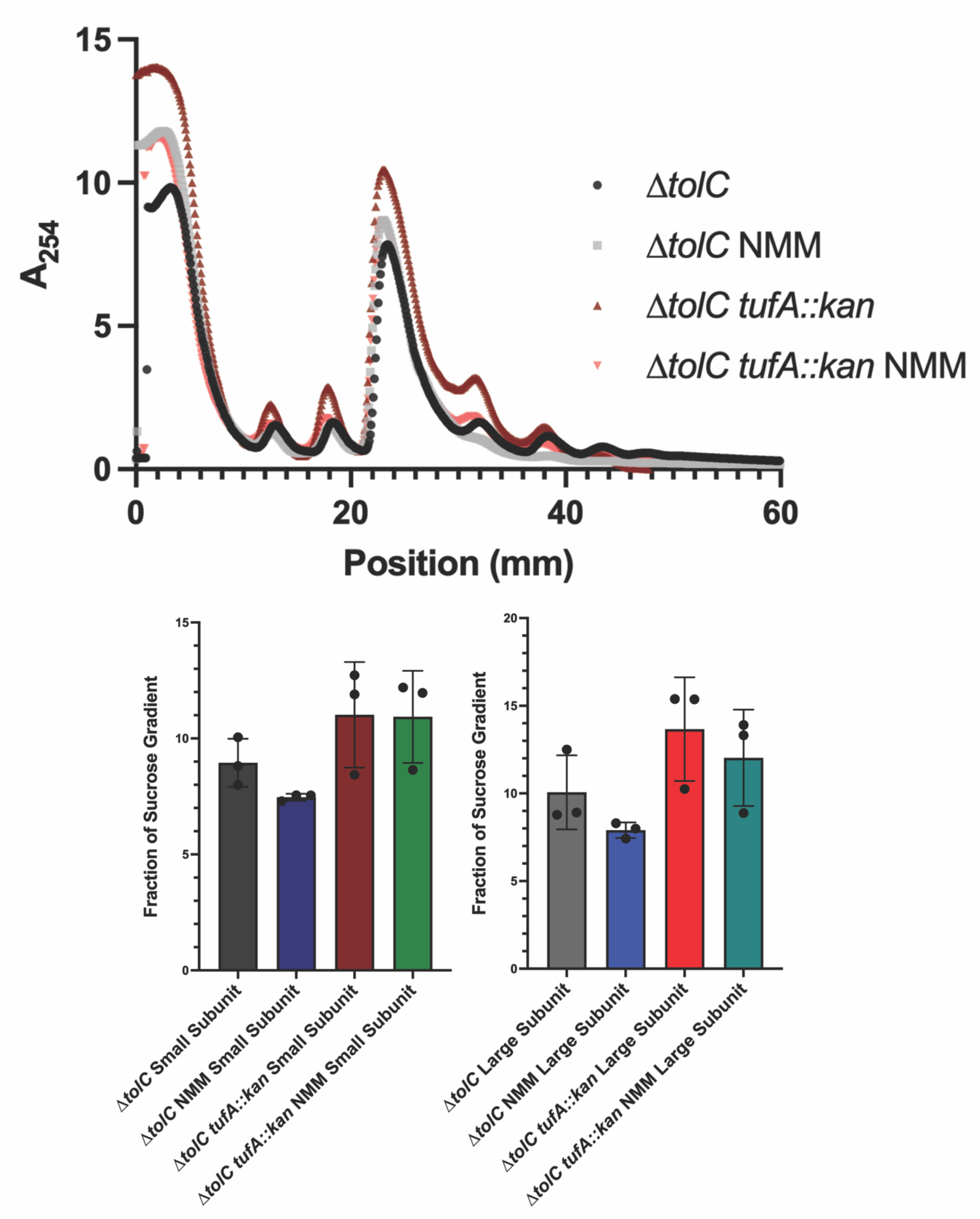

A254
Position (mm)
ΔtolC
ΔtolC NMM
ΔtolC tufA::kan
ΔtolC tufA::kan NMM
Fraction of Sucrose Gradient
ΔtolC Small Subunit
ΔtolC NMM Small Subunit
ΔtolC tufA::kan Small Subunit
ΔtolC tufA::kan NMM Small Subunit
ΔtolC Large Subunit
ΔtolC NMM Large Subunit
ΔtolC tufA::kan Large Subunit
ΔtolC tufA::kan NMM Large Subunit

**Figure S8**: **Sucrose gradients separating ribosome components reveals differences in ∆*tolC* NMM monosome and polysome abundance**. Top: Representative sucrose gradients of each growth condition. Bottom: quantification of area under the curve for small subunit and large subunit. P-values that were significant via Welch's t-test are displayed.

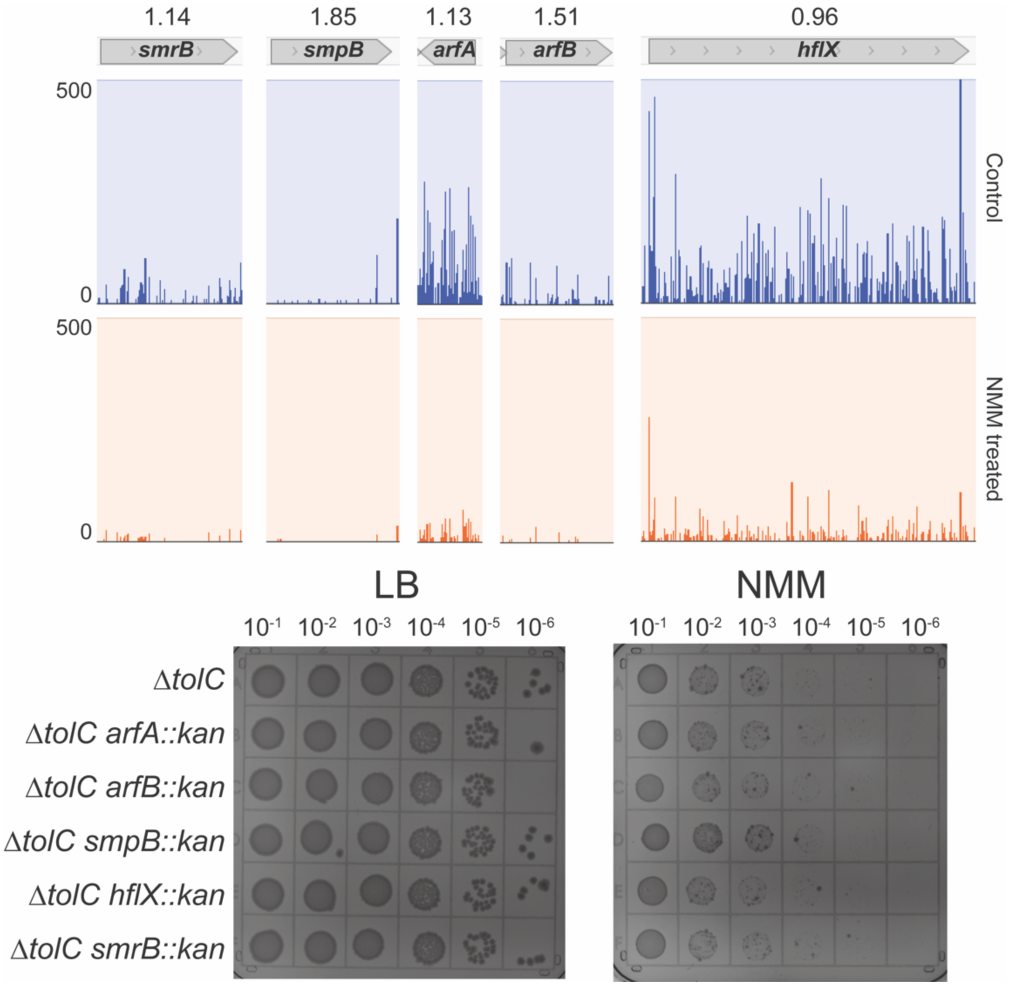


**Figure S9: Ribosome rescue factors in Tn-seq screen and spot plates.** Top: Mochiview plots of transposon insertions across various ribosome rescue factors. $Log_{10}$(ratio weighted reads) are above each Mochiview plot for each gene. Bottom: Spot plates of strains harboring ribosome rescue factor deletions in the presence and absence of NMM.

**Supplementary Tables**

**Table S1: Gene ontology analysis of Tn-seq hits for which insertions are negatively selected in G4 stabilizing conditions**

| Gene ontology biological process | Fold Enrichment | p-value (FDR correction) |
|---|---|---|
| heme transport (GO:0015886) | 2.56 | 1.40E-02 |
| molybdopterin cofactor metabolic process (GO:0043545) | 2.38 | 3.52E-02 |
| Mo-molybdopterin cofactor biosynthetic process (GO:0006777) | 2.38 | 3.45E-02 |
| molybdopterin cofactor biosynthetic process (GO:0032324) | 2.38 | 3.39E-02 |
| Mo-molybdopterin cofactor metabolic process (GO:0019720) | 2.38 | 3.32E-02 |
| iron ion transmembrane transport (GO:0034755) | 2.36 | 8.27E-04 |
| IMP biosynthetic process (GO:0006188) | 2.27 | 3.65E-02 |
| rRNA modification (GO:0000154) | 2.08 | 4.54E-03 |
| cellular response to oxidative stress (GO:0034599) | 1.89 | 4.10E-02 |
| iron ion transport (GO:0006826) | 1.87 | 2.90E-03 |
| monosaccharide transmembrane transport (GO:0015749) | 1.7 | 4.26E-02 |
| transition metal ion transport (GO:0000041) | 1.65 | 1.01E-02 |
| cellular response to chemical stress (GO:0062197) | 1.65 | 4.12E-02 |
| metal ion transport (GO:0030001) | 1.56 | 2.54E-03 |
| proteinogenic amino acid biosynthetic process (GO:0170038) | 1.53 | 3.60E-02 |
| L-amino acid biosynthetic process (GO:0170034) | 1.53 | 3.54E-02 |
| amino acid biosynthetic process (GO:0008652) | 1.46 | 1.31E-02 |

| | | |
|---|---|---|
| alpha-amino acid biosynthetic process (GO:1901607) | 1.45 | 3.36E-02 |
| monoatomic cation transmembrane transport (GO:0098655) | 1.42 | 3.64E-02 |
| inorganic cation transmembrane transport (GO:0098662) | 1.42 | 3.91E-02 |
| monoatomic cation transport (GO:0006812) | 1.4 | 1.31E-02 |
| monoatomic ion transmembrane transport (GO:0034220) | 1.38 | 3.42E-02 |
| DNA damage response (GO:0006974) | 1.37 | 2.67E-03 |
| monoatomic ion transport (GO:0006811) | 1.34 | 2.75E-02 |
| cellular response to stress (GO:0033554) | 1.33 | 7.10E-04 |
| regulation of RNA biosynthetic process (GO:2001141) | 1.32 | 2.89E-03 |
| regulation of DNA-templated transcription (GO:0006355) | 1.32 | 2.65E-03 |
| regulation of RNA metabolic process (GO:0051252) | 1.3 | 3.60E-03 |
| regulation of nucleobase-containing compound metabolic process (GO:0019219) | 1.27 | 8.09E-03 |
| response to stress (GO:0006950) | 1.24 | 2.79E-03 |
| cellular response to stimulus (GO:0051716) | 1.2 | 3.24E-02 |
| transport (GO:0006810) | 1.17 | 3.41E-02 |
| response to stimulus (GO:0050896) | 1.16 | 7.08E-03 |
| cellular component organization (GO:0016043) | 0.77 | 1.55E-02 |
| small molecule catabolic process (GO:0044282) | 0.71 | 9.88E-03 |
| generation of precursor metabolites and energy (GO:0006091) | 0.69 | 4.61E-02 |
| amide metabolic process (GO:0043603) | 0.67 | 2.72E-02 |
| lipid metabolic process (GO:0006629) | 0.64 | 1.22E-02 |
| monocarboxylic acid metabolic process (GO:0032787) | 0.62 | 2.79E-03 |
| organelle organization (GO:0006996) | 0.57 | 1.58E-02 |
| energy derivation by oxidation of organic compounds (GO:0015980) | 0.56 | 9.97E-03 |

| | | |
|---|---|---|
| peptide metabolic process (GO:0006518) | 0.55 | 6.45E-03 |
| electron transport chain (GO:0022900) | 0.54 | 1.57E-02 |
| amide biosynthetic process (GO:0043604) | 0.53 | 1.46E-03 |
| cellular lipid metabolic process (GO:0044255) | 0.49 | 3.29E-03 |
| peptide biosynthetic process (GO:0043043) | 0.44 | 5.78E-04 |
| cell motility (GO:0048870) | 0.43 | 3.69E-02 |
| cellular respiration (GO:0045333) | 0.39 | 3.29E-04 |
| anaerobic respiration (GO:0009061) | 0.36 | 9.87E-03 |
| fatty acid metabolic process (GO:0006631) | 0.36 | 7.23E-03 |
| non-membrane-bounded organelle assembly (GO:0140694) | 0.33 | 3.12E-03 |
| organelle assembly (GO:0070925) | 0.33 | 2.92E-03 |
| anaerobic electron transport chain (GO:0019645) | 0.33 | 3.61E-02 |
| ribosome assembly (GO:0042255) | 0.31 | 2.26E-02 |
| oligosaccharide metabolic process (GO:0009311) | 0.29 | 3.64E-02 |
| translation (GO:0006412) | 0.28 | 1.10E-06 |
| respiratory electron transport chain (GO:0022904) | 0.27 | 6.53E-04 |
| protein-RNA complex organization (GO:0071826) | 0.2 | 3.53E-03 |
| protein-RNA complex assembly (GO:0022618) | 0.2 | 3.37E-03 |
| transposition (GO:0032196) | 0.18 | 1.30E-04 |
| DNA transposition (GO:0006313) | 0.17 | 8.11E-04 |
| ribosomal small subunit biogenesis (GO:0042274) | 0.12 | 3.42E-02 |
| nitrogen cycle metabolic process (GO:0071941) | 0.1 | 1.04E-02 |
| cytoplasmic translation (GO:0002181) | < 0.01 | 4.45E-07 |
| reactive nitrogen species metabolic process (GO:2001057) | < 0.01 | 2.86E-03 |
| response to cold (GO:0009409) | < 0.01 | 3.77E-02 |
| nitrate assimilation (GO:0042128) | < 0.01 | 6.72E-03 |
| nitrate metabolic process (GO:0042126) | < 0.01 | 6.48E-03 |

| | | |
|---|---|---|
| formate oxidation (GO:0015944) | < 0.01 | 3.71E-02 |
| ribosomal small subunit assembly (GO:0000028) | < 0.01 | 2.94E-02 |

**Table S2: Gene ontology analysis of Tn-seq hits for which insertions are positively selected in G4 stabilizing conditions**

| Gene ontology biological process | Fold Enrichment | p-value (FDR correction) |
|---|---|---|
| canonical glycolysis (GO:0061621) | 35.49 | 2.05E-02 |
| glycolytic process through glucose-6-phosphate (GO:0061620) | 35.49 | 2.03E-02 |
| lipoate biosynthetic process (GO:0009107) | 35.49 | 2.01E-02 |
| lipoate metabolic process (GO:0009106) | 35.49 | 1.99E-02 |
| regulation of glycogen metabolic process (GO:0070873) | 35.49 | 1.98E-02 |
| acetate biosynthetic process (GO:0019413) | 35.49 | 1.96E-02 |
| regulation of generation of precursor metabolites and energy (GO:0043467) | 35.49 | 1.94E-02 |
| NADH regeneration (GO:0006735) | 35.49 | 1.93E-02 |
| nitrogen fixation (GO:0009399) | 35.49 | 1.91E-02 |
| negative regulation of carbohydrate metabolic process (GO:0045912) | 35.49 | 1.90E-02 |
| glucose catabolic process to pyruvate (GO:0061718) | 35.49 | 1.88E-02 |
| tRNA wobble position uridine thiolation (GO:0002143) | 30.42 | 6.42E-07 |

| | | |
|---|---|---|
| acetyl-CoA biosynthetic process (GO:0006085) | 26.62 | 2.78E-03 |
| glycolytic process through fructose-6-phosphate (GO:0061615) | 23.66 | 4.52E-02 |
| tRNA thio-modification (GO:0034227) | 23.66 | 6.43E-06 |
| lipid translocation (GO:0034204) | 23.66 | 4.49E-02 |
| regulation of membrane lipid distribution (GO:0097035) | 23.66 | 4.46E-02 |
| NAD catabolic process (GO:0019677) | 23.66 | 4.43E-02 |
| NADH metabolic process (GO:0006734) | 21.3 | 6.39E-03 |
| regulation of response to nutrient levels (GO:0032107) | 17.75 | 1.11E-02 |
| regulation of response to extracellular stimulus (GO:0032104) | 17.75 | 1.10E-02 |
| ribosomal small subunit assembly (GO:0000028) | 15.97 | 2.47E-07 |
| ribosomal small subunit biogenesis (GO:0042274) | 13.65 | 1.96E-07 |
| thioester biosynthetic process (GO:0035384) | 13.31 | 2.47E-02 |
| acyl-CoA biosynthetic process (GO:0071616) | 13.31 | 2.45E-02 |
| ribonucleoside diphosphate catabolic process (GO:0009191) | 13.08 | 5.15E-05 |
| purine ribonucleoside diphosphate catabolic process (GO:0009181) | 13.08 | 4.96E-05 |
| ADP catabolic process (GO:0046032) | 13.08 | 4.79E-05 |
| glycolytic process (GO:0006096) | 13.08 | 4.62E-05 |
| purine nucleoside diphosphate catabolic process (GO:0009137) | 12.42 | 5.92E-05 |

| | | |
|---|---|---|
| nucleoside diphosphate catabolic process (GO:0009134) | 12.42 | 5.75E-05 |
| pyridine nucleotide catabolic process (GO:0019364) | 12.42 | 5.59E-05 |
| ADP metabolic process (GO:0046031) | 12.42 | 5.44E-05 |
| purine ribonucleoside diphosphate metabolic process (GO:0009179) | 11.83 | 7.03E-05 |
| purine ribonucleotide catabolic process (GO:0009154) | 11.83 | 6.86E-05 |
| ubiquinone biosynthetic process (GO:0006744) | 11.83 | 6.71E-05 |
| ubiquinone metabolic process (GO:0006743) | 11.83 | 6.56E-05 |
| acetyl-CoA metabolic process (GO:0006084) | 11.83 | 3.38E-02 |
| pyridine-containing compound catabolic process (GO:0072526) | 11.29 | 8.81E-05 |
| purine nucleoside diphosphate metabolic process (GO:0009135) | 11.29 | 8.63E-05 |
| tRNA wobble uridine modification (GO:0002098) | 11.21 | 4.06E-04 |
| ribonucleotide catabolic process (GO:0009261) | 10.35 | 1.54E-04 |
| purine nucleotide catabolic process (GO:0006195) | 9.94 | 2.06E-04 |
| ribonucleoside diphosphate metabolic process (GO:0009185) | 9.56 | 2.47E-04 |
| quinone biosynthetic process (GO:1901663) | 8.87 | 4.11E-04 |
| quinone metabolic process (GO:1901661) | 8.87 | 4.05E-04 |
| ATP metabolic process (GO:0046034) | 8.87 | 3.98E-04 |

| | | |
|---|---|---|
| glyceraldehyde-3-phosphate metabolic process (GO:0019682) | 8.87 | 1.97E-02 |
| ketone biosynthetic process (GO:0042181) | 8.57 | 4.90E-04 |
| purine ribonucleoside triphosphate metabolic process (GO:0009205) | 8.28 | 5.97E-04 |
| tRNA wobble base modification (GO:0002097) | 8.19 | 2.16E-03 |
| nucleoside diphosphate metabolic process (GO:0009132) | 8.01 | 7.43E-04 |
| pyruvate metabolic process (GO:0006090) | 7.76 | 8.46E-04 |
| nicotinamide nucleotide metabolic process (GO:0046496) | 7.55 | 4.63E-05 |
| nucleotide catabolic process (GO:0009166) | 7.53 | 1.04E-03 |
| mRNA metabolic process (GO:0016071) | 7.47 | 3.44E-02 |
| ribonucleoside triphosphate metabolic process (GO:0009199) | 7.31 | 1.21E-03 |
| purine nucleoside triphosphate metabolic process (GO:0009144) | 7.31 | 1.20E-03 |
| nucleoside phosphate catabolic process (GO:1901292) | 7.31 | 1.18E-03 |
| pyridine nucleotide metabolic process (GO:0019362) | 7.24 | 5.53E-05 |
| ribosome assembly (GO:0042255) | 7.24 | 5.39E-05 |
| protein-RNA complex organization (GO:0071826) | 6.94 | 2.06E-04 |
| protein-RNA complex assembly (GO:0022618) | 6.94 | 2.02E-04 |
| negative regulation of translation (GO:0017148) | 6.83 | 1.75E-02 |
| purine-containing compound catabolic process (GO:0072523) | 6.6 | 8.14E-04 |

| | | |
|---|---|---|
| nucleoside triphosphate metabolic process (GO:0009141) | 6.6 | 8.03E-04 |
| negative regulation of amide metabolic process (GO:0034249) | 6.57 | 1.90E-02 |
| pyridine-containing compound metabolic process (GO:0072524) | 6.51 | 5.43E-05 |
| negative regulation of gene expression (GO:0010629) | 6.14 | 4.97E-04 |
| nitrogen cycle metabolic process (GO:0071941) | 5.92 | 2.89E-02 |
| negative regulation of protein metabolic process (GO:0051248) | 5.72 | 3.34E-02 |
| positive regulation of gene expression (GO:0010628) | 5.72 | 3.32E-02 |
| non-membrane-bounded organelle assembly (GO:0140694) | 5.35 | 2.35E-04 |
| organelle assembly (GO:0070925) | 5.35 | 2.31E-04 |
| nucleobase-containing compound catabolic process (GO:0034655) | 5.32 | 1.01E-04 |
| ribosome biogenesis (GO:0042254) | 5.27 | 1.51E-05 |
| ribonucleoprotein complex biogenesis (GO:0022613) | 5.22 | 1.56E-05 |
| cellular ketone metabolic process (GO:0042180) | 5.18 | 9.27E-03 |
| sulfur compound biosynthetic process (GO:0044272) | 5.07 | 8.20E-04 |
| cytoplasmic translation (GO:0002181) | 4.98 | 5.23E-03 |
| post-transcriptional regulation of gene expression (GO:0010608) | 4.91 | 2.35E-03 |
| tRNA processing (GO:0008033) | 4.86 | 1.12E-03 |
| regulation of translation (GO:0006417) | 4.81 | 6.35E-03 |
| tRNA modification (GO:0006400) | 4.81 | 6.29E-03 |

| | | |
|---|---|---|
| regulation of amide metabolic process (GO:0034248) | 4.73 | 6.88E-03 |
| translation (GO:0006412) | 4.73 | 2.22E-05 |
| peptide biosynthetic process (GO:0043043) | 4.57 | 1.63E-05 |
| organophosphate catabolic process (GO:0046434) | 4.44 | 2.03E-02 |
| tRNA metabolic process (GO:0006399) | 4.26 | 7.74E-04 |
| amide biosynthetic process (GO:0043604) | 4.25 | 4.84E-06 |
| purine nucleotide metabolic process (GO:0006163) | 4.12 | 5.22E-04 |
| purine ribonucleotide metabolic process (GO:0009150) | 4.1 | 8.42E-03 |
| peptide metabolic process (GO:0006518) | 4.08 | 5.40E-05 |
| RNA processing (GO:0006396) | 3.91 | 8.28E-04 |
| ncRNA processing (GO:0034470) | 3.91 | 1.56E-03 |
| regulation of protein metabolic process (GO:0051246) | 3.84 | 2.29E-02 |
| organelle organization (GO:0006996) | 3.83 | 2.96E-04 |
| purine-containing compound metabolic process (GO:0072521) | 3.65 | 8.47E-04 |
| ribose phosphate metabolic process (GO:0019693) | 3.61 | 5.97E-03 |
| ncRNA metabolic process (GO:0034660) | 3.52 | 1.21E-03 |
| RNA modification (GO:0009451) | 3.47 | 2.29E-02 |
| sulfur compound metabolic process (GO:0006790) | 3.42 | 8.82E-03 |
| ribonucleotide metabolic process (GO:0009259) | 3.36 | 2.82E-02 |
| amide metabolic process (GO:0043603) | 3.33 | 8.86E-05 |

| | | |
|---|---|---|
| heterocycle catabolic process (GO:0046700) | 3.33 | 6.44E-03 |
| protein-containing complex assembly (GO:0065003) | 3.33 | 1.23E-03 |
| carbohydrate derivative catabolic process (GO:1901136) | 3.3 | 5.00E-02 |
| cellular nitrogen compound catabolic process (GO:0044270) | 3.3 | 6.86E-03 |
| gene expression (GO:0010467) | 3.27 | 9.52E-09 |
| protein-containing complex organization (GO:0043933) | 3.1 | 2.60E-03 |
| RNA metabolic process (GO:0016070) | 3.08 | 6.18E-05 |
| nucleotide metabolic process (GO:0009117) | 3.01 | 5.82E-03 |
| aromatic compound catabolic process (GO:0019439) | 2.98 | 1.62E-02 |
| nucleoside phosphate metabolic process (GO:0006753) | 2.94 | 6.91E-03 |
| small molecule biosynthetic process (GO:0044283) | 2.86 | 6.67E-05 |
| macromolecule modification (GO:0043412) | 2.86 | 2.02E-02 |
| macromolecule biosynthetic process (GO:0009059) | 2.84 | 3.57E-10 |
| positive regulation of cellular metabolic process (GO:0031325) | 2.79 | 3.62E-02 |
| positive regulation of macromolecule metabolic process (GO:0010604) | 2.71 | 4.41E-02 |
| organic cyclic compound catabolic process (GO:1901361) | 2.66 | 3.40E-02 |
| generation of precursor metabolites and energy (GO:0006091) | 2.57 | 1.62E-02 |
| cellular biosynthetic process (GO:0044249) | 2.51 | 4.41E-15 |

| | | |
|---|---|---|
| organic substance biosynthetic process (GO:1901576) | 2.51 | 2.77E-15 |
| protein metabolic process (GO:0019538) | 2.49 | 5.53E-04 |
| negative regulation of metabolic process (GO:0009892) | 2.49 | 2.03E-02 |
| biosynthetic process (GO:0009058) | 2.48 | 2.96E-15 |
| cellular nitrogen compound biosynthetic process (GO:0044271) | 2.44 | 5.33E-05 |
| negative regulation of cellular metabolic process (GO:0031324) | 2.38 | 4.05E-02 |
| negative regulation of macromolecule metabolic process (GO:0010605) | 2.34 | 4.51E-02 |
| nucleobase-containing small molecule metabolic process (GO:0055086) | 2.3 | 4.12E-02 |
| organonitrogen compound biosynthetic process (GO:1901566) | 2.26 | 1.28E-04 |
| cellular component biogenesis (GO:0044085) | 2.24 | 5.81E-03 |
| organophosphate metabolic process (GO:0019637) | 2.16 | 3.32E-02 |
| cellular nitrogen compound metabolic process (GO:0034641) | 2.12 | 1.97E-07 |
| phosphate-containing compound metabolic process (GO:0006796) | 2.07 | 9.02E-03 |
| phosphorus metabolic process (GO:0006793) | 2.04 | 1.11E-02 |
| cellular component organization or biogenesis (GO:0071840) | 2.01 | 3.99E-03 |
| nucleobase-containing compound metabolic process (GO:0006139) | 2 | 2.09E-04 |

| | | |
|---|---|---|
| organic cyclic compound biosynthetic process (GO:1901362) | 2 | 3.39E-02 |
| heterocycle biosynthetic process (GO:0018130) | 1.99 | 4.42E-02 |
| heterocycle metabolic process (GO:0046483) | 1.95 | 6.58E-05 |
| macromolecule metabolic process (GO:0043170) | 1.94 | 1.55E-05 |
| organonitrogen compound metabolic process (GO:1901564) | 1.94 | 2.55E-05 |
| carbohydrate derivative metabolic process (GO:1901135) | 1.93 | 4.47E-02 |
| organic cyclic compound metabolic process (GO:1901360) | 1.89 | 9.88E-05 |
| nitrogen compound metabolic process (GO:0006807) | 1.84 | 1.98E-08 |
| regulation of metabolic process (GO:0019222) | 1.84 | 4.01E-02 |
| nucleic acid metabolic process (GO:0090304) | 1.82 | 4.41E-02 |
| cellular aromatic compound metabolic process (GO:0006725) | 1.76 | 1.89E-03 |
| cellular metabolic process (GO:0044237) | 1.74 | 6.38E-11 |
| primary metabolic process (GO:0044238) | 1.73 | 1.14E-07 |
| biological regulation (GO:0065007) | 1.69 | 3.41E-02 |
| metabolic process (GO:0008152) | 1.65 | 6.49E-11 |
| organic substance metabolic process (GO:0071704) | 1.65 | 1.66E-08 |
| small molecule metabolic process (GO:0044281) | 1.6 | 2.89E-02 |
| cellular process (GO:0009987) | 1.31 | 2.62E-05 |
| biological_process (GO:0008150) | 1.23 | 2.58E-05 |
| Unclassified (UNCLASSIFIED) | 0.24 | 2.48E-05 |

**Table S3: Gene ontology analysis of proteins less abundant in G4 stabilizing conditions**

| Gene ontology biological process | Fold Enrichment | p-value (FDR correction) |
|---|---|---|
| ribosomal large subunit assembly (GO:0000027) | 6.47 | 5.15E-08 |
| ribosomal large subunit biogenesis (GO:0042273) | 6.22 | 8.54E-08 |
| cytoplasmic translation (GO:0002181) | 6.03 | 1.84E-16 |
| protein-RNA complex organization (GO:0071826) | 6.01 | 4.14E-13 |
| protein-RNA complex assembly (GO:0022618) | 6.01 | 4.00E-13 |
| ribosome assembly (GO:0042255) | 5.91 | 1.35E-13 |
| ribosomal small subunit assembly (GO:0000028) | 5.73 | 4.05E-05 |
| bacteriocin transport (GO:0043213) | 5.39 | 2.57E-02 |
| regulation of DNA-templated transcription elongation (GO:0032784) | 4.9 | 3.65E-02 |
| ribosomal small subunit biogenesis (GO:0042274) | 4.67 | 1.44E-04 |
| ribosome biogenesis (GO:0042254) | 4.54 | 1.81E-17 |
| ribonucleoprotein complex biogenesis (GO:0022613) | 4.49 | 2.52E-17 |
| amino acid activation (GO:0043038) | 4.41 | 4.37E-04 |
| non-membrane-bounded organelle assembly (GO:0140694) | 4.34 | 1.76E-11 |
| organelle assembly (GO:0070925) | 4.34 | 1.71E-11 |
| translation (GO:0006412) | 4.32 | 6.99E-19 |
| tRNA aminoacylation for protein translation (GO:0006418) | 4.31 | 8.91E-04 |
| lipopolysaccharide core region metabolic process (GO:0046401) | 4.31 | 8.80E-04 |
| lipopolysaccharide core region biosynthetic process (GO:0009244) | 4.31 | 8.68E-04 |
| tRNA aminoacylation (GO:0043039) | 4.31 | 8.57E-04 |
| peptide biosynthetic process (GO:0043043) | 4.14 | 5.10E-19 |

| | | |
|---|---|---|
| DNA conformation change (GO:0071103) | 4.13 | 2.71E-04 |
| DNA-templated DNA replication maintenance of fidelity (GO:0045005) | 4.04 | 4.75E-02 |
| peptide metabolic process (GO:0006518) | 3.78 | 7.45E-18 |
| oligosaccharide biosynthetic process (GO:0009312) | 3.72 | 2.82E-03 |
| DNA duplex unwinding (GO:0032508) | 3.65 | 1.26E-02 |
| DNA geometric change (GO:0032392) | 3.65 | 1.25E-02 |
| amide biosynthetic process (GO:0043604) | 3.59 | 3.92E-18 |
| rRNA modification (GO:0000154) | 3.58 | 2.46E-03 |
| tRNA modification (GO:0006400) | 3.54 | 5.44E-06 |
| organelle organization (GO:0006996) | 3.54 | 1.61E-14 |
| rRNA methylation (GO:0031167) | 3.52 | 2.51E-02 |
| tRNA metabolic process (GO:0006399) | 3.5 | 4.15E-10 |
| DNA-templated DNA replication (GO:0006261) | 3.46 | 1.38E-03 |
| RNA modification (GO:0009451) | 3.44 | 6.31E-09 |
| tRNA processing (GO:0008033) | 3.42 | 7.08E-07 |
| RNA methylation (GO:0001510) | 3.37 | 1.22E-03 |
| negative regulation of translation (GO:0017148) | 3.37 | 2.17E-02 |
| DNA replication (GO:0006260) | 3.31 | 3.90E-05 |
| ncRNA metabolic process (GO:0034660) | 3.3 | 1.45E-12 |
| rRNA processing (GO:0006364) | 3.29 | 7.51E-04 |
| ncRNA processing (GO:0034470) | 3.28 | 1.75E-09 |
| negative regulation of amide metabolic process (GO:0034249) | 3.25 | 2.71E-02 |
| chromosome organization (GO:0051276) | 3.24 | 4.16E-04 |
| RNA processing (GO:0006396) | 3.2 | 9.95E-10 |
| macromolecule methylation (GO:0043414) | 3.16 | 7.86E-04 |
| rRNA metabolic process (GO:0016072) | 3.15 | 1.14E-03 |
| tRNA wobble base modification (GO:0002097) | 3.11 | 4.95E-02 |
| lipid localization (GO:0010876) | 3.1 | 8.49E-03 |
| lipid transport (GO:0006869) | 3.1 | 8.40E-03 |
| positive regulation of gene expression (GO:0010628) | 3.04 | 2.81E-02 |
| amide metabolic process (GO:0043603) | 2.94 | 9.88E-15 |

| | | |
|---|---|---|
| establishment of localization in cell (GO:0051649) | 2.63 | 4.81E-02 |
| gene expression (GO:0010467) | 2.62 | 1.14E-21 |
| cell cycle process (GO:0022402) | 2.6 | 8.76E-03 |
| macromolecule modification (GO:0043412) | 2.58 | 4.39E-07 |
| methylation (GO:0032259) | 2.53 | 8.46E-03 |
| macromolecule biosynthetic process (GO:0009059) | 2.5 | 1.77E-31 |
| RNA metabolic process (GO:0016070) | 2.49 | 5.65E-12 |
| negative regulation of gene expression (GO:0010629) | 2.46 | 4.30E-02 |
| lipopolysaccharide biosynthetic process (GO:0009103) | 2.43 | 6.52E-03 |
| response to radiation (GO:0009314) | 2.43 | 7.46E-04 |
| cell cycle (GO:0007049) | 2.42 | 1.62E-03 |
| regulation of translation (GO:0006417) | 2.4 | 2.82E-02 |
| regulation of amide metabolic process (GO:0034248) | 2.36 | 3.08E-02 |
| cellular nitrogen compound biosynthetic process (GO:0044271) | 2.36 | 1.64E-19 |
| cell division (GO:0051301) | 2.35 | 5.12E-03 |
| lipopolysaccharide metabolic process (GO:0008653) | 2.34 | 1.20E-02 |
| protein-containing complex organization (GO:0043933) | 2.31 | 5.42E-06 |
| response to antibiotic (GO:0046677) | 2.29 | 1.98E-03 |
| post-transcriptional regulation of gene expression (GO:0010608) | 2.28 | 4.10E-02 |
| protein-containing complex assembly (GO:0065003) | 2.27 | 3.06E-05 |
| polysaccharide biosynthetic process (GO:0000271) | 2.27 | 1.21E-03 |
| carbohydrate derivative biosynthetic process (GO:1901137) | 2.24 | 3.05E-08 |
| protein metabolic process (GO:0019538) | 2.22 | 1.79E-11 |
| organonitrogen compound biosynthetic process (GO:1901566) | 2.19 | 1.20E-17 |
| cellular component biogenesis (GO:0044085) | 2.16 | 5.01E-11 |
| liposaccharide metabolic process (GO:1903509) | 2.14 | 2.49E-02 |

| | | |
|---|---|---|
| polysaccharide metabolic process (GO:0005976) | 2.1 | 3.81E-03 |
| cellular biosynthetic process (GO:0044249) | 2.04 | 3.07E-33 |
| nucleotide biosynthetic process (GO:0009165) | 2.04 | 2.97E-02 |
| nucleoside phosphate biosynthetic process (GO:1901293) | 2.04 | 2.94E-02 |
| cellular component organization or biogenesis (GO:0071840) | 2.03 | 1.38E-13 |
| regulation of biological quality (GO:0065008) | 2.03 | 2.50E-02 |
| organic substance biosynthetic process (GO:1901576) | 2.02 | 1.69E-33 |
| biosynthetic process (GO:0009058) | 1.99 | 7.70E-33 |
| lipid biosynthetic process (GO:0008610) | 1.99 | 5.60E-03 |
| nucleic acid metabolic process (GO:0090304) | 1.98 | 7.17E-12 |
| macromolecule metabolic process (GO:0043170) | 1.98 | 2.87E-28 |
| carbohydrate biosynthetic process (GO:0016051) | 1.96 | 1.25E-02 |
| nucleobase-containing compound biosynthetic process (GO:0034654) | 1.95 | 5.29E-05 |
| macromolecule localization (GO:0033036) | 1.93 | 8.81E-03 |
| cellular component assembly (GO:0022607) | 1.92 | 4.39E-05 |
| cellular component organization (GO:0016043) | 1.91 | 1.75E-09 |
| cellular nitrogen compound metabolic process (GO:0034641) | 1.88 | 5.58E-22 |
| nucleobase-containing compound metabolic process (GO:0006139) | 1.78 | 2.34E-12 |
| organophosphate biosynthetic process (GO:0090407) | 1.76 | 1.18E-02 |
| aromatic compound biosynthetic process (GO:0019438) | 1.76 | 1.40E-04 |
| response to abiotic stimulus (GO:0009628) | 1.68 | 5.44E-03 |
| organic cyclic compound biosynthetic process (GO:1901362) | 1.67 | 4.85E-04 |
| nitrogen compound transport (GO:0071705) | 1.67 | 6.30E-03 |
| heterocycle biosynthetic process (GO:0018130) | 1.66 | 8.66E-04 |
| heterocycle metabolic process (GO:0046483) | 1.63 | 2.97E-10 |

| | | |
|---|---|---|
| carbohydrate derivative metabolic process (GO:1901135) | 1.6 | 1.14E-03 |
| cellular aromatic compound metabolic process (GO:0006725) | 1.6 | 1.87E-09 |
| organic cyclic compound metabolic process (GO:1901360) | 1.57 | 3.31E-09 |
| nitrogen compound metabolic process (GO:0006807) | 1.55 | 2.73E-17 |
| primary metabolic process (GO:0044238) | 1.49 | 1.47E-16 |
| organonitrogen compound metabolic process (GO:1901564) | 1.47 | 7.98E-07 |
| cellular metabolic process (GO:0044237) | 1.42 | 2.52E-16 |
| organic substance metabolic process (GO:0071704) | 1.38 | 7.79E-14 |
| metabolic process (GO:0008152) | 1.32 | 2.24E-12 |
| biological regulation (GO:0065007) | 1.31 | 4.98E-02 |
| cellular process (GO:0009987) | 1.28 | 3.40E-22 |
| biological_process (GO:0008150) | 1.19 | 1.15E-18 |
| cellular catabolic process (GO:0044248) | 0.48 | 1.32E-03 |
| monocarboxylic acid metabolic process (GO:0032787) | 0.48 | 3.57E-02 |
| organonitrogen compound catabolic process (GO:1901565) | 0.48 | 2.82E-02 |
| organic substance catabolic process (GO:1901575) | 0.47 | 2.04E-05 |
| catabolic process (GO:0009056) | 0.45 | 3.77E-06 |
| carboxylic acid catabolic process (GO:0046395) | 0.37 | 6.74E-03 |
| Unclassified (UNCLASSIFIED) | 0.36 | 1.06E-18 |
| organic acid catabolic process (GO:0016054) | 0.35 | 2.86E-03 |
| small molecule catabolic process (GO:0044282) | 0.29 | 1.77E-06 |
| carbohydrate catabolic process (GO:0016052) | 0.2 | 1.31E-03 |
| monosaccharide metabolic process (GO:0005996) | 0.12 | 1.13E-03 |
| carbohydrate transport (GO:0008643) | 0.1 | 1.87E-04 |
| transposition (GO:0032196) | 0.1 | 3.73E-02 |
| hexose metabolic process (GO:0019318) | 0.09 | 1.29E-02 |
| carbohydrate transmembrane transport (GO:0034219) | 0.06 | 2.32E-04 |

**Table S4: Gene ontology analysis of proteins more abundant in G4 stabilizing conditions**

| Gene ontology biological process | Fold Enrichment | p-value (FDR correction) |
|---|---|---|
| glycogen metabolic process (GO:0005977) | 8.08 | 1.32E-02 |
| energy reserve metabolic process (GO:0006112) | 8.08 | 1.30E-02 |
| N-acetylneuraminate catabolic process (GO:0019262) | 6.46 | 1.30E-02 |
| alditol catabolic process (GO:0019405) | 6.28 | 3.88E-04 |
| ketone catabolic process (GO:0042182) | 6.21 | 4.17E-03 |
| glycerol catabolic process (GO:0019563) | 6.06 | 9.76E-03 |
| polyol catabolic process (GO:0046174) | 6.06 | 2.61E-04 |
| methylglyoxal metabolic process (GO:0009438) | 5.87 | 1.78E-02 |
| N-acetylneuraminate metabolic process (GO:0006054) | 5.87 | 1.76E-02 |
| methylglyoxal catabolic process (GO:0051596) | 5.65 | 3.47E-02 |
| polysaccharide catabolic process (GO:0000272) | 5.38 | 2.28E-02 |
| hexitol metabolic process (GO:0006059) | 5.14 | 4.56E-02 |
| oligosaccharide catabolic process (GO:0009313) | 5.14 | 4.53E-02 |
| dicarboxylic acid catabolic process (GO:0043649) | 4.97 | 3.04E-02 |
| disaccharide metabolic process (GO:0005984) | 4.85 | 1.99E-02 |
| amino sugar catabolic process (GO:0046348) | 4.85 | 1.97E-02 |
| ribonucleoside diphosphate catabolic process (GO:0009191) | 4.68 | 9.96E-03 |

| | | |
|---|---|---|
| purine ribonucleoside diphosphate catabolic process (GO:0009181) | 4.68 | 9.80E-03 |
| aldehyde catabolic process (GO:0046185) | 4.68 | 9.64E-03 |
| ADP catabolic process (GO:0046032) | 4.68 | 9.49E-03 |
| glycolytic process (GO:0006096) | 4.68 | 9.34E-03 |
| purine nucleoside diphosphate catabolic process (GO:0009137) | 4.44 | 1.23E-02 |
| nucleoside diphosphate catabolic process (GO:0009134) | 4.44 | 1.21E-02 |
| pyridine nucleotide catabolic process (GO:0019364) | 4.44 | 1.19E-02 |
| ADP metabolic process (GO:0046031) | 4.44 | 1.18E-02 |
| purine ribonucleoside diphosphate metabolic process (GO:0009179) | 4.23 | 1.49E-02 |
| purine ribonucleotide catabolic process (GO:0009154) | 4.23 | 1.47E-02 |
| nucleotide catabolic process (GO:0009166) | 4.16 | 1.45E-03 |
| alditol metabolic process (GO:0019400) | 4.16 | 1.40E-03 |
| pyridine-containing compound catabolic process (GO:0072526) | 4.04 | 1.75E-02 |
| purine nucleoside diphosphate metabolic process (GO:0009135) | 4.04 | 1.73E-02 |
| pyruvate metabolic process (GO:0006090) | 4.04 | 2.02E-03 |
| nucleoside phosphate catabolic process (GO:1901292) | 4.04 | 1.44E-03 |
| polyol metabolic process (GO:0019751) | 3.93 | 1.37E-03 |
| purine nucleotide catabolic process (GO:0006195) | 3.88 | 1.45E-02 |
| ribonucleotide catabolic process (GO:0009261) | 3.7 | 2.60E-02 |
| glycerol metabolic process (GO:0006071) | 3.67 | 3.93E-02 |
| cellular oxidant detoxification (GO:0098869) | 3.51 | 4.72E-02 |

| | | |
|---|---|---|
| carbohydrate catabolic process (GO:0016052) | 3.51 | 3.30E-11 |
| carbohydrate derivative catabolic process (GO:1901136) | 3.47 | 1.66E-06 |
| ribonucleoside diphosphate metabolic process (GO:0009185) | 3.42 | 3.77E-02 |
| organophosphate catabolic process (GO:0046434) | 3.32 | 1.05E-03 |
| alcohol catabolic process (GO:0046164) | 3.16 | 6.72E-03 |
| cellular response to oxidative stress (GO:0034599) | 3.03 | 4.76E-02 |
| nicotinamide nucleotide metabolic process (GO:0046496) | 2.92 | 1.47E-02 |
| response to reactive oxygen species (GO:0000302) | 2.88 | 2.63E-02 |
| pyridine nucleotide metabolic process (GO:0019362) | 2.8 | 1.86E-02 |
| organic hydroxy compound catabolic process (GO:1901616) | 2.8 | 2.04E-02 |
| cellular response to chemical stress (GO:0062197) | 2.55 | 3.03E-02 |
| response to oxidative stress (GO:0006979) | 2.53 | 1.50E-03 |
| response to oxygen-containing compound (GO:1901700) | 2.35 | 1.80E-02 |
| monosaccharide metabolic process (GO:0005996) | 2.34 | 2.90E-03 |
| amino acid catabolic process (GO:0009063) | 2.32 | 2.12E-02 |
| hexose metabolic process (GO:0019318) | 2.31 | 2.74E-02 |
| small molecule catabolic process (GO:0044282) | 2.26 | 1.67E-08 |
| protein complex oligomerization (GO:0051259) | 2.25 | 4.15E-02 |
| organic substance catabolic process (GO:1901575) | 2.12 | 3.45E-12 |

| | | |
|---|---|---|
| catabolic process (GO:0009056) | 2.12 | 2.23E-12 |
| organonitrogen compound catabolic process (GO:1901565) | 2.11 | 7.97E-05 |
| carbohydrate transmembrane transport (GO:0034219) | 2.11 | 2.37E-02 |
| carbohydrate metabolic process (GO:0005975) | 2.09 | 4.72E-08 |
| cellular catabolic process (GO:0044248) | 2.04 | 3.75E-07 |
| carboxylic acid catabolic process (GO:0046395) | 2.01 | 1.49E-03 |
| carbohydrate transport (GO:0008643) | 1.94 | 4.15E-02 |
| organic hydroxy compound metabolic process (GO:1901615) | 1.93 | 4.23E-02 |
| organic acid catabolic process (GO:0016054) | 1.9 | 3.85E-03 |
| cellular response to stress (GO:0033554) | 1.89 | 8.57E-06 |
| DNA damage response (GO:0006974) | 1.81 | 2.09E-03 |
| monocarboxylic acid metabolic process (GO:0032787) | 1.73 | 1.62E-02 |
| response to stress (GO:0006950) | 1.7 | 1.56E-05 |
| oxoacid metabolic process (GO:0043436) | 1.59 | 1.11E-03 |
| carboxylic acid metabolic process (GO:0019752) | 1.58 | 1.50E-03 |
| cellular response to stimulus (GO:0051716) | 1.56 | 1.42E-03 |
| organic acid metabolic process (GO:0006082) | 1.55 | 1.92E-03 |
| small molecule metabolic process (GO:0044281) | 1.49 | 8.12E-05 |
| response to stimulus (GO:0050896) | 1.27 | 3.79E-02 |
| establishment of localization (GO:0051234) | 0.69 | 3.94E-02 |
| localization (GO:0051179) | 0.69 | 3.43E-02 |
| biosynthetic process (GO:0009058) | 0.68 | 2.12E-03 |

| | | |
|---|---|---|
| organic substance biosynthetic process (GO:1901576) | 0.67 | 1.44E-03 |
| organic cyclic compound metabolic process (GO:1901360) | 0.66 | 3.35E-03 |
| heterocycle metabolic process (GO:0046483) | 0.65 | 3.10E-03 |
| regulation of cellular process (GO:0050794) | 0.64 | 3.05E-02 |
| macromolecule metabolic process (GO:0043170) | 0.63 | 4.38E-04 |
| cellular aromatic compound metabolic process (GO:0006725) | 0.62 | 1.14E-03 |
| cellular biosynthetic process (GO:0044249) | 0.61 | 8.47E-05 |
| regulation of primary metabolic process (GO:0080090) | 0.6 | 4.15E-02 |
| nucleobase-containing compound metabolic process (GO:0006139) | 0.6 | 1.86E-03 |
| regulation of nitrogen compound metabolic process (GO:0051171) | 0.59 | 3.94E-02 |
| regulation of metabolic process (GO:0019222) | 0.59 | 2.32E-02 |
| cellular nitrogen compound metabolic process (GO:0034641) | 0.57 | 3.41E-05 |
| regulation of nucleobase-containing compound metabolic process (GO:0019219) | 0.57 | 4.39E-02 |
| regulation of cellular metabolic process (GO:0031323) | 0.57 | 1.79E-02 |
| regulation of macromolecule metabolic process (GO:0060255) | 0.57 | 1.74E-02 |
| regulation of RNA metabolic process (GO:0051252) | 0.55 | 3.14E-02 |
| regulation of RNA biosynthetic process (GO:2001141) | 0.55 | 3.46E-02 |
| regulation of DNA-templated transcription (GO:0006355) | 0.55 | 3.43E-02 |

| | | |
|---|---|---|
| organonitrogen compound biosynthetic process (GO:1901566) | 0.55 | 3.79E-03 |
| regulation of biosynthetic process (GO:0009889) | 0.54 | 1.18E-02 |
| regulation of gene expression (GO:0010468) | 0.53 | 1.11E-02 |
| regulation of macromolecule biosynthetic process (GO:0010556) | 0.53 | 9.92E-03 |
| regulation of cellular biosynthetic process (GO:0031326) | 0.53 | 8.12E-03 |
| gene expression (GO:0010467) | 0.5 | 9.54E-03 |
| macromolecule biosynthetic process (GO:0009059) | 0.5 | 2.80E-04 |
| organic cyclic compound biosynthetic process (GO:1901362) | 0.48 | 8.06E-03 |
| nitrogen compound transport (GO:0071705) | 0.48 | 4.10E-02 |
| heterocycle biosynthetic process (GO:0018130) | 0.48 | 9.78E-03 |
| carbohydrate derivative biosynthetic process (GO:1901137) | 0.44 | 2.56E-02 |
| nucleobase-containing compound biosynthetic process (GO:0034654) | 0.39 | 1.06E-02 |
| aromatic compound biosynthetic process (GO:0019438) | 0.39 | 1.47E-03 |
| cellular nitrogen compound biosynthetic process (GO:0044271) | 0.39 | 3.01E-05 |
| RNA metabolic process (GO:0016070) | 0.34 | 1.95E-03 |
| DNA metabolic process (GO:0006259) | 0.33 | 1.52E-03 |
| nucleic acid metabolic process (GO:0090304) | 0.3 | 1.02E-07 |
| lipid biosynthetic process (GO:0008610) | 0.28 | 3.08E-02 |
| macromolecule modification (GO:0043412) | 0.27 | 1.90E-02 |
| translation (GO:0006412) | 0.2 | 2.01E-02 |
| tRNA metabolic process (GO:0006399) | 0.16 | 3.04E-02 |
| RNA processing (GO:0006396) | 0.14 | 8.72E-03 |

| | | |
|---|---|---|
| organelle organization (GO:0006996) | 0.12 | 1.51E-03 |
| ncRNA metabolic process (GO:0034660) | 0.11 | 1.46E-03 |
| cell projection organization (GO:0030030) | 0.1 | 4.05E-02 |
| cell division (GO:0051301) | 0.1 | 3.07E-02 |
| liposaccharide metabolic process (GO:1903509) | 0.1 | 2.29E-02 |
| RNA modification (GO:0009451) | 0.09 | 1.36E-02 |
| DNA recombination (GO:0006310) | 0.07 | 1.45E-03 |
| tRNA processing (GO:0008033) | < 0.01 | 1.56E-02 |
| non-membrane-bounded organelle assembly (GO:0140694) | < 0.01 | 1.54E-02 |
| organelle assembly (GO:0070925) | < 0.01 | 1.52E-02 |
| proton transmembrane transport (GO:1902600) | < 0.01 | 2.60E-02 |
| ribosome biogenesis (GO:0042254) | < 0.01 | 1.47E-03 |
| ribonucleoprotein complex biogenesis (GO:0022613) | < 0.01 | 1.17E-03 |
| ncRNA processing (GO:0034470) | < 0.01 | 9.22E-04 |
| cell cycle process (GO:0022402) | < 0.01 | 4.76E-02 |
| cell cycle (GO:0007049) | < 0.01 | 3.01E-03 |
| transposition (GO:0032196) | < 0.01 | 1.91E-02 |

**Table S5: Comparison of GO terms identified in proteomics (as proteins less abundant in G4 stabilizing conditions) and genes that harbored more insertions in G4 stabilizing conditions**

| **GO terms detected in Tn-seq (more transposon insertions in NMM) and GO terms detected in proteomics (as less abundant in NMM)** |
|---|
| ribosomal small subunit assembly (GO:0000028) |
| ribosomal small subunit biogenesis (GO:0042274) |
| tRNA wobble base modification (GO:0002097) |
| ribosome assembly (GO:0042255) |
| protein-RNA complex organization (GO:0071826) |

| |
|---|
| protein-RNA complex assembly (GO:0022618) |
| negative regulation of translation (GO:0017148) |
| negative regulation of amide metabolic process (GO:0034249) |
| negative regulation of gene expression (GO:0010629) |
| positive regulation of gene expression (GO:0010628) |
| non-membrane-bounded organelle assembly (GO:0140694) |
| organelle assembly (GO:0070925) |
| ribosome biogenesis (GO:0042254) |
| ribonucleoprotein complex biogenesis (GO:0022613) |
| cytoplasmic translation (GO:0002181) |
| post-transcriptional regulation of gene expression (GO:0010608) |
| tRNA processing (GO:0008033) |
| regulation of translation (GO:0006417) |
| tRNA modification (GO:0006400) |
| regulation of amide metabolic process (GO:0034248) |
| translation (GO:0006412) |
| peptide biosynthetic process (GO:0043043) |
| tRNA metabolic process (GO:0006399) |
| amide biosynthetic process (GO:0043604) |
| peptide metabolic process (GO:0006518) |
| RNA processing (GO:0006396) |
| ncRNA processing (GO:0034470) |
| organelle organization (GO:0006996) |
| ncRNA metabolic process (GO:0034660) |
| RNA modification (GO:0009451) |
| amide metabolic process (GO:0043603) |
| protein-containing complex assembly (GO:0065003) |
| gene expression (GO:0010467) |
| protein-containing complex organization (GO:0043933) |
| RNA metabolic process (GO:0016070) |
| macromolecule modification (GO:0043412) |
| macromolecule biosynthetic process (GO:0009059) |
| cellular biosynthetic process (GO:0044249) |
| organic substance biosynthetic process (GO:1901576) |
| protein metabolic process (GO:0019538) |
| biosynthetic process (GO:0009058) |
| cellular nitrogen compound biosynthetic process (GO:0044271) |
| organonitrogen compound biosynthetic process (GO:1901566) |
| cellular component biogenesis (GO:0044085) |

| |
|---|
| cellular nitrogen compound metabolic process (GO:0034641) |
| cellular component organization or biogenesis (GO:0071840) |
| nucleobase-containing compound metabolic process (GO:0006139) |
| organic cyclic compound biosynthetic process (GO:1901362) |
| heterocycle biosynthetic process (GO:0018130) |
| heterocycle metabolic process (GO:0046483) |
| macromolecule metabolic process (GO:0043170) |
| organonitrogen compound metabolic process (GO:1901564) |
| carbohydrate derivative metabolic process (GO:1901135) |
| organic cyclic compound metabolic process (GO:1901360) |
| nitrogen compound metabolic process (GO:0006807) |
| nucleic acid metabolic process (GO:0090304) |
| cellular aromatic compound metabolic process (GO:0006725) |
| cellular metabolic process (GO:0044237) |
| primary metabolic process (GO:0044238) |
| biological regulation (GO:0065007) |
| metabolic process (GO:0008152) |
| organic substance metabolic process (GO:0071704) |
| cellular process (GO:0009987) |
| biological process (GO:0008150) |

**Table S6: Upregulated proteins in ∆*tolC tufA::kan* compared to ∆*tolC***

| Gene ontology biological process | Fold Enrichment | p-value (FDR correction) |
|---|---|---|
| alanine catabolic process (GO:0006524) | 27.97 | 3.79E-02 |
| leucine biosynthetic process (GO:0009098) | 23.31 | 1.50E-03 |
| leucine metabolic process (GO:0006551) | 23.31 | 1.44E-03 |
| valine biosynthetic process (GO:0009099) | 20.34 | 3.85E-04 |
| valine metabolic process (GO:0006573) | 20.34 | 3.68E-04 |
| isoleucine biosynthetic process (GO:0009097) | 19.89 | 1.05E-05 |
| isoleucine metabolic process (GO:0006549) | 19.89 | 9.91E-06 |
| branched-chain amino acid biosynthetic process (GO:0009082) | 17.9 | 1.65E-08 |

| | | |
|---|---|---|
| branched-chain amino acid metabolic process (GO:0009081) | 17.9 | 1.51E-08 |
| indolalkylamine biosynthetic process (GO:0046219) | 14.92 | 2.74E-02 |
| indole-containing compound biosynthetic process (GO:0042435) | 14.92 | 2.67E-02 |
| tryptophan biosynthetic process (GO:0000162) | 14.92 | 2.61E-02 |
| methionine biosynthetic process (GO:0009086) | 11.66 | 1.60E-02 |
| biogenic amine biosynthetic process (GO:0042401) | 11.66 | 1.55E-02 |
| amine biosynthetic process (GO:0009309) | 11.66 | 1.50E-02 |
| methionine metabolic process (GO:0006555) | 11.66 | 1.46E-02 |
| indole-containing compound metabolic process (GO:0042430) | 11.48 | 4.88E-02 |
| indolalkylamine metabolic process (GO:0006586) | 11.48 | 4.78E-02 |
| tryptophan metabolic process (GO:0006568) | 11.48 | 4.69E-02 |
| alpha-amino acid biosynthetic process (GO:1901607) | 9.25 | 7.08E-16 |
| aromatic amino acid family biosynthetic process (GO:0009073) | 8.61 | 1.48E-02 |
| aspartate family amino acid biosynthetic process (GO:0009067) | 8.29 | 1.57E-03 |
| sulfur amino acid biosynthetic process (GO:0000097) | 8.29 | 1.51E-02 |
| amino acid biosynthetic process (GO:0008652) | 8.2 | 1.64E-15 |
| tricarboxylic acid cycle (GO:0006099) | 7.46 | 2.29E-02 |
| non-proteinogenic amino acid metabolic process (GO:0170041) | 6.78 | 3.17E-02 |
| aromatic amino acid metabolic process (GO:0009072) | 6.78 | 3.10E-02 |

| | | |
|---|---|---|
| proteinogenic amino acid biosynthetic process (GO:0170038) | 6.43 | 8.02E-06 |
| L-amino acid biosynthetic process (GO:0170034) | 6.43 | 7.48E-06 |
| alpha-amino acid metabolic process (GO:1901605) | 6.34 | 1.94E-14 |
| dicarboxylic acid biosynthetic process (GO:0043650) | 6.22 | 4.37E-02 |
| sulfur amino acid metabolic process (GO:0000096) | 6.05 | 4.86E-02 |
| organic acid biosynthetic process (GO:0016053) | 5.82 | 1.15E-12 |
| carboxylic acid biosynthetic process (GO:0046394) | 5.82 | 9.56E-13 |
| aspartate family amino acid metabolic process (GO:0009066) | 5.59 | 6.67E-03 |
| amino acid metabolic process (GO:0006520) | 5.48 | 8.86E-14 |
| aerobic respiration (GO:0009060) | 4.97 | 2.65E-02 |
| sulfur compound biosynthetic process (GO:0044272) | 4.8 | 1.56E-02 |
| proteinogenic amino acid metabolic process (GO:0170039) | 4.66 | 1.04E-05 |
| L-amino acid metabolic process (GO:0170033) | 4.63 | 9.68E-06 |
| small molecule biosynthetic process (GO:0044283) | 4.09 | 7.81E-10 |
| carboxylic acid metabolic process (GO:0019752) | 3.43 | 1.12E-11 |
| oxoacid metabolic process (GO:0043436) | 3.38 | 4.87E-12 |
| organic acid metabolic process (GO:0006082) | 3.3 | 1.11E-11 |
| energy derivation by oxidation of organic compounds (GO:0015980) | 3.17 | 3.72E-02 |
| organonitrogen compound biosynthetic process (GO:1901566) | 2.45 | 8.71E-05 |

| | | |
|---|---|---|
| small molecule metabolic process (GO:0044281) | 2.19 | 2.47E-06 |
| organonitrogen compound metabolic process (GO:1901564) | 1.85 | 1.17E-03 |
| biosynthetic process (GO:0009058) | 1.58 | 4.77E-02 |
| primary metabolic process (GO:0044238) | 1.52 | 2.17E-03 |
| nucleobase-containing compound metabolic process (GO:0006139) | 0.3 | 1.85E-02 |

**Table S7: Downregulated proteins in ∆*tolC tufA::kan* compared to ∆*tolC***

| Gene ontology biological process | Fold enrichment | p-value (FDR correction) |
|---|---|---|
| 'de novo' UMP biosynthetic process (GO:0044205) | 60.84 | 2.66E-02 |
| 'de novo' pyrimidine nucleobase biosynthetic process (GO:0006207) | 47.32 | 3.22E-02 |
| pyrimidine ribonucleoside monophosphate biosynthetic process (GO:0009174) | 42.59 | 3.58E-02 |
| UMP biosynthetic process (GO:0006222) | 42.59 | 3.13E-02 |
| pyrimidine nucleobase biosynthetic process (GO:0019856) | 40.56 | 1.66E-02 |
| pyrimidine ribonucleoside monophosphate metabolic process (GO:0009173) | 35.49 | 3.94E-02 |
| UMP metabolic process (GO:0046049) | 35.49 | 3.59E-02 |
| pyrimidine ribonucleotide biosynthetic process (GO:0009220) | 32.76 | 4.03E-02 |
| nucleobase biosynthetic process (GO:0046112) | 28.39 | 2.80E-02 |

| | | |
|---|---|---|
| pyrimidine nucleotide biosynthetic process (GO:0006221) | 24.69 | 3.03E-02 |
| pyrimidine nucleotide metabolic process (GO:0006220) | 21.03 | 3.20E-02 |
| pyrimidine nucleobase metabolic process (GO:0006206) | 18.32 | 2.90E-02 |

**Table S8: Proteins less abundant in ∆*tolC tufA::kan* NMM than ∆*tolC* NMM**

| Gene ontology biological process | Fold enrichment | p-value (FDR correction) |
|---|---|---|
| response to reactive oxygen species (GO:0000302) | 4.47 | 3.11E-02 |
| nicotinamide nucleotide metabolic process (GO:0046496) | 4.36 | 2.58E-02 |
| cellular response to chemical stress (GO:0062197) | 4.19 | 1.06E-02 |
| pyridine nucleotide metabolic process (GO:0019362) | 4.18 | 2.91E-02 |
| response to oxidative stress (GO:0006979) | 3.68 | 2.34E-03 |
| cellular response to stress (GO:0033554) | 1.9 | 2.47E-02 |
| response to stress (GO:0006950) | 1.82 | 3.11E-03 |
| transport (GO:0006810) | 0.38 | 1.32E-03 |
| establishment of localization (GO:0051234) | 0.37 | 1.25E-03 |
| localization (GO:0051179) | 0.37 | 1.33E-03 |
| transmembrane transport (GO:0055085) | 0.35 | 2.59E-03 |

**Table S9: Proteins more abundant in ∆*tolC tufA::kan* NMM than ∆*tolC* NMM**

| Gene ontology biological process | Fold enrichment | p-value (FDR correction) |
|---|---|---|
| leucine biosynthetic process (GO:0009098) | 13.42 | 9.27E-03 |

| | | |
|---|---|---|
| leucine metabolic process (GO:0006551) | 13.42 | 9.10E-03 |
| tRNA aminoacylation for protein translation (GO:0006418) | 7.73 | 8.67E-04 |
| branched-chain amino acid biosynthetic process (GO:0009082) | 7.73 | 8.46E-04 |
| branched-chain amino acid metabolic process (GO:0009081) | 7.73 | 8.25E-04 |
| tRNA aminoacylation (GO:0043039) | 7.73 | 8.06E-04 |
| amino acid activation (GO:0043038) | 7.43 | 9.73E-04 |
| DNA biosynthetic process (GO:0071897) | 6.13 | 4.73E-02 |
| DNA replication (GO:0006260) | 4.73 | 8.57E-04 |
| DNA-templated DNA replication (GO:0006261) | 4.64 | 3.44E-02 |
| alpha-amino acid biosynthetic process (GO:1901607) | 4.61 | 1.95E-07 |
| tRNA metabolic process (GO:0006399) | 4.29 | 2.26E-05 |
| amino acid biosynthetic process (GO:0008652) | 4.26 | 1.80E-07 |
| proteinogenic amino acid biosynthetic process (GO:0170038) | 3.95 | 7.86E-04 |
| L-amino acid biosynthetic process (GO:0170034) | 3.95 | 7.65E-04 |
| amino acid metabolic process (GO:0006520) | 3.86 | 4.08E-11 |
| cytoplasmic translation (GO:0002181) | 3.77 | 3.29E-02 |
| ncRNA metabolic process (GO:0034660) | 3.65 | 1.96E-05 |
| translation (GO:0006412) | 3.58 | 2.44E-04 |
| alpha-amino acid metabolic process (GO:1901605) | 3.54 | 2.84E-07 |
| ribosome biogenesis (GO:0042254) | 3.4 | 3.03E-03 |
| ribonucleoprotein complex biogenesis (GO:0022613) | 3.37 | 3.30E-03 |
| RNA modification (GO:0009451) | 3.27 | 1.16E-02 |

| | | |
|---|---|---|
| peptide biosynthetic process (GO:0043043) | 3.25 | 7.66E-04 |
| organic acid biosynthetic process (GO:0016053) | 3.14 | 1.17E-05 |
| carboxylic acid biosynthetic process (GO:0046394) | 3.14 | 1.13E-05 |
| proteinogenic amino acid metabolic process (GO:0170039) | 3.11 | 5.83E-04 |
| L-amino acid metabolic process (GO:0170033) | 3.09 | 6.22E-04 |
| ncRNA processing (GO:0034470) | 2.95 | 1.78E-02 |
| RNA processing (GO:0006396) | 2.91 | 1.34E-02 |
| peptide metabolic process (GO:0006518) | 2.9 | 2.77E-03 |
| organelle organization (GO:0006996) | 2.78 | 9.79E-03 |
| RNA metabolic process (GO:0016070) | 2.75 | 2.21E-05 |
| organonitrogen compound biosynthetic process (GO:1901566) | 2.74 | 1.14E-11 |
| amide biosynthetic process (GO:0043604) | 2.7 | 4.32E-03 |
| nucleic acid metabolic process (GO:0090304) | 2.37 | 3.83E-07 |
| small molecule biosynthetic process (GO:0044283) | 2.29 | 1.03E-03 |
| gene expression (GO:0010467) | 2.24 | 1.26E-04 |
| macromolecule biosynthetic process (GO:0009059) | 2.21 | 3.63E-07 |
| cellular biosynthetic process (GO:0044249) | 2.2 | 6.47E-14 |
| cellular nitrogen compound biosynthetic process (GO:0044271) | 2.19 | 2.57E-05 |
| organic substance biosynthetic process (GO:1901576) | 2.16 | 4.42E-14 |
| biosynthetic process (GO:0009058) | 2.13 | 5.41E-14 |
| cellular component biogenesis (GO:0044085) | 2 | 9.76E-03 |

| | | |
|---|---|---|
| nucleobase-containing compound metabolic process (GO:0006139) | 1.99 | 2.72E-06 |
| cellular component organization or biogenesis (GO:0071840) | 1.94 | 8.07E-04 |
| macromolecule metabolic process (GO:0043170) | 1.92 | 3.00E-08 |
| cellular nitrogen compound metabolic process (GO:0034641) | 1.9 | 1.67E-07 |
| nitrogen compound metabolic process (GO:0006807) | 1.88 | 5.76E-14 |
| heterocycle metabolic process (GO:0046483) | 1.84 | 1.05E-05 |
| organic cyclic compound metabolic process (GO:1901360) | 1.82 | 7.84E-06 |
| cellular aromatic compound metabolic process (GO:0006725) | 1.82 | 1.57E-05 |
| cellular component organization (GO:0016043) | 1.78 | 2.79E-02 |
| organonitrogen compound metabolic process (GO:1901564) | 1.77 | 1.15E-05 |
| primary metabolic process (GO:0044238) | 1.72 | 1.12E-11 |
| cellular metabolic process (GO:0044237) | 1.52 | 4.80E-08 |
| organic substance metabolic process (GO:0071704) | 1.51 | 4.40E-08 |
| metabolic process (GO:0008152) | 1.43 | 3.43E-07 |
| cellular process (GO:0009987) | 1.24 | 5.26E-05 |
| biological_process (GO:0008150) | 1.19 | 1.01E-05 |
| Unclassified (UNCLASSIFIED) | 0.36 | 9.62E-06 |

**Table S10 Transcripts less abundant in ∆*tolC* NMM compared to ∆*tolC***

| Gene ontology biological process | Fold Enrichment | p-value (FDR correction) |
|---|---|---|

| | | |
|---|---|---|
| GMP biosynthetic process (GO:0006177) | 3.87 | 1.66E-02 |
| tripeptide import across plasma membrane (GO:0140207) | 3.87 | 1.65E-02 |
| 'de novo' UMP biosynthetic process (GO:0044205) | 3.87 | 1.64E-03 |
| DNA-templated transcription termination (GO:0006353) | 3.87 | 5.31E-03 |
| CTP metabolic process (GO:0046036) | 3.87 | 4.98E-02 |
| dTDP-rhamnose biosynthetic process (GO:0019305) | 3.87 | 4.96E-02 |
| RNA 5'-end processing (GO:0000966) | 3.87 | 4.94E-02 |
| CTP biosynthetic process (GO:0006241) | 3.87 | 4.92E-02 |
| proton motive force-driven plasma membrane ATP synthesis (GO:0042777) | 3.87 | 4.73E-04 |
| protein insertion into membrane from inner side (GO:0032978) | 3.87 | 4.69E-04 |
| UDP-glucose metabolic process (GO:0006011) | 3.87 | 4.90E-02 |
| ncRNA 5'-end processing (GO:0034471) | 3.87 | 4.88E-02 |
| IMP salvage (GO:0032264) | 3.87 | 4.86E-02 |
| sister chromatid cohesion (GO:0007062) | 3.87 | 4.85E-02 |
| pyrimidine ribonucleoside triphosphate biosynthetic process (GO:0009209) | 3.87 | 4.83E-02 |
| dTDP-rhamnose metabolic process (GO:0046383) | 3.87 | 4.81E-02 |
| malonyl-CoA biosynthetic process (GO:2001295) | 3.87 | 4.79E-02 |
| malonyl-CoA metabolic process (GO:2001293) | 3.87 | 4.78E-02 |

| | | |
|---|---|---|
| CDP-diacylglycerol metabolic process (GO:0046341) | 3.87 | 4.76E-02 |
| cytoplasmic translation (GO:0002181) | 3.8 | 5.04E-30 |
| ribosomal small subunit assembly (GO:0000028) | 3.68 | 3.89E-09 |
| protein-RNA complex organization (GO:0071826) | 3.62 | 2.82E-20 |
| protein-RNA complex assembly (GO:0022618) | 3.62 | 2.68E-20 |
| ribonucleoside triphosphate biosynthetic process (GO:0009201) | 3.59 | 7.29E-06 |
| pyrimidine ribonucleotide biosynthetic process (GO:0009220) | 3.57 | 2.54E-05 |
| ribosomal large subunit assembly (GO:0000027) | 3.56 | 2.20E-10 |
| ribosome assembly (GO:0042255) | 3.55 | 2.08E-20 |
| regulation of DNA-templated transcription elongation (GO:0032784) | 3.52 | 2.82E-04 |
| purine nucleoside triphosphate biosynthetic process (GO:0009145) | 3.52 | 2.79E-04 |
| purine ribonucleoside triphosphate biosynthetic process (GO:0009206) | 3.52 | 2.77E-04 |
| pyrimidine ribonucleoside monophosphate biosynthetic process (GO:0009174) | 3.48 | 8.83E-04 |
| UMP biosynthetic process (GO:0006222) | 3.48 | 8.77E-04 |
| 'de novo' pyrimidine nucleobase biosynthetic process (GO:0006207) | 3.44 | 2.82E-03 |
| protein transport by the Sec complex (GO:0043952) | 3.44 | 2.80E-03 |
| ATP biosynthetic process (GO:0006754) | 3.44 | 2.78E-03 |
| electron transport coupled proton transport (GO:0015990) | 3.44 | 2.76E-03 |

| | | |
|---|---|---|
| energy coupled proton transmembrane transport, against electrochemical gradient (GO:0015988) | 3.44 | 2.74E-03 |
| proton motive force-driven ATP synthesis (GO:0015986) | 3.44 | 2.72E-03 |
| ribosomal large subunit biogenesis (GO:0042273) | 3.42 | 1.27E-09 |
| thioester biosynthetic process (GO:0035384) | 3.39 | 8.08E-03 |
| enzyme-directed rRNA pseudouridine synthesis (GO:0000455) | 3.39 | 8.04E-03 |
| IMP biosynthetic process (GO:0006188) | 3.39 | 1.20E-05 |
| acyl-CoA biosynthetic process (GO:0071616) | 3.39 | 7.99E-03 |
| rRNA pseudouridine synthesis (GO:0031118) | 3.39 | 7.95E-03 |
| pyrimidine nucleoside monophosphate biosynthetic process (GO:0009130) | 3.35 | 3.96E-05 |
| DNA topological change (GO:0006265) | 3.32 | 2.15E-02 |
| purine ribonucleotide salvage (GO:0106380) | 3.32 | 2.14E-02 |
| maintenance of translational fidelity (GO:1990145) | 3.32 | 2.13E-02 |
| ribonucleoside monophosphate biosynthetic process (GO:0009156) | 3.23 | 3.51E-11 |
| 'de novo' IMP biosynthetic process (GO:0006189) | 3.23 | 1.12E-03 |
| nucleoside monophosphate biosynthetic process (GO:0009124) | 3.21 | 1.51E-12 |
| purine ribonucleoside monophosphate biosynthetic process (GO:0009168) | 3.2 | 6.32E-07 |

| | | |
|---|---|---|
| purine nucleoside monophosphate biosynthetic process (GO:0009127) | 3.2 | 6.25E-07 |
| IMP metabolic process (GO:0046040) | 3.19 | 4.79E-05 |
| purine nucleotide salvage (GO:0032261) | 3.17 | 3.15E-03 |
| translation (GO:0006412) | 3.16 | 9.03E-36 |
| ribosomal small subunit biogenesis (GO:0042274) | 3.13 | 2.45E-07 |
| ribonucleoprotein complex biogenesis (GO:0022613) | 3.11 | 5.49E-29 |
| ribosome biogenesis (GO:0042254) | 3.1 | 1.66E-28 |
| nucleobase biosynthetic process (GO:0046112) | 3.1 | 1.81E-05 |
| purine ribonucleoside monophosphate metabolic process (GO:0009167) | 3.06 | 2.17E-06 |
| purine nucleoside monophosphate metabolic process (GO:0009126) | 3.06 | 2.15E-06 |
| ribonucleotide biosynthetic process (GO:0009260) | 3.06 | 1.77E-16 |
| nucleoside triphosphate biosynthetic process (GO:0009142) | 3.06 | 5.20E-05 |
| pyrimidine nucleobase biosynthetic process (GO:0019856) | 3.04 | 1.19E-03 |
| intracellular transport (GO:0046907) | 3.01 | 2.15E-02 |
| intracellular protein transport (GO:0006886) | 3.01 | 2.14E-02 |
| intracellular protein transmembrane transport (GO:0065002) | 3.01 | 2.13E-02 |
| menaquinone biosynthetic process (GO:0009234) | 3.01 | 2.12E-02 |
| menaquinone metabolic process (GO:0009233) | 3.01 | 2.11E-02 |
| purine ribonucleotide biosynthetic process (GO:0009152) | 2.98 | 7.23E-12 |

| | | |
|---|---|---|
| xenobiotic detoxification by transmembrane export across the cell outer membrane (GO:0140330) | 2.98 | 3.19E-03 |
| export across cell outer membrane (GO:0140317) | 2.98 | 3.17E-03 |
| ribose phosphate biosynthetic process (GO:0046390) | 2.98 | 6.05E-16 |
| ribonucleoside monophosphate metabolic process (GO:0009161) | 2.98 | 1.28E-09 |
| pyrimidine nucleoside monophosphate metabolic process (GO:0009129) | 2.96 | 4.31E-04 |
| nucleotide salvage (GO:0043173) | 2.96 | 4.27E-04 |
| peptide biosynthetic process (GO:0043043) | 2.9 | 1.17E-30 |
| pyrimidine ribonucleoside monophosphate metabolic process (GO:0009173) | 2.9 | 8.44E-03 |
| pseudouridine synthesis (GO:0001522) | 2.9 | 8.40E-03 |
| UMP metabolic process (GO:0046049) | 2.9 | 8.35E-03 |
| pyrimidine ribonucleotide metabolic process (GO:0009218) | 2.9 | 1.17E-03 |
| pyrimidine nucleotide biosynthetic process (GO:0006221) | 2.86 | 5.12E-05 |
| DNA replication initiation (GO:0006270) | 2.82 | 2.01E-02 |
| nucleoside monophosphate metabolic process (GO:0009123) | 2.79 | 3.96E-10 |
| tRNA aminoacylation for protein translation (GO:0006418) | 2.79 | 4.84E-05 |
| tRNA aminoacylation (GO:0043039) | 2.79 | 4.80E-05 |
| protein insertion into membrane (GO:0051205) | 2.76 | 7.73E-03 |
| purine nucleotide biosynthetic process (GO:0006164) | 2.73 | 1.43E-11 |

| | | |
|---|---|---|
| DNA unwinding involved in DNA replication (GO:0006268) | 2.71 | 4.80E-02 |
| amide biosynthetic process (GO:0043604) | 2.69 | 6.46E-31 |
| amino acid activation (GO:0043038) | 2.68 | 1.11E-04 |
| peptide metabolic process (GO:0006518) | 2.67 | 1.09E-26 |
| non-membrane-bounded organelle assembly (GO:0140694) | 2.65 | 1.15E-12 |
| organelle assembly (GO:0070925) | 2.65 | 1.12E-12 |
| nucleoside bisphosphate biosynthetic process (GO:0033866) | 2.58 | 1.54E-02 |
| purine nucleoside bisphosphate biosynthetic process (GO:0034033) | 2.58 | 1.53E-02 |
| ribonucleoside bisphosphate biosynthetic process (GO:0034030) | 2.58 | 1.53E-02 |
| nucleotide biosynthetic process (GO:0009165) | 2.58 | 7.85E-16 |
| rRNA processing (GO:0006364) | 2.58 | 3.30E-07 |
| pyrimidine nucleotide metabolic process (GO:0006220) | 2.58 | 2.46E-04 |
| fatty acid biosynthetic process (GO:0006633) | 2.58 | 7.02E-04 |
| nucleoside phosphate biosynthetic process (GO:1901293) | 2.58 | 7.58E-16 |
| purine-containing compound biosynthetic process (GO:0072522) | 2.56 | 3.51E-10 |
| rRNA modification (GO:0000154) | 2.54 | 6.69E-05 |
| DNA-templated DNA replication (GO:0006261) | 2.51 | 1.88E-05 |
| thioester metabolic process (GO:0035383) | 2.5 | 1.31E-02 |
| establishment of protein localization to membrane (GO:0090150) | 2.5 | 1.31E-02 |
| acyl-CoA metabolic process (GO:0006637) | 2.5 | 1.30E-02 |

| | | |
|---|---|---|
| translational elongation (GO:0006414) | 2.49 | 3.23E-02 |
| amino acid import across plasma membrane (GO:0089718) | 2.49 | 4.72E-04 |
| cellular component disassembly (GO:0022411) | 2.49 | 3.22E-02 |
| enterobacterial common antigen biosynthetic process (GO:0009246) | 2.49 | 3.20E-02 |
| enterobacterial common antigen metabolic process (GO:0046378) | 2.49 | 3.19E-02 |
| rRNA metabolic process (GO:0016072) | 2.47 | 1.39E-06 |
| rRNA base methylation (GO:0070475) | 2.44 | 1.05E-02 |
| tRNA modification (GO:0006400) | 2.36 | 3.67E-07 |
| ribonucleotide metabolic process (GO:0009259) | 2.32 | 8.18E-11 |
| negative regulation of termination of DNA-templated transcription (GO:0060567) | 2.32 | 1.75E-02 |
| transcription antitermination (GO:0031564) | 2.32 | 1.74E-02 |
| tRNA metabolic process (GO:0006399) | 2.32 | 2.18E-11 |
| negative regulation of protein-containing complex disassembly (GO:0043242) | 2.32 | 1.74E-02 |
| lipopolysaccharide core region metabolic process (GO:0046401) | 2.32 | 5.47E-03 |
| lipopolysaccharide core region biosynthetic process (GO:0009244) | 2.32 | 5.43E-03 |
| DNA replication (GO:0006260) | 2.3 | 2.13E-06 |
| oxidative phosphorylation (GO:0006119) | 2.29 | 1.46E-02 |
| RNA modification (GO:0009451) | 2.27 | 7.92E-10 |
| pyrimidine-containing compound biosynthetic process (GO:0072528) | 2.27 | 3.31E-04 |

| | | |
|---|---|---|
| ncRNA metabolic process (GO:0034660) | 2.25 | 1.48E-14 |
| DNA conformation change (GO:0071103) | 2.25 | 4.19E-03 |
| ncRNA processing (GO:0034470) | 2.24 | 5.59E-11 |
| amide metabolic process (GO:0043603) | 2.24 | 1.35E-21 |
| negative regulation of translation (GO:0017148) | 2.23 | 1.47E-02 |
| nucleotide-sugar metabolic process (GO:0009225) | 2.23 | 1.46E-02 |
| purine ribonucleotide metabolic process (GO:0009150) | 2.23 | 7.62E-08 |
| RNA processing (GO:0006396) | 2.23 | 8.11E-12 |
| tRNA processing (GO:0008033) | 2.23 | 2.83E-07 |
| macromolecule methylation (GO:0043414) | 2.21 | 9.19E-05 |
| organelle organization (GO:0006996) | 2.2 | 3.12E-13 |
| rRNA methylation (GO:0031167) | 2.19 | 3.21E-02 |
| ribose phosphate metabolic process (GO:0019693) | 2.19 | 5.80E-10 |
| negative regulation of amide metabolic process (GO:0034249) | 2.15 | 1.76E-02 |
| RNA methylation (GO:0001510) | 2.13 | 1.78E-03 |
| positive regulation of gene expression (GO:0010628) | 2.12 | 1.02E-02 |
| regulation of protein-containing complex disassembly (GO:0043244) | 2.12 | 1.01E-02 |
| ribonucleoside bisphosphate metabolic process (GO:0033875) | 2.08 | 3.07E-02 |
| nucleoside bisphosphate metabolic process (GO:0033865) | 2.08 | 3.05E-02 |
| localization within membrane (GO:0051668) | 2.08 | 3.04E-02 |
| purine nucleoside bisphosphate metabolic process (GO:0034032) | 2.08 | 3.03E-02 |

| | | |
|---|---|---|
| protein localization to membrane (GO:0072657) | 2.08 | 3.02E-02 |
| nucleoside triphosphate metabolic process (GO:0009141) | 2.07 | 2.81E-03 |
| regulation of termination of DNA-templated transcription (GO:0031554) | 2.06 | 1.77E-02 |
| ribonucleoside triphosphate metabolic process (GO:0009199) | 2.05 | 1.64E-02 |
| cellular nitrogen compound biosynthetic process (GO:0044271) | 2.02 | 2.80E-36 |
| negative regulation of cellular component organization (GO:0051129) | 2.01 | 4.15E-02 |
| organophosphate biosynthetic process (GO:0090407) | 2 | 2.81E-13 |
| oligosaccharide biosynthetic process (GO:0009312) | 2 | 4.87E-02 |
| negative regulation of protein metabolic process (GO:0051248) | 2 | 3.39E-02 |
| carbohydrate derivative biosynthetic process (GO:1901137) | 2 | 1.78E-17 |
| gene expression (GO:0010467) | 1.98 | 3.81E-28 |
| purine nucleotide metabolic process (GO:0006163) | 1.97 | 3.53E-07 |
| regulation of translation (GO:0006417) | 1.97 | 8.09E-04 |
| chromosome organization (GO:0051276) | 1.94 | 4.52E-03 |
| purine nucleoside triphosphate metabolic process (GO:0009144) | 1.94 | 3.07E-02 |
| regulation of amide metabolic process (GO:0034248) | 1.94 | 1.56E-03 |
| L-alpha-amino acid transmembrane transport (GO:1902475) | 1.94 | 1.53E-02 |
| methylation (GO:0032259) | 1.94 | 8.10E-04 |

| | | |
|---|---|---|
| nucleotide metabolic process (GO:0009117) | 1.92 | 9.65E-10 |
| organonitrogen compound biosynthetic process (GO:1901566) | 1.9 | 1.01E-32 |
| nucleoside phosphate metabolic process (GO:0006753) | 1.9 | 1.59E-09 |
| aerobic respiration (GO:0009060) | 1.87 | 3.10E-03 |
| macromolecule modification (GO:0043412) | 1.87 | 1.02E-07 |
| negative regulation of gene expression (GO:0010629) | 1.86 | 1.01E-02 |
| glycosaminoglycan biosynthetic process (GO:0006024) | 1.86 | 1.37E-02 |
| aminoglycan biosynthetic process (GO:0006023) | 1.86 | 1.37E-02 |
| peptidoglycan biosynthetic process (GO:0009252) | 1.86 | 1.36E-02 |
| macromolecule biosynthetic process (GO:0009059) | 1.85 | 3.34E-34 |
| post-transcriptional regulation of gene expression (GO:0010608) | 1.85 | 2.94E-03 |
| xenobiotic transport (GO:0042908) | 1.84 | 3.65E-02 |
| nucleobase-containing compound biosynthetic process (GO:0034654) | 1.83 | 2.96E-12 |
| response to antibiotic (GO:0046677) | 1.83 | 4.80E-05 |
| peptidoglycan-based cell wall biogenesis (GO:0009273) | 1.83 | 1.22E-02 |
| protein-containing complex organization (GO:0043933) | 1.82 | 4.70E-08 |
| cell wall macromolecule biosynthetic process (GO:0044038) | 1.82 | 1.55E-02 |
| proton transmembrane transport (GO:1902600) | 1.81 | 4.69E-03 |
| lipid biosynthetic process (GO:0008610) | 1.8 | 2.57E-06 |
| response to radiation (GO:0009314) | 1.8 | 2.67E-04 |
| cell wall biogenesis (GO:0042546) | 1.79 | 1.93E-02 |

| | | |
|---|---|---|
| L-amino acid transport (GO:0015807) | 1.78 | 2.34E-02 |
| protein-containing complex assembly (GO:0065003) | 1.74 | 3.14E-06 |
| RNA metabolic process (GO:0016070) | 1.74 | 2.20E-10 |
| phospholipid biosynthetic process (GO:0008654) | 1.74 | 2.64E-02 |
| amino acid transmembrane transport (GO:0003333) | 1.73 | 7.37E-03 |
| cellular biosynthetic process (GO:0044249) | 1.71 | 1.26E-47 |
| phospholipid metabolic process (GO:0006644) | 1.71 | 2.22E-02 |
| pyrimidine-containing compound metabolic process (GO:0072527) | 1.7 | 2.09E-02 |
| protein metabolic process (GO:0019538) | 1.7 | 1.09E-12 |
| organic substance biosynthetic process (GO:1901576) | 1.7 | 1.90E-47 |
| liposaccharide metabolic process (GO:1903509) | 1.68 | 6.70E-03 |
| biosynthetic process (GO:0009058) | 1.68 | 4.08E-46 |
| nucleobase-containing small molecule metabolic process (GO:0055086) | 1.68 | 3.35E-08 |
| inorganic cation transmembrane transport (GO:0098662) | 1.67 | 3.78E-04 |
| purine-containing compound metabolic process (GO:0072521) | 1.65 | 3.78E-04 |
| aromatic compound biosynthetic process (GO:0019438) | 1.65 | 2.16E-10 |
| monoatomic cation transmembrane transport (GO:0098655) | 1.65 | 4.85E-04 |
| amino acid transport (GO:0006865) | 1.64 | 6.65E-03 |
| cellular component biogenesis (GO:0044085) | 1.63 | 7.51E-11 |

| | | |
|---|---|---|
| peptidoglycan metabolic process (GO:0000270) | 1.62 | 2.34E-02 |
| glycosaminoglycan metabolic process (GO:0030203) | 1.62 | 2.33E-02 |
| regulation of protein metabolic process (GO:0051246) | 1.62 | 3.34E-02 |
| heterocycle biosynthetic process (GO:0018130) | 1.62 | 4.01E-10 |
| lipopolysaccharide biosynthetic process (GO:0009103) | 1.61 | 4.48E-02 |
| organic cyclic compound biosynthetic process (GO:1901362) | 1.61 | 1.85E-10 |
| carbohydrate derivative metabolic process (GO:1901135) | 1.61 | 2.53E-11 |
| cellular component organization or biogenesis (GO:0071840) | 1.6 | 4.57E-15 |
| cellular nitrogen compound metabolic process (GO:0034641) | 1.6 | 5.63E-31 |
| organophosphate metabolic process (GO:0019637) | 1.6 | 9.03E-08 |
| inorganic ion transmembrane transport (GO:0098660) | 1.59 | 8.82E-04 |
| cell wall organization or biogenesis (GO:0071554) | 1.59 | 1.75E-02 |
| aminoglycan metabolic process (GO:0006022) | 1.59 | 2.80E-02 |
| polysaccharide biosynthetic process (GO:0000271) | 1.54 | 1.51E-02 |
| monoatomic ion transmembrane transport (GO:0034220) | 1.51 | 2.95E-03 |
| nucleobase-containing compound metabolic process (GO:0006139) | 1.5 | 1.64E-15 |
| cellular component organization (GO:0016043) | 1.48 | 1.33E-08 |
| carbohydrate biosynthetic process (GO:0016051) | 1.47 | 2.23E-02 |

| | | |
|---|---|---|
| macromolecule metabolic process (GO:0043170) | 1.47 | 2.78E-20 |
| lipid metabolic process (GO:0006629) | 1.47 | 2.71E-03 |
| monoatomic cation transport (GO:0006812) | 1.47 | 5.65E-03 |
| polysaccharide metabolic process (GO:0005976) | 1.46 | 4.04E-02 |
| cellular lipid metabolic process (GO:0044255) | 1.44 | 4.62E-02 |
| heterocycle metabolic process (GO:0046483) | 1.43 | 1.98E-14 |
| nucleic acid metabolic process (GO:0090304) | 1.4 | 1.12E-06 |
| nitrogen compound transport (GO:0071705) | 1.4 | 2.20E-03 |
| cellular component assembly (GO:0022607) | 1.4 | 2.94E-03 |
| cellular aromatic compound metabolic process (GO:0006725) | 1.4 | 2.03E-12 |
| organic cyclic compound metabolic process (GO:1901360) | 1.39 | 1.35E-12 |
| organonitrogen compound metabolic process (GO:1901564) | 1.38 | 4.48E-13 |
| phosphate-containing compound metabolic process (GO:0006796) | 1.37 | 1.10E-04 |
| phosphorus metabolic process (GO:0006793) | 1.36 | 8.57E-05 |
| small molecule biosynthetic process (GO:0044283) | 1.36 | 3.08E-03 |
| monoatomic ion transport (GO:0006811) | 1.36 | 2.16E-02 |
| nitrogen compound metabolic process (GO:0006807) | 1.35 | 3.40E-19 |
| response to abiotic stimulus (GO:0009628) | 1.33 | 1.89E-02 |

| | | |
|---|---|---|
| primary metabolic process (GO:0044238) | 1.29 | 8.57E-17 |
| cellular metabolic process (GO:0044237) | 1.26 | 9.98E-18 |
| organic substance metabolic process (GO:0071704) | 1.22 | 4.24E-13 |
| metabolic process (GO:0008152) | 1.2 | 1.56E-12 |
| cellular process (GO:0009987) | 1.17 | 1.83E-20 |
| biological_process (GO:0008150) | 1.11 | 3.57E-14 |
| regulation of RNA metabolic process (GO:0051252) | 0.74 | 2.23E-02 |
| regulation of RNA biosynthetic process (GO:2001141) | 0.73 | 1.76E-02 |
| regulation of DNA-templated transcription (GO:0006355) | 0.73 | 1.75E-02 |
| organonitrogen compound catabolic process (GO:1901565) | 0.67 | 2.32E-02 |
| Unclassified (UNCLASSIFIED) | 0.63 | 3.68E-14 |
| organic substance catabolic process (GO:1901575) | 0.63 | 1.44E-06 |
| cellular catabolic process (GO:0044248) | 0.63 | 1.23E-04 |
| response to external stimulus (GO:0009605) | 0.62 | 1.68E-02 |
| catabolic process (GO:0009056) | 0.61 | 1.70E-07 |
| carboxylic acid catabolic process (GO:0046395) | 0.58 | 3.59E-03 |
| organic acid catabolic process (GO:0016054) | 0.57 | 1.60E-03 |
| cell communication (GO:0007154) | 0.55 | 3.76E-03 |
| small molecule catabolic process (GO:0044282) | 0.55 | 5.17E-06 |
| signaling (GO:0023052) | 0.5 | 3.02E-02 |
| carbohydrate transmembrane transport (GO:0034219) | 0.49 | 1.36E-02 |
| carbohydrate transport (GO:0008643) | 0.48 | 3.91E-03 |

| | | |
|---|---|---|
| intracellular signal transduction (GO:0035556) | 0.46 | 4.01E-02 |
| phosphorelay signal transduction system (GO:0000160) | 0.42 | 3.22E-02 |
| DNA transposition (GO:0006313) | 0.37 | 4.71E-02 |
| transposition (GO:0032196) | 0.35 | 1.08E-02 |
| cellular response to xenobiotic stimulus (GO:0071466) | 0.12 | 2.14E-02 |
| xenobiotic metabolic process (GO:0006805) | 0.12 | 2.13E-02 |
| benzene-containing compound metabolic process (GO:0042537) | 0.12 | 1.54E-02 |
| primary alcohol catabolic process (GO:0034310) | < 0.01 | 1.93E-02 |
| cellular biogenic amine catabolic process (GO:0042402) | < 0.01 | 3.75E-03 |
| xenobiotic catabolic process (GO:0042178) | < 0.01 | 8.90E-03 |
| amine catabolic process (GO:0009310) | < 0.01 | 3.72E-03 |

**Table S11 Transcripts more abundant in Δ*tolC* NMM compared to Δ*tolC***

| Gene ontology biological process | Fold Enrichment | p-value (FDR correction) |
|---|---|---|
| xanthine metabolic process (GO:0046110) | > 100 | 2.03E-02 |
| glutaminyl-tRNAGln biosynthesis via transamidation (GO:0070681) | > 100 | 4.77E-04 |
| deoxyadenosine metabolic process (GO:0046090) | > 100 | 4.11E-02 |
| deoxyadenosine catabolic process (GO:0006157) | > 100 | 3.97E-02 |
| deoxyribonucleoside catabolic process (GO:0046121) | 99 | 3.43E-03 |
| deoxyribonucleoside metabolic process (GO:0009120) | 54 | 1.70E-02 |

| | | |
|---|---|---|
| serine family amino acid biosynthetic process (GO:0009070) | 33 | 4.75E-02 |
| nucleoside catabolic process (GO:0009164) | 31.68 | 7.79E-03 |
| glycosyl compound catabolic process (GO:1901658) | 29.12 | 1.61E-03 |
| nucleobase-containing small molecule catabolic process (GO:0034656) | 28.29 | 1.16E-02 |
| serine family amino acid metabolic process (GO:0009069) | 23.29 | 1.93E-02 |
| carbohydrate catabolic process (GO:0016052) | 14.4 | 4.38E-04 |
| glycosyl compound metabolic process (GO:1901657) | 11.65 | 4.16E-02 |
| purine ribonucleotide catabolic process (GO:0009154) | 10.76 | 4.93E-02 |
| nucleotide catabolic process (GO:0009166) | 9.5 | 2.74E-02 |
| nucleoside phosphate catabolic process (GO:1901292) | 8.87 | 3.72E-02 |
| amino acid metabolic process (GO:0006520) | 7.48 | 6.57E-04 |
| small molecule catabolic process (GO:0044282) | 7.27 | 2.10E-04 |
| carbohydrate derivative catabolic process (GO:1901136) | 7.07 | 1.73E-02 |
| nucleobase-containing compound catabolic process (GO:0034655) | 5.3 | 3.30E-02 |
| carbohydrate metabolic process (GO:0005975) | 4.44 | 4.44E-02 |
| cellular catabolic process (GO:0044248) | 3.5 | 1.58E-03 |
| amide metabolic process (GO:0043603) | 3.35 | 4.04E-02 |
| organonitrogen compound catabolic process (GO:1901565) | 3.34 | 3.76E-03 |

| | | |
|---|---|---|
| organic substance catabolic process (GO:1901575) | 3.27 | 3.91E-04 |
| catabolic process (GO:0009056) | 2.86 | 6.80E-04 |
| small molecule metabolic process (GO:0044281) | 2.66 | 1.73E-02 |
| organonitrogen compound metabolic process (GO:1901564) | 1.82 | 1.94E-02 |
| cellular metabolic process (GO:0044237) | 1.73 | 2.88E-03 |
| organic substance metabolic process (GO:0071704) | 1.62 | 3.82E-03 |
| metabolic process (GO:0008152) | 1.61 | 2.26E-03 |
| primary metabolic process (GO:0044238) | 1.6 | 1.86E-02 |

**Table S12 Transcripts less abundant in ∆*tolC tufA::kan* NMM compared to ∆*tolC* NMM**

| Gene ontology biological process | Fold Enrichment | p-value (FDR correction) |
|---|---|---|
| galactarate metabolic process (GO:0019580) | 4.11 | 6.80E-03 |
| aldaric acid catabolic process (GO:0019579) | 4.11 | 2.19E-02 |
| aldaric acid metabolic process (GO:0019577) | 4.11 | 2.39E-03 |
| hydrogen sulfide biosynthetic process (GO:0070814) | 4.11 | 6.72E-03 |
| hydrogen sulfide metabolic process (GO:0070813) | 4.11 | 6.65E-03 |
| glucarate metabolic process (GO:0019392) | 4.11 | 6.58E-03 |
| D-glucarate metabolic process (GO:0042836) | 4.11 | 6.51E-03 |
| sulfate assimilation (GO:0000103) | 3.65 | 3.58E-03 |
| lysine catabolic process (GO:0006554) | 3.52 | 2.69E-02 |

| | | |
|---|---|---|
| uracil catabolic process (GO:0006212) | 3.28 | 9.50E-03 |
| macrolide metabolic process (GO:0033067) | 3.19 | 2.70E-02 |
| enterobactin metabolic process (GO:0009238) | 3.19 | 2.68E-02 |
| polyketide metabolic process (GO:0030638) | 3.19 | 2.66E-02 |
| phenol-containing compound metabolic process (GO:0018958) | 3.08 | 9.01E-03 |
| pyrimidine nucleobase catabolic process (GO:0006208) | 3.08 | 8.92E-03 |
| catechol-containing compound metabolic process (GO:0009712) | 2.99 | 2.42E-02 |
| uracil metabolic process (GO:0019860) | 2.99 | 2.40E-02 |
| antibiotic metabolic process (GO:0016999) | 2.74 | 4.95E-02 |
| pyrimidine-containing compound catabolic process (GO:0072529) | 2.35 | 2.95E-02 |
| cellular response to starvation (GO:0009267) | 2.15 | 3.53E-03 |
| cellular response to nutrient levels (GO:0031669) | 2.1 | 5.24E-03 |
| response to starvation (GO:0042594) | 2.05 | 2.68E-03 |
| organic hydroxy compound catabolic process (GO:1901616) | 1.97 | 7.04E-03 |
| alcohol catabolic process (GO:0046164) | 1.96 | 1.92E-02 |
| response to nutrient levels (GO:0031667) | 1.82 | 2.89E-03 |
| cellular response to external stimulus (GO:0071496) | 1.79 | 5.85E-03 |
| cellular response to extracellular stimulus (GO:0031668) | 1.79 | 5.78E-03 |
| response to extracellular stimulus (GO:0009991) | 1.69 | 2.62E-03 |
| response to oxidative stress (GO:0006979) | 1.65 | 1.09E-02 |

| | | |
|---|---|---|
| cellular response to stress (GO:0033554) | 1.48 | 8.34E-06 |
| DNA damage response (GO:0006974) | 1.42 | 2.91E-03 |
| response to stress (GO:0006950) | 1.42 | 8.11E-07 |
| cellular response to stimulus (GO:0051716) | 1.36 | 7.08E-05 |
| small molecule catabolic process (GO:0044282) | 1.35 | 8.25E-03 |
| catabolic process (GO:0009056) | 1.3 | 2.25E-03 |
| organic substance catabolic process (GO:1901575) | 1.29 | 3.51E-03 |
| cellular catabolic process (GO:0044248) | 1.28 | 4.49E-02 |
| Unclassified (UNCLASSIFIED) | 1.24 | 1.53E-04 |
| response to stimulus (GO:0050896) | 1.23 | 2.27E-04 |
| biological_process (GO:0008150) | 0.93 | 1.48E-04 |
| cellular process (GO:0009987) | 0.88 | 2.39E-07 |
| metabolic process (GO:0008152) | 0.88 | 6.02E-04 |
| organic substance metabolic process (GO:0071704) | 0.86 | 3.84E-04 |
| cellular metabolic process (GO:0044237) | 0.85 | 8.75E-05 |
| primary metabolic process (GO:0044238) | 0.85 | 3.97E-04 |
| organonitrogen compound metabolic process (GO:1901564) | 0.82 | 6.94E-03 |
| localization (GO:0051179) | 0.78 | 6.76E-03 |
| transport (GO:0006810) | 0.78 | 6.48E-03 |
| nitrogen compound metabolic process (GO:0006807) | 0.78 | 6.74E-07 |
| establishment of localization (GO:0051234) | 0.77 | 4.33E-03 |
| macromolecule metabolic process (GO:0043170) | 0.75 | 3.39E-05 |
| transmembrane transport (GO:0055085) | 0.75 | 3.33E-03 |

| | | |
|---|---|---|
| cellular aromatic compound metabolic process (GO:0006725) | 0.73 | 2.29E-05 |
| organic cyclic compound metabolic process (GO:1901360) | 0.72 | 7.70E-06 |
| carbohydrate derivative metabolic process (GO:1901135) | 0.71 | 1.14E-02 |
| heterocycle metabolic process (GO:0046483) | 0.7 | 1.18E-06 |
| organic substance transport (GO:0071702) | 0.69 | 1.32E-03 |
| nucleobase-containing compound metabolic process (GO:0006139) | 0.68 | 8.03E-06 |
| biosynthetic process (GO:0009058) | 0.68 | 1.29E-09 |
| organic substance biosynthetic process (GO:1901576) | 0.67 | 2.11E-10 |
| macromolecule biosynthetic process (GO:0009059) | 0.66 | 2.32E-05 |
| nucleic acid metabolic process (GO:0090304) | 0.66 | 1.53E-04 |
| gene expression (GO:0010467) | 0.65 | 1.37E-03 |
| heterocycle biosynthetic process (GO:0018130) | 0.64 | 2.32E-03 |
| cellular biosynthetic process (GO:0044249) | 0.64 | 9.62E-12 |
| cellular nitrogen compound metabolic process (GO:0034641) | 0.64 | 9.29E-11 |
| RNA metabolic process (GO:0016070) | 0.62 | 6.48E-03 |
| organic cyclic compound biosynthetic process (GO:1901362) | 0.62 | 4.37E-04 |
| cellular component biogenesis (GO:0044085) | 0.61 | 3.18E-04 |
| aromatic compound biosynthetic process (GO:0019438) | 0.6 | 6.50E-04 |
| cellular component organization (GO:0016043) | 0.6 | 1.35E-05 |
| amide metabolic process (GO:0043603) | 0.6 | 1.13E-02 |

| | | |
|---|---|---|
| nucleobase-containing compound biosynthetic process (GO:0034654) | 0.59 | 3.93E-03 |
| cellular component organization or biogenesis (GO:0071840) | 0.58 | 3.32E-07 |
| carbohydrate derivative biosynthetic process (GO:1901137) | 0.53 | 3.20E-04 |
| import into cell (GO:0098657) | 0.52 | 1.32E-02 |
| small molecule biosynthetic process (GO:0044283) | 0.52 | 2.24E-05 |
| cellular nitrogen compound biosynthetic process (GO:0044271) | 0.51 | 6.66E-09 |
| alpha-amino acid biosynthetic process (GO:1901607) | 0.51 | 2.78E-02 |
| cellular localization (GO:0051641) | 0.5 | 1.59E-02 |
| monoatomic cation transmembrane transport (GO:0098655) | 0.49 | 1.23E-02 |
| organic acid biosynthetic process (GO:0016053) | 0.48 | 3.89E-04 |
| carboxylic acid biosynthetic process (GO:0046394) | 0.48 | 3.80E-04 |
| inorganic cation transmembrane transport (GO:0098662) | 0.47 | 7.84E-03 |
| peptide biosynthetic process (GO:0043043) | 0.47 | 6.57E-03 |
| DNA recombination (GO:0006310) | 0.46 | 8.27E-03 |
| organonitrogen compound biosynthetic process (GO:1901566) | 0.45 | 9.92E-13 |
| carbohydrate transport (GO:0008643) | 0.45 | 5.57E-03 |
| import across plasma membrane (GO:0098739) | 0.44 | 5.76E-03 |
| cell wall organization or biogenesis (GO:0071554) | 0.43 | 2.38E-02 |
| amide biosynthetic process (GO:0043604) | 0.42 | 2.74E-04 |
| cell wall organization (GO:0071555) | 0.42 | 4.59E-02 |
| amino acid biosynthetic process (GO:0008652) | 0.41 | 8.74E-04 |

| | | |
|---|---|---|
| peptidoglycan metabolic process (GO:0000270) | 0.41 | 3.42E-02 |
| glycosaminoglycan metabolic process (GO:0030203) | 0.41 | 3.39E-02 |
| tRNA processing (GO:0008033) | 0.39 | 4.76E-02 |
| RNA processing (GO:0006396) | 0.38 | 2.28E-03 |
| ncRNA processing (GO:0034470) | 0.38 | 2.80E-03 |
| organelle organization (GO:0006996) | 0.35 | 2.24E-04 |
| macromolecule localization (GO:0033036) | 0.34 | 9.00E-05 |
| protein transport (GO:0015031) | 0.34 | 7.84E-03 |
| cellular macromolecule localization (GO:0070727) | 0.34 | 3.70E-03 |
| protein localization (GO:0008104) | 0.34 | 3.65E-03 |
| nucleotide biosynthetic process (GO:0009165) | 0.33 | 2.24E-03 |
| nucleoside phosphate biosynthetic process (GO:1901293) | 0.33 | 2.20E-03 |
| tRNA metabolic process (GO:0006399) | 0.33 | 1.62E-03 |
| proton transmembrane transport (GO:1902600) | 0.32 | 2.54E-02 |
| ncRNA metabolic process (GO:0034660) | 0.32 | 4.97E-05 |
| RNA modification (GO:0009451) | 0.31 | 2.17E-03 |
| establishment of protein localization (GO:0045184) | 0.31 | 2.17E-03 |
| translation (GO:0006412) | 0.27 | 5.22E-05 |
| cell projection organization (GO:0030030) | 0.26 | 2.29E-03 |
| rRNA metabolic process (GO:0016072) | 0.26 | 4.25E-02 |
| purine-containing compound biosynthetic process (GO:0072522) | 0.25 | 6.86E-03 |
| ribose phosphate biosynthetic process (GO:0046390) | 0.25 | 6.78E-03 |
| phosphoenolpyruvate-dependent sugar phosphotransferase system (GO:0009401) | 0.21 | 4.22E-03 |

| | | |
|---|---|---|
| purine nucleotide biosynthetic process (GO:0006164) | 0.2 | 4.34E-03 |
| nucleoside monophosphate metabolic process (GO:0009123) | 0.16 | 6.54E-03 |
| ribosome biogenesis (GO:0042254) | 0.16 | 9.13E-06 |
| ribonucleoprotein complex biogenesis (GO:0022613) | 0.16 | 7.40E-06 |
| ribonucleotide biosynthetic process (GO:0009260) | 0.13 | 5.68E-04 |
| membrane organization (GO:0061024) | 0.12 | 2.68E-02 |
| ribonucleoside monophosphate biosynthetic process (GO:0009156) | 0.11 | 1.93E-02 |
| non-membrane-bounded organelle assembly (GO:0140694) | 0.11 | 7.24E-05 |
| organelle assembly (GO:0070925) | 0.11 | 6.96E-05 |
| ribonucleoside monophosphate metabolic process (GO:0009161) | 0.11 | 1.30E-02 |
| nucleoside monophosphate biosynthetic process (GO:0009124) | 0.1 | 9.03E-03 |
| purine ribonucleotide biosynthetic process (GO:0009152) | 0.09 | 2.46E-03 |
| cytoplasmic translation (GO:0002181) | < 0.01 | 2.31E-05 |
| defense response to other organism (GO:0098542) | < 0.01 | 2.46E-02 |
| defense response (GO:0006952) | < 0.01 | 2.44E-02 |
| protein-RNA complex organization (GO:0071826) | < 0.01 | 3.83E-04 |
| protein transmembrane transport (GO:0071806) | < 0.01 | 1.30E-03 |
| ribosomal large subunit biogenesis (GO:0042273) | < 0.01 | 2.41E-02 |
| ribosome assembly (GO:0042255) | < 0.01 | 1.68E-04 |
| protein-RNA complex assembly (GO:0022618) | < 0.01 | 3.73E-04 |
| ribosomal large subunit assembly (GO:0000027) | < 0.01 | 3.54E-02 |

**Table S13 Transcripts more abundant in Δ*tolC tufA::kan* NMM compared to Δ*tolC* NMM**

| Gene ontology biological process | Fold Enrichment | p-value (FDR correction) |
|---|---|---|
| purine nucleoside triphosphate biosynthetic process (GO:0009145) | 4.06 | 1.03E-05 |
| thioester biosynthetic process (GO:0035384) | 4.06 | 4.27E-04 |
| maltodextrin transmembrane transport (GO:0042956) | 4.06 | 1.55E-02 |
| dextrin transport (GO:0042955) | 4.06 | 1.54E-02 |
| dTDP-rhamnose biosynthetic process (GO:0019305) | 4.06 | 4.91E-02 |
| GMP biosynthetic process (GO:0006177) | 4.06 | 1.53E-02 |
| regulation of membrane invagination (GO:1905153) | 4.06 | 1.52E-02 |
| proton motive force-driven plasma membrane ATP synthesis (GO:0042777) | 4.06 | 4.23E-04 |
| acyl-CoA biosynthetic process (GO:0071616) | 4.06 | 4.18E-04 |
| acetyl-CoA biosynthetic process (GO:0006085) | 4.06 | 4.89E-02 |
| tripeptide import across plasma membrane (GO:0140207) | 4.06 | 1.51E-02 |
| IMP salvage (GO:0032264) | 4.06 | 4.87E-02 |
| purine ribonucleoside triphosphate biosynthetic process (GO:0009206) | 4.06 | 1.01E-05 |
| dTDP-rhamnose metabolic process (GO:0046383) | 4.06 | 4.84E-02 |
| malonyl-CoA biosynthetic process (GO:2001295) | 4.06 | 4.82E-02 |
| malonyl-CoA metabolic process (GO:2001293) | 4.06 | 4.80E-02 |
| maltose transport (GO:0015768) | 4.06 | 4.84E-03 |
| ATP biosynthetic process (GO:0006754) | 4.06 | 1.25E-04 |

| | | |
|---|---|---|
| proton motive force-driven ATP synthesis (GO:0015986) | 4.06 | 1.23E-04 |
| cytoplasmic translation (GO:0002181) | 3.99 | 1.81E-30 |
| ribonucleoside triphosphate biosynthetic process (GO:0009201) | 3.77 | 7.17E-06 |
| electron transport coupled proton transport (GO:0015990) | 3.61 | 2.45E-03 |
| energy coupled proton transmembrane transport, against electrochemical gradient (GO:0015988) | 3.61 | 2.43E-03 |
| leucine biosynthetic process (GO:0009098) | 3.55 | 7.20E-03 |
| protein insertion into membrane from inner side (GO:0032978) | 3.55 | 7.15E-03 |
| leucine metabolic process (GO:0006551) | 3.55 | 7.10E-03 |
| galactose transmembrane transport (GO:0015757) | 3.48 | 1.99E-02 |
| purine ribonucleotide salvage (GO:0106380) | 3.48 | 1.98E-02 |
| maintenance of translational fidelity (GO:1990145) | 3.48 | 1.97E-02 |
| ribosomal small subunit assembly (GO:0000028) | 3.45 | 1.41E-06 |
| protein-RNA complex organization (GO:0071826) | 3.44 | 1.35E-15 |
| protein-RNA complex assembly (GO:0022618) | 3.44 | 1.29E-15 |
| ribosomal large subunit assembly (GO:0000027) | 3.41 | 4.93E-08 |
| ribosome assembly (GO:0042255) | 3.4 | 5.89E-16 |
| regulation of DNA-templated transcription elongation (GO:0032784) | 3.32 | 2.64E-03 |
| purine nucleotide salvage (GO:0032261) | 3.32 | 2.62E-03 |
| ribosomal large subunit biogenesis (GO:0042273) | 3.28 | 1.87E-07 |
| protein transport by the Sec complex (GO:0043952) | 3.16 | 1.97E-02 |

| | | |
|---|---|---|
| xenobiotic detoxification by transmembrane export across the cell outer membrane (GO:0140330) | 3.12 | 2.65E-03 |
| export across cell outer membrane (GO:0140317) | 3.12 | 2.63E-03 |
| translation (GO:0006412) | 3.11 | 1.87E-30 |
| nucleotide salvage (GO:0043173) | 3.1 | 3.38E-04 |
| translational termination (GO:0006415) | 3.04 | 4.91E-02 |
| plasmid maintenance (GO:0006276) | 3.04 | 4.89E-02 |
| protein-containing complex disassembly (GO:0032984) | 3.04 | 4.87E-02 |
| nucleoside triphosphate biosynthetic process (GO:0009142) | 2.99 | 3.13E-04 |
| ribosomal small subunit biogenesis (GO:0042274) | 2.97 | 1.28E-05 |
| non-membrane-bounded organelle assembly (GO:0140694) | 2.89 | 4.93E-15 |
| organelle assembly (GO:0070925) | 2.89 | 4.71E-15 |
| thioester metabolic process (GO:0035383) | 2.87 | 2.22E-03 |
| acyl-CoA metabolic process (GO:0006637) | 2.87 | 2.20E-03 |
| peptide biosynthetic process (GO:0043043) | 2.83 | 8.55E-26 |
| ribonucleoprotein complex biogenesis (GO:0022613) | 2.83 | 1.03E-19 |
| ribosome biogenesis (GO:0042254) | 2.81 | 2.81E-19 |
| tRNA aminoacylation for protein translation (GO:0006418) | 2.76 | 2.00E-04 |
| branched-chain amino acid biosynthetic process (GO:0009082) | 2.76 | 1.97E-04 |
| branched-chain amino acid metabolic process (GO:0009081) | 2.76 | 1.95E-04 |
| tRNA aminoacylation (GO:0043039) | 2.76 | 1.93E-04 |
| nucleoside bisphosphate biosynthetic process (GO:0033866) | 2.71 | 1.27E-02 |

| | | |
|---|---|---|
| purine nucleoside bisphosphate biosynthetic process (GO:0034033) | 2.71 | 1.26E-02 |
| ribonucleoside bisphosphate biosynthetic process (GO:0034030) | 2.71 | 1.25E-02 |
| chromosome segregation (GO:0007059) | 2.71 | 1.24E-02 |
| amino acid activation (GO:0043038) | 2.65 | 4.01E-04 |
| purine ribonucleotide biosynthetic process (GO:0009152) | 2.62 | 3.15E-07 |
| protein insertion into membrane (GO:0051205) | 2.61 | 2.79E-02 |
| enterobacterial common antigen biosynthetic process (GO:0009246) | 2.61 | 2.77E-02 |
| enterobacterial common antigen metabolic process (GO:0046378) | 2.61 | 2.76E-02 |
| amide biosynthetic process (GO:0043604) | 2.6 | 1.37E-25 |
| ribonucleotide biosynthetic process (GO:0009260) | 2.55 | 1.25E-08 |
| peptide metabolic process (GO:0006518) | 2.55 | 3.76E-21 |
| ribose phosphate biosynthetic process (GO:0046390) | 2.5 | 1.82E-08 |
| oligosaccharide transport (GO:0015772) | 2.48 | 1.76E-02 |
| amino acid import across plasma membrane (GO:0089718) | 2.46 | 1.41E-03 |
| purine nucleotide biosynthetic process (GO:0006164) | 2.46 | 1.45E-07 |
| negative regulation of termination of DNA-templated transcription (GO:0060567) | 2.44 | 1.40E-02 |
| pyrimidine nucleoside monophosphate biosynthetic process (GO:0009130) | 2.44 | 4.81E-02 |
| disaccharide transport (GO:0015766) | 2.44 | 4.79E-02 |
| isoleucine biosynthetic process (GO:0009097) | 2.44 | 4.77E-02 |
| transcription antitermination (GO:0031564) | 2.44 | 1.39E-02 |
| negative regulation of protein-containing complex disassembly (GO:0043242) | 2.44 | 1.38E-02 |

| | | |
|---|---|---|
| purine-containing compound salvage (GO:0043101) | 2.44 | 4.75E-02 |
| isoleucine metabolic process (GO:0006549) | 2.44 | 4.72E-02 |
| oxidative phosphorylation (GO:0006119) | 2.4 | 1.09E-02 |
| organelle organization (GO:0006996) | 2.34 | 5.43E-15 |
| purine-containing compound biosynthetic process (GO:0072522) | 2.31 | 1.98E-06 |
| nucleoside monophosphate biosynthetic process (GO:0009124) | 2.28 | 7.13E-04 |
| ribonucleoside monophosphate biosynthetic process (GO:0009156) | 2.26 | 2.57E-03 |
| positive regulation of gene expression (GO:0010628) | 2.23 | 9.01E-03 |
| regulation of protein-containing complex disassembly (GO:0043244) | 2.23 | 8.95E-03 |
| nucleotide biosynthetic process (GO:0009165) | 2.21 | 1.16E-08 |
| nucleoside phosphate biosynthetic process (GO:1901293) | 2.21 | 1.13E-08 |
| fatty acid biosynthetic process (GO:0006633) | 2.2 | 3.94E-02 |
| taxis (GO:0042330) | 2.19 | 3.22E-03 |
| chemotaxis (GO:0006935) | 2.19 | 3.20E-03 |
| DNA-templated DNA replication (GO:0006261) | 2.19 | 3.17E-03 |
| locomotion (GO:0040011) | 2.19 | 3.15E-03 |
| ribonucleoside bisphosphate metabolic process (GO:0033875) | 2.19 | 2.84E-02 |
| nucleoside bisphosphate metabolic process (GO:0033865) | 2.19 | 2.83E-02 |
| negative regulation of translation (GO:0017148) | 2.19 | 2.81E-02 |
| purine nucleoside bisphosphate metabolic process (GO:0034032) | 2.19 | 2.80E-02 |
| regulation of termination of DNA-templated transcription (GO:0031554) | 2.17 | 1.49E-02 |

| | | |
|---|---|---|
| amide metabolic process (GO:0043603) | 2.13 | 4.37E-17 |
| tRNA metabolic process (GO:0006399) | 2.11 | 1.88E-07 |
| negative regulation of amide metabolic process (GO:0034249) | 2.11 | 3.51E-02 |
| ribonucleoside monophosphate metabolic process (GO:0009161) | 2.08 | 9.94E-03 |
| rRNA processing (GO:0006364) | 2.08 | 3.43E-03 |
| tRNA modification (GO:0006400) | 2.06 | 4.03E-04 |
| DNA replication (GO:0006260) | 2.06 | 3.99E-04 |
| bacterial-type flagellum organization (GO:0044781) | 2.03 | 2.70E-02 |
| ribonucleoside triphosphate metabolic process (GO:0009199) | 2.03 | 2.82E-02 |
| purine nucleoside triphosphate metabolic process (GO:0009144) | 2.03 | 2.81E-02 |
| nucleoside monophosphate metabolic process (GO:0009123) | 2.03 | 2.51E-03 |
| rRNA modification (GO:0000154) | 2.03 | 2.69E-02 |
| tricarboxylic acid cycle (GO:0006099) | 2.03 | 3.48E-02 |
| purine ribonucleoside triphosphate metabolic process (GO:0009205) | 2.03 | 3.47E-02 |
| dicarboxylic acid biosynthetic process (GO:0043650) | 2.03 | 2.09E-02 |
| rRNA metabolic process (GO:0016072) | 1.99 | 5.88E-03 |
| ncRNA metabolic process (GO:0034660) | 1.99 | 1.84E-08 |
| nucleoside triphosphate metabolic process (GO:0009141) | 1.98 | 1.05E-02 |
| purine ribonucleotide metabolic process (GO:0009150) | 1.98 | 1.33E-04 |
| neutral amino acid transport (GO:0015804) | 1.98 | 2.55E-02 |
| ribonucleotide metabolic process (GO:0009259) | 1.97 | 1.89E-05 |
| aerobic respiration (GO:0009060) | 1.96 | 1.69E-03 |
| tRNA processing (GO:0008033) | 1.95 | 4.21E-04 |
| RNA modification (GO:0009451) | 1.94 | 4.53E-05 |

| | | |
|---|---|---|
| L-alpha-amino acid transmembrane transport (GO:1902475) | 1.93 | 2.62E-02 |
| RNA processing (GO:0006396) | 1.93 | 3.25E-06 |
| ribose phosphate metabolic process (GO:0019693) | 1.92 | 1.60E-05 |
| proton transmembrane transport (GO:1902600) | 1.9 | 2.55E-03 |
| ncRNA processing (GO:0034470) | 1.9 | 1.90E-05 |
| purine nucleotide metabolic process (GO:0006163) | 1.88 | 2.40E-05 |
| protein transport (GO:0015031) | 1.88 | 4.68E-04 |
| protein transmembrane transport (GO:0071806) | 1.88 | 3.93E-02 |
| archaeal or bacterial-type flagellum-dependent cell motility (GO:0097588) | 1.85 | 1.03E-02 |
| bacterial-type flagellum-dependent cell motility (GO:0071973) | 1.85 | 1.02E-02 |
| cilium or flagellum-dependent cell motility (GO:0001539) | 1.85 | 1.02E-02 |
| organonitrogen compound biosynthetic process (GO:1901566) | 1.85 | 7.26E-27 |
| cell motility (GO:0048870) | 1.84 | 7.19E-03 |
| gene expression (GO:0010467) | 1.83 | 1.43E-18 |
| macromolecule methylation (GO:0043414) | 1.82 | 3.09E-02 |
| cellular nitrogen compound biosynthetic process (GO:0044271) | 1.81 | 1.16E-21 |
| negative regulation of gene expression (GO:0010629) | 1.8 | 2.78E-02 |
| establishment of protein localization (GO:0045184) | 1.79 | 1.20E-03 |
| L-amino acid transport (GO:0015807) | 1.79 | 3.60E-02 |
| cellular macromolecule localization (GO:0070727) | 1.78 | 1.03E-03 |
| protein localization (GO:0008104) | 1.78 | 1.02E-03 |
| amino acid transmembrane transport (GO:0003333) | 1.76 | 1.03E-02 |

| | | |
|---|---|---|
| macromolecule localization (GO:0033036) | 1.73 | 6.29E-05 |
| nucleotide metabolic process (GO:0009117) | 1.72 | 1.94E-05 |
| regulation of translation (GO:0006417) | 1.72 | 4.46E-02 |
| nucleoside phosphate metabolic process (GO:0006753) | 1.71 | 1.91E-05 |
| regulation of amide metabolic process (GO:0034248) | 1.69 | 4.82E-02 |
| response to antibiotic (GO:0046677) | 1.69 | 3.15E-03 |
| protein-containing complex organization (GO:0043933) | 1.68 | 5.01E-05 |
| macromolecule biosynthetic process (GO:0009059) | 1.65 | 6.16E-19 |
| regulation of cellular component organization (GO:0051128) | 1.65 | 4.09E-02 |
| amino acid transport (GO:0006865) | 1.63 | 1.32E-02 |
| response to radiation (GO:0009314) | 1.63 | 1.31E-02 |
| inorganic cation transmembrane transport (GO:0098662) | 1.62 | 2.61E-03 |
| protein-containing complex assembly (GO:0065003) | 1.62 | 4.25E-04 |
| respiratory electron transport chain (GO:0022904) | 1.61 | 4.32E-02 |
| purine-containing compound metabolic process (GO:0072521) | 1.61 | 2.56E-03 |
| protein metabolic process (GO:0019538) | 1.61 | 8.17E-09 |
| macromolecule modification (GO:0043412) | 1.61 | 1.16E-03 |
| monoatomic cation transmembrane transport (GO:0098655) | 1.61 | 2.51E-03 |
| organophosphate biosynthetic process (GO:0090407) | 1.6 | 1.29E-04 |
| carbohydrate derivative biosynthetic process (GO:1901137) | 1.6 | 4.75E-06 |
| nucleobase-containing compound biosynthetic process (GO:0034654) | 1.6 | 7.21E-06 |

| | | |
|---|---|---|
| cellular localization (GO:0051641) | 1.58 | 5.88E-03 |
| cellular biosynthetic process (GO:0044249) | 1.56 | 7.12E-28 |
| import across plasma membrane (GO:0098739) | 1.56 | 1.09E-02 |
| inorganic ion transmembrane transport (GO:0098660) | 1.56 | 3.48E-03 |
| RNA metabolic process (GO:0016070) | 1.55 | 2.10E-05 |
| nitrogen compound transport (GO:0071705) | 1.54 | 1.94E-05 |
| organic substance biosynthetic process (GO:1901576) | 1.54 | 8.90E-27 |
| amino acid biosynthetic process (GO:0008652) | 1.53 | 9.70E-03 |
| organic acid biosynthetic process (GO:0016053) | 1.52 | 8.50E-04 |
| carboxylic acid biosynthetic process (GO:0046394) | 1.52 | 8.42E-04 |
| biosynthetic process (GO:0009058) | 1.52 | 1.41E-25 |
| cellular component organization or biogenesis (GO:0071840) | 1.51 | 4.28E-10 |
| monoatomic ion transmembrane transport (GO:0034220) | 1.51 | 5.26E-03 |
| cellular component biogenesis (GO:0044085) | 1.48 | 8.73E-06 |
| import into cell (GO:0098657) | 1.48 | 2.36E-02 |
| aromatic compound biosynthetic process (GO:0019438) | 1.48 | 3.54E-05 |
| carboxylic acid transport (GO:0046942) | 1.46 | 2.75E-02 |
| organic acid transport (GO:0015849) | 1.45 | 2.84E-02 |
| nucleobase-containing small molecule metabolic process (GO:0055086) | 1.45 | 1.95E-03 |
| cellular component organization (GO:0016043) | 1.43 | 2.84E-06 |
| monoatomic cation transport (GO:0006812) | 1.43 | 2.13E-02 |

| | | |
|---|---|---|
| cellular nitrogen compound metabolic process (GO:0034641) | 1.42 | 1.47E-14 |
| organic cyclic compound biosynthetic process (GO:1901362) | 1.42 | 1.16E-04 |
| monoatomic ion transport (GO:0006811) | 1.41 | 1.06E-02 |
| transmembrane transport (GO:0055085) | 1.39 | 2.57E-07 |
| small molecule biosynthetic process (GO:0044283) | 1.39 | 2.41E-03 |
| carbohydrate derivative metabolic process (GO:1901135) | 1.38 | 3.97E-04 |
| heterocycle biosynthetic process (GO:0018130) | 1.38 | 1.12E-03 |
| cellular component assembly (GO:0022607) | 1.37 | 1.00E-02 |
| organic substance transport (GO:0071702) | 1.37 | 6.58E-05 |
| amino acid metabolic process (GO:0006520) | 1.36 | 2.10E-02 |
| organonitrogen compound metabolic process (GO:1901564) | 1.34 | 2.21E-09 |
| organophosphate metabolic process (GO:0019637) | 1.33 | 1.69E-02 |
| nucleobase-containing compound metabolic process (GO:0006139) | 1.32 | 5.46E-06 |
| transport (GO:0006810) | 1.32 | 8.75E-06 |
| macromolecule metabolic process (GO:0043170) | 1.32 | 1.35E-08 |
| establishment of localization (GO:0051234) | 1.31 | 1.22E-05 |
| localization (GO:0051179) | 1.31 | 1.87E-05 |
| phosphate-containing compound metabolic process (GO:0006796) | 1.28 | 1.35E-02 |
| heterocycle metabolic process (GO:0046483) | 1.26 | 4.51E-05 |
| nitrogen compound metabolic process (GO:0006807) | 1.26 | 4.18E-10 |

| | | |
|---|---|---|
| nucleic acid metabolic process (GO:0090304) | 1.26 | 1.04E-02 |
| cellular aromatic compound metabolic process (GO:0006725) | 1.25 | 7.66E-05 |
| phosphorus metabolic process (GO:0006793) | 1.25 | 2.78E-02 |
| primary metabolic process (GO:0044238) | 1.25 | 5.25E-11 |
| organic cyclic compound metabolic process (GO:1901360) | 1.24 | 1.48E-04 |
| cellular metabolic process (GO:0044237) | 1.22 | 5.13E-11 |
| cellular process (GO:0009987) | 1.18 | 3.81E-21 |
| organic substance metabolic process (GO:0071704) | 1.16 | 4.21E-06 |
| metabolic process (GO:0008152) | 1.16 | 3.27E-07 |
| biological_process (GO:0008150) | 1.12 | 8.60E-16 |
| regulation of RNA biosynthetic process (GO:2001141) | 0.75 | 4.13E-02 |
| regulation of DNA-templated transcription (GO:0006355) | 0.75 | 4.12E-02 |
| organic substance catabolic process (GO:1901575) | 0.75 | 8.31E-03 |
| catabolic process (GO:0009056) | 0.74 | 4.04E-03 |
| Unclassified (UNCLASSIFIED) | 0.6 | 9.07E-16 |
| cell communication (GO:0007154) | 0.51 | 1.99E-03 |
| signal transduction (GO:0007165) | 0.45 | 2.78E-02 |
| signaling (GO:0023052) | 0.45 | 1.55E-02 |
| intracellular signal transduction (GO:0035556) | 0.37 | 1.77E-02 |
| phosphorelay signal transduction system (GO:0000160) | 0.33 | 1.01E-02 |
| DNA transposition (GO:0006313) | 0.31 | 2.92E-02 |
| transposition (GO:0032196) | 0.3 | 9.71E-03 |
| benzene-containing compound metabolic process (GO:0042537) | 0.12 | 2.71E-02 |
| cellular response to xenobiotic stimulus (GO:0071466) | < 0.01 | 4.90E-03 |

| | | |
|---|---|---|
| xenobiotic metabolic process (GO:0006805) | < 0.01 | 4.87E-03 |
| xenobiotic catabolic process (GO:0042178) | < 0.01 | 1.69E-02 |

**Table S14 Transcripts less abundant in Δ*tolC tufA::kan* compared to Δ*tolC***

| Gene ontology biological process | Fold Enrichment | p-value (FDR correction) |
|---|---|---|
| cytosine metabolic process (GO:0019858) | 60.29 | 5.74E-02 |
| NADH oxidation (GO:0006116) | 60.29 | 5.36E-02 |
| 'de novo' UMP biosynthetic process (GO:0044205) | 25.84 | 3.93E-02 |
| pyrimidine ribonucleotide biosynthetic process (GO:0009220) | 18.55 | 4.39E-02 |
| pyrimidine nucleobase biosynthetic process (GO:0019856) | 17.23 | 4.55E-02 |
| aerobic electron transport chain (GO:0019646) | 17.23 | 3.64E-02 |
| pyrimidine ribonucleotide metabolic process (GO:0009218) | 15.07 | 4.04E-02 |
| pyrimidine nucleotide biosynthetic process (GO:0006221) | 13.11 | 4.32E-02 |
| siderophore-dependent iron import into cell (GO:0033214) | 12.06 | 5.11E-02 |
| nucleobase biosynthetic process (GO:0046112) | 12.06 | 4.81E-02 |
| pyrimidine nucleobase metabolic process (GO:0006206) | 11.67 | 2.66E-02 |
| pyrimidine nucleotide metabolic process (GO:0006220) | 11.16 | 3.28E-02 |
| iron import into cell (GO:0033212) | 9.72 | 4.37E-02 |
| nucleobase metabolic process (GO:0009112) | 7.09 | 4.17E-02 |
| iron ion transport (GO:0006826) | 6.58 | 5.87E-02 |

| | | |
|---|---|---|
| pyrimidine-containing compound metabolic process (GO:0072527) | 6.39 | 3.89E-02 |
| monoatomic cation transport (GO:0006812) | 3.7 | 4.31E-02 |

**Table S15 Transcripts more abundant in ∆*tolC tufA::kan* compared to ∆*tolC***

| Gene ontology biological process | Fold enrichment | p-value (FDR correction) |
|---|---|---|
| galactitol metabolic process (GO:0019402) | 17.46 | 5.66E-05 |
| behavior (GO:0007610) | 17.46 | 9.16E-03 |
| galactitol catabolic process (GO:0019404) | 17.46 | 9.01E-03 |
| response to mechanical stimulus (GO:0009612) | 17.46 | 8.86E-03 |
| multicellular organismal process (GO:0032501) | 17.46 | 8.72E-03 |
| mechanosensory behavior (GO:0007638) | 17.46 | 8.59E-03 |
| valine biosynthetic process (GO:0009099) | 14.29 | 6.06E-08 |
| valine metabolic process (GO:0006573) | 14.29 | 5.66E-08 |
| arginine catabolic process to succinate (GO:0019545) | 13.97 | 2.97E-03 |
| arginine catabolic process to glutamate (GO:0019544) | 13.97 | 2.91E-03 |
| alanine catabolic process (GO:0006524) | 13.1 | 2.81E-02 |
| methylgalactoside transport (GO:0015765) | 13.1 | 2.77E-02 |
| leucine biosynthetic process (GO:0009098) | 13.1 | 7.82E-05 |
| pyruvate family amino acid metabolic process (GO:0009078) | 13.1 | 2.73E-02 |
| glycine decarboxylation via glycine cleavage system (GO:0019464) | 13.1 | 2.70E-02 |
| L-alanine metabolic process (GO:0042851) | 13.1 | 2.66E-02 |
| leucine metabolic process (GO:0006551) | 13.1 | 7.58E-05 |
| branched-chain amino acid biosynthetic process (GO:0009082) | 11.88 | 1.20E-13 |
| branched-chain amino acid metabolic process (GO:0009081) | 11.88 | 1.07E-13 |

| | | |
|---|---|---|
| isoleucine biosynthetic process (GO:0009097) | 11.64 | 1.37E-07 |
| isoleucine metabolic process (GO:0006549) | 11.64 | 1.29E-07 |
| galactose transmembrane transport (GO:0015757) | 9.98 | 1.37E-02 |
| glutamate metabolic process (GO:0006536) | 8.22 | 1.36E-04 |
| tricarboxylic acid cycle (GO:0006099) | 8.15 | 4.15E-08 |
| hexitol metabolic process (GO:0006059) | 7.94 | 9.36E-03 |
| alanine metabolic process (GO:0006522) | 7.76 | 3.80E-02 |
| non-proteinogenic amino acid catabolic process (GO:0170044) | 7.28 | 1.40E-02 |
| alditol catabolic process (GO:0019405) | 6.79 | 2.16E-03 |
| polyol catabolic process (GO:0046174) | 6.11 | 4.15E-03 |
| phenylacetate catabolic process (GO:0010124) | 5.82 | 3.94E-02 |
| dipeptide transmembrane transport (GO:0035442) | 5.24 | 2.61E-02 |
| alpha-amino acid biosynthetic process (GO:1901607) | 5.2 | 8.90E-15 |
| amino acid biosynthetic process (GO:0008652) | 5.08 | 1.54E-16 |
| non-proteinogenic amino acid metabolic process (GO:0170041) | 4.76 | 3.74E-03 |
| alditol metabolic process (GO:0019400) | 4.76 | 3.67E-03 |
| proteinogenic amino acid catabolic process (GO:0170040) | 4.68 | 3.08E-05 |
| L-amino acid catabolic process (GO:0170035) | 4.68 | 2.97E-05 |
| aerobic respiration (GO:0009060) | 4.66 | 1.46E-05 |
| alpha-amino acid catabolic process (GO:1901606) | 4.37 | 1.58E-05 |
| pyruvate metabolic process (GO:0006090) | 4.37 | 1.38E-02 |
| polyol metabolic process (GO:0019751) | 4.25 | 8.83E-03 |
| amino acid catabolic process (GO:0009063) | 4.15 | 7.50E-06 |
| alpha-amino acid metabolic process (GO:1901605) | 4.1 | 3.19E-15 |

| | | |
|---|---|---|
| organic hydroxy compound catabolic process (GO:1901616) | 4.03 | 1.78E-03 |
| alcohol catabolic process (GO:0046164) | 3.8 | 9.26E-03 |
| amino acid metabolic process (GO:0006520) | 3.64 | 1.89E-14 |
| organic acid biosynthetic process (GO:0016053) | 3.58 | 2.14E-11 |
| carboxylic acid biosynthetic process (GO:0046394) | 3.58 | 1.95E-11 |
| proteinogenic amino acid biosynthetic process (GO:0170038) | 3.41 | 4.48E-04 |
| L-amino acid biosynthetic process (GO:0170034) | 3.41 | 4.36E-04 |
| proteinogenic amino acid metabolic process (GO:0170039) | 3.33 | 6.00E-07 |
| L-amino acid metabolic process (GO:0170033) | 3.31 | 6.73E-07 |
| aspartate family amino acid metabolic process (GO:0009066) | 3.2 | 1.97E-02 |
| carboxylic acid catabolic process (GO:0046395) | 3.04 | 2.03E-07 |
| dicarboxylic acid metabolic process (GO:0043648) | 2.91 | 4.62E-03 |
| carboxylic acid metabolic process (GO:0019752) | 2.9 | 7.22E-18 |
| organic acid catabolic process (GO:0016054) | 2.88 | 7.03E-07 |
| monocarboxylic acid catabolic process (GO:0072329) | 2.79 | 7.50E-03 |
| oxoacid metabolic process (GO:0043436) | 2.77 | 4.01E-17 |
| organic acid metabolic process (GO:0006082) | 2.7 | 1.57E-16 |
| cellular respiration (GO:0045333) | 2.56 | 9.12E-03 |
| small molecule biosynthetic process (GO:0044283) | 2.54 | 3.18E-07 |
| small molecule catabolic process (GO:0044282) | 2.47 | 4.51E-07 |
| energy derivation by oxidation of organic compounds (GO:0015980) | 2.23 | 4.20E-02 |

| | | |
|---|---|---|
| monocarboxylic acid metabolic process (GO:0032787) | 2.2 | 2.15E-03 |
| organonitrogen compound catabolic process (GO:1901565) | 2.17 | 2.46E-03 |
| cellular catabolic process (GO:0044248) | 2.09 | 9.43E-05 |
| small molecule metabolic process (GO:0044281) | 1.97 | 4.25E-11 |
| organic substance catabolic process (GO:1901575) | 1.86 | 2.05E-04 |
| catabolic process (GO:0009056) | 1.85 | 1.83E-04 |
| organonitrogen compound metabolic process (GO:1901564) | 1.44 | 9.85E-03 |
| biological_process (GO:0008150) | 1.14 | 1.95E-03 |
| Unclassified (UNCLASSIFIED) | 0.55 | 1.91E-03 |
| heterocycle metabolic process (GO:0046483) | 0.54 | 4.66E-03 |
| nucleobase-containing compound metabolic process (GO:0006139) | 0.47 | 1.91E-03 |
| cellular nitrogen compound metabolic process (GO:0034641) | 0.45 | 3.07E-05 |
| macromolecule biosynthetic process (GO:0009059) | 0.37 | 4.42E-04 |
| macromolecule metabolic process (GO:0043170) | 0.36 | 1.50E-07 |
| nucleic acid metabolic process (GO:0090304) | 0.31 | 4.38E-04 |
| carbohydrate derivative biosynthetic process (GO:1901137) | 0.2 | 1.13E-02 |
| DNA metabolic process (GO:0006259) | 0.13 | 1.94E-03 |
| amide biosynthetic process (GO:0043604) | 0.1 | 3.54E-02 |
| translation (GO:0006412) | < 0.01 | 4.13E-02 |
| ncRNA metabolic process (GO:0034660) | < 0.01 | 1.39E-02 |
| peptide biosynthetic process (GO:0043043) | < 0.01 | 3.01E-02 |

**Table S16 Transcripts that are more abundant in Δ*tolC tufA::kan* NMM compared to Δ*tolC tufA::kan***

| Gene ontology biological process | Fold Enrichment | p-value (FDR correction) |
|---|---|---|
| heme oxidation (GO:0006788) | > 100 | 1.26E-04 |
| tetrapyrrole catabolic process (GO:0033015) | > 100 | 2.03E-04 |
| porphyrin-containing compound catabolic process (GO:0006787) | > 100 | 1.88E-04 |
| aerobic electron transport chain (GO:0019646) | 67.36 | 1.59E-03 |
| nitrate assimilation (GO:0042128) | 60.45 | 3.91E-06 |
| nitrate metabolic process (GO:0042126) | 60.45 | 3.42E-06 |
| nitrogen cycle metabolic process (GO:0071941) | 52.39 | 6.55E-06 |
| reactive nitrogen species metabolic process (GO:2001057) | 52.39 | 5.89E-06 |
| formate oxidation (GO:0015944) | 49.64 | 3.96E-03 |
| pigment metabolic process (GO:0042440) | 44.91 | 5.14E-03 |
| heme metabolic process (GO:0042168) | 44.91 | 4.88E-03 |
| oxidative phosphorylation (GO:0006119) | 42.87 | 5.37E-03 |
| anaerobic electron transport chain (GO:0019645) | 41 | 1.30E-06 |
| porphyrin-containing compound metabolic process (GO:0006778) | 39.29 | 6.71E-03 |
| formate metabolic process (GO:0015942) | 39.29 | 6.42E-03 |
| respiratory electron transport chain (GO:0022904) | 36.27 | 5.94E-10 |
| tetrapyrrole metabolic process (GO:0033013) | 32.52 | 1.10E-02 |
| anaerobic respiration (GO:0009061) | 28.15 | 7.29E-06 |
| electron transport chain (GO:0022900) | 23.98 | 1.40E-08 |
| cellular respiration (GO:0045333) | 23 | 1.36E-08 |
| energy derivation by oxidation of organic compounds (GO:0015980) | 20.07 | 3.57E-08 |
| generation of precursor metabolites and energy (GO:0006091) | 13.67 | 9.18E-07 |
| oxoacid metabolic process (GO:0043436) | 5.35 | 1.11E-03 |

| | | |
|---|---|---|
| organic acid metabolic process (GO:0006082) | 5.21 | 1.30E-03 |
| small molecule metabolic process (GO:0044281) | 3.19 | 4.65E-02 |
